# Biophysics of microRNA-34a targeting and its influence on down-regulation

**DOI:** 10.1101/2024.02.14.580117

**Authors:** Lara Sweetapple, David M Kosek, Elnaz Banijamali, Walter Becker, Juliane Müller, Christina Karadiakos, Lorenzo Baronti, Ileana Guzzetti, Dimitri Schritt, Alan Chen, Emma R Andersson, Katja Petzold

## Abstract

microRNAs (miRNAs) regulate target mRNA expression post-transcriptionally through their association with Argonaute 2 (AGO2) proteins. Predicting the efficiency of mRNA repression by miRNA has been limited by our comprehension of the structure-function relationship within miRNA binding sites. Using a combination of EMSA, luciferase reporter assays, and structural probing, we investigated the interaction between the human tumour suppressor *miR-34a* and 12 mRNA targets. Comparison of direct RNA:RNA interactions and those within the functional AGO2 protein revealed that the isolated mRNA:miRNA duplex serves as a strong predictor for duplex affinity and structure within AGO2. Our findings reveal that AGO2 has a bidirectional capacity to modulate affinity; weakening tight RNA:RNA binders while strengthening weak ones. We identified three distinct structural groups that form upon *miR-34a* binding and reveal a novel structural group that exhibits a guide strand bulge. MD simulations indicate a conceivable fit of this miRNA-bulge structure within AGO2. Our results demonstrate that the structural characteristics of mRNA:miRNA duplexes could serve as contributing determinants of repression efficacy.

## INTRODUCTION

microRNAs (miRNAs) are short, 19-25 nucleotide long, endogenous RNA molecules that play a key role in post-transcriptional gene regulation. Argonaute (AGO) proteins loaded with miRNAs form the RNA-induced Silencing Complex (RISC). In eukaryotes, miRNA guides RISC to selectively recognise and bind mRNA targets, typically in the 3’ untranslated region (3’UTR), leading to either translational repression or mRNA degradation. With conserved miRNA target sites present in over 60 % of human mRNAs^1^, miRNAs regulate numerous biological pathways and contribute to diverse aspects of cellular function. Despite their importance, and extensive research connecting miRNAs to gene regulation and disease^2^, there is limited progress in understanding their functional mechanisms and application to translational research^3^.

The interaction of a miRNA with an mRNA target is guided by a complementary seed^4^. The seed is located on the miRNA (also called guide RNA) 5’ nucleotides 2-8 (Figure 1), is highly conserved, and is sufficient for downregulation^5,6^. miRNAs are grouped into families sharing identical seeds, and mRNA target sites are classified according to the number of miRNA seed-complementary nucleotides^7^. In order of decreasing efficiency, the 8mer seed has a match between miRNA positions 2-8 with an A in target position 1, the 7mer-m8 has a 2-8 match, and the 7mer-A1 has a 2-7 match with an A in target position 1. Beyond the seed region, imperfect base pairing patterns most commonly occur in mammalian miRNAs^8^. The supplementary region, spanning nucleotides 13-16, can enhance target affinity by over tenfold compared to seed binding alone, and determines target retention by AGO^9^. A recent high-throughput study has proposed that some miRNAs have two high-affinity binding modes; one alternative mode with a bridging loop on the target side between the seed and supplementary regions, and a more conventional mode with zero offset between the guide and target^10^. While binding affinity is regarded as the strongest determinant of miRNA targeting efficacy^11^, the precise impact of binding energy on functional activity within RISC remains incompletely understood.

**Figure 1.**
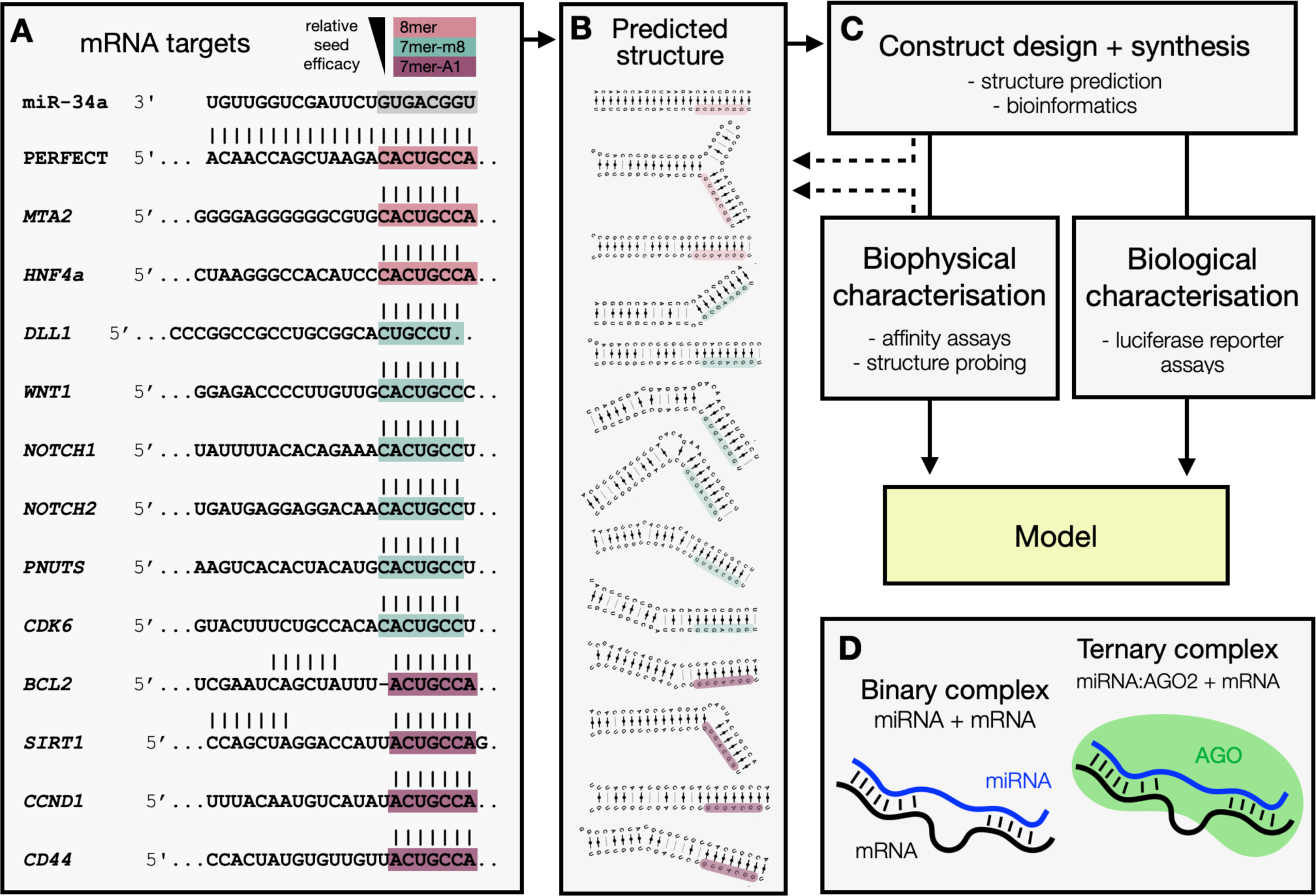
Study design and methodological approach. RNA constructs were designed from twelve biologically and predicted structurally diverse *miR-34a* targets, and were characterised using EMSA, RABS and luciferase assays to assess their biophysical and biological properties upon binding to *miR-34a*. (A) Sequences of *miR-34a* (22 nucleotides long) and the twelve selected *miR-34a* targets with base pairing patterns informed by TargetScan (23-nucleotide long sites). (B) Secondary structures for each mRNA:*miR-34a* duplex predicted by MC-fold, using the chosen construct lengths. (C) Characterisation pipeline following construct design. (D) These properties were assessed within the context of both the binary and ternary complex environments.

Other than target binding strength, gene regulation by miRNA involves an interplay of factors, including sequence composition, conservation, accessibility, miRNA and mRNA target abundance, and competing interactions^12–14^. The miRNA seed and its matched target site nucleotides are conserved, with their probability of conservation correlating with repression, while the conservation of the supplementary region in seed-matched targets is rare^1^. The structural landscape surrounding the miRNA binding site is also of critical importance, as secondary structure can prevent accessibility by RISC^14^. However, efficient pairing to the miRNA 3’ can compensate for low accessibility of the seed^6^. Contributing yet an additional layer of complexity, the effects of the seed and supplementary regions on targeting efficacy appear to be specific to the individual miRNA^11^. While extensive research has focused predominantly on sequence-related aspects of the seed region, a full explanation of functional targeting outcomes remains elusive.

Most miRNA prediction tools utilise common elements: seed identity, sequence conservation, site accessibility, and binding free energy^12^. However, even the most accurate predictive model for targeting efficacy (TargetScanHuman 8.0) is informed by biochemical analyses of 12mers^11^, which neglects the effect of potential secondary structure within the miRNA target site interaction due to the short length of RNAs assessed. Notably, TargetScanHuman 8.0 accounts for approximately half of the observed variability in mRNA repression, leaving the remaining half unexplained^11^. This suggests the existence of undiscovered factors influencing targeting efficacy that are yet to be incorporated into predictive models.

Crystallography has revealed several important insights into the miRNA targeting mechanism. Upon initial pairing to a seed-matched target, human AGO2 undergoes a conformational shift in helix-7, enabling subsequent pairing in the supplementary region (nucleotides 13–16) of the mRNA:miRNA duplex^16^. In addition to base pairing interaction, sequence-independent recognition occurs along the minor groove, highlighting the importance of shape complementarity. MacRae and colleagues demonstrated that AGO2 avoids base pairing within the miRNA central region (nucleotides 9–11) and can accommodate up to five contiguous base pairs in AGO2’s supplementary chamber^17^. Between the seed and supplementary regions, a bridge of up to 15 unpaired nucleotides can occur on the target side^10,17,18^, and both hairpins and unstructured bulges are tolerated in this bridge^6^. A recent crystallographic study^19^ revealed that certain miRNA targets exhibiting extended miRNA 3’ pairing with central mismatches adopt a unique conformation in which the miRNA 3’ tail is exposed and leads to its degradation, termed Target-Directed miRNA Degradation (TDMD)^20,21^. A similar but less-explored phenomenon has been observed whereby highly complementary targets with extensive 3’ tail pairing induce mRNA:miRNA duplex release from AGO2, although the function and mechanism remain unclear^22,23^.

The microRNA *miR-34a* belongs to the conserved miR-34/449 family, which regulates cell cycle regulation and is controlled by p53. In human cells, *miR-34a* acts as a tumour suppressor, regulating hundreds of mRNA targets, with implications in cancer, cardiovascular and neurological disease^24,25^. Recently, an NMR-derived structural model of *miR-34a* binding to its mRNA target, *SIRT1,* was solved, revealing a bent structure with an asymmetric 4 nucleotide bulge on the target side^26^. Relaxation dispersion experiments^27,28^ demonstrated a dynamic shift from the 7mer-A1 ground state to a transient extended 8mer seed. Molecular dynamics simulations indicated that the excited state altered the angle between the seed and supplementary regions, leading to release of the miRNA 3’ end, and stabilising this excited state resulted in enhanced repression of *SIRT1*. It was found that this dynamic seed extension can occur in 18% of all *miR-34a* targets^26^. These findings underscore the significance of structural dynamics in mRNA:miRNA interaction and raise questions about the broader implications of such structural features in the understanding of miRNA activity.

Here we employ electromobility shift assay (EMSA), structural probing (RABS^29^), and luciferase gene reporter assays to study the structure-function relationship involved in microRNA targeting, focusing on human *miR-34a* and 12 of its mRNA targets: *BCL2*, *CD44*, *CDK6*, *CCND1*, *DLL1*, *HNF4α*, *MTA2*, *NOTCH1*, *NOTCH2*, *PNUTS*, *SIRT1*, and *WNT1*. We find that mRNA:*miR-34a*-AGO2 binding affinity correlates with mRNA:*miR- 34a* affinity, and that AGO2 acts to constrict the range of affinities observed in RNA:RNA binding alone; strengthening weak binders while weakening strong binders. Similarly, we find that the structure within the *miR-34a*-AGO2 complex is determined by the underlying RNA:RNA interaction, resulting in three distinct modes of binding: miRNA-bulged, mRNA- bulged, and symmetrical. Each of these three groups displays distinct correlations between its affinity and degree of down-regulation. We show via 3D structure simulation that the novel miRNA-bulge mode is plausible within AGO2. We propose that these modes contribute towards determining the mRNA target repression outcome and should be considered in microRNA targeting prediction algorithms.

## MATERIALS AND METHODS

### Target pool selection and construct design

The *miR-34a* target pool was designed to represent a diversity in biological function, sequence, seed, and predicted mRNA:*miR-34a* structure. We reasoned that *miR-34a* can bind to stretches of mRNA target shorter or longer than the conventional 23 nucleotide binding site reported by TargetScan if unpaired nucleotides can occur on either side of the interaction between stronger binding areas. This hypothesis is substantiated by previous findings demonstrating that the mRNA target *SIRT1* interacts with *miR-34a* through a 26-nucleotide long binding site^26^. Consequently, we expanded our examination of binding sites to include sequence lengths of up to 35 nucleotides for the mRNA target site; starting at the mRNA predicted seed (3’ end) and extending in the 5’ direction.

Thirty-seven previously validated^30,31^ direct *miR-34a* targets were assessed, and potential secondary structures were predicted using MC-fold^32^ for each mRNA:*miR-34a* pair. We compared the 50 lowest energy secondary structures between the 35-nucleotide target sequence with successively shorter iterations, down to an 18-nucleotide binding site. We chose the shortest target length in which overall structural features were preserved, and the interaction with *miR-34a* was maximised, whilst controlling for self-binding tails that might perturb the structure of the binding site. All candidate RNA pairs were characterised in terms of sequence, predicted secondary structure, and biological features, and were then visualised using Principal Component Analysis (PCA) followed by K-means clustering (Supplementary Figure S1). A total of 12 targets were then selected for further study. Extraction of secondary structures features was carried out using the Forgi package^33^. Analysis code is provided in Supplementary File 1.

### Denaturing RNA polyacrylamide gel electrophoresis (PAGE)

Quality assessment of all synthesised RNA samples was conducted using denaturing polyacrylamide gel electrophoresis in a Mini-PROTEAN® Tetra System (Bio-Rad). Gels of 0.75 mm thickness were cast using approximately 5 mL PAGE mix (20% acrylamide-bis 19:1), 8 M urea in 1X TBE (100 mM Tris, 90 mM boric acid, 1 mM EDTA), and polymerisation was initiated with 40 µL 10% APS and 4 µL TEMED. Gels were pre-heated in 1X TBE at 350 V for 10 minutes, and samples denatured in loading buffer (240 µM bromophenol blue in formamide, 5 mM EDTA) at 95 °C for 3 minutes. The denatured solutions were loaded into pre-rinsed wells and run in 1X TBE buffer at 350 V for 1 hour, allowing for single-nucleotide resolution length separation within the 20 – 35 nucleotide range. The gels were then stained using SYBR Gold according to the manufacturer’s protocol (Invitrogen) and imaged with a GE Healthcare ImageQuant LAS 4000.

### T7 in vitro transcription

Unlabelled *miR-34a* was synthesised via *T7 in vitro* transcription using tandem repeats on a plasmid template^34^. In brief, a plasmid encoding 26X repeats of the *miR-34a* transcript (Genscript) was transcribed via T7 RNA polymerase (T7 RNAP; produced by Protein Science Facility at Karolinska Institute) in a reaction volume of 10 mL and annealed to a chimeric cleavage guide (ordered from Integrated DNA technologies, IDT). The RNA:DNA hybrid was then cleaved by RNase H (NEB catalogue # M0297L) to produce the final 22 nucleotide product. The desired RNA was separated from additional cleavage products via ion exchange HPLC (see below). For plasmid and cleavage guide sequence details see Feyrer et al. 2020^34^. RNA scaffolds for RABS experiments^29^ were transcribed in 100 µL volumes containing 100 mM Tris–HCl pH 8.0, 10 mM MgCl_2_, 10 mM dithiothreitol, 20 mM spermidine, 5 mM GMP, 3 mM NTPs, 0.3 mg/ml T7 polymerase, 0.1 mg/mL inorganic pyrophosphatase (produced by Protein Science Facility at Karolinska Institute), and 3.6 ng/mL of dsDNA template.

### Solid-phase RNA synthesis

The remaining oligonucleotides were synthesised using an H-8 DNA/RNA synthesiser (K&A Labs GmbH). Syntheses were conducted DMT-ON using a 1 µM scale and RNA amidites Bz-A-CE, Ac-C-CE, iBu-G-CE, U-CE (Sigma-Aldrich) dissolved in acetonitrile at 70 mM, with 12-minute coupling times. For cleavage of oligonucleotides from the solid support following synthesis, samples were incubated in 1 mL ammonium hydroxide methylamine (AMA; a 1:1 solution of 28% ammonium hydroxide and 40% aqueous methylamine) for 30 minutes at room temperature. An additional 0.5 mL AMA was added to elute cleaved product from the support, and the eluate was then incubated for 30 min at 65 °C. Excess AMA was then evaporated from the oligonucleotides by drying the samples for two hours at low speed without heating using a vacuum concentrator (Savant SpeedVac, ThermoScientific), followed by overnight lyophilisation. Pellets were dissolved in 115 µL anhydrous DMSO followed by 60 µL triethylamine (TEA). To remove the 2’hydroxyl protecting group TBDMS, 75 µL TEA.3HF was added and heated up to 65 °C for 2.5 h. Quenching was conducted with 1.75 mL 1 M TRIS, pH 8.0. To purify the full-length oligonucleotides from shorter oligonucleotides a DMT-ON purification with Glen-Pak RNA purification cartridges (#60-6200) was conducted. The cartridges were prepared before sample loading with 0.5 mL acetonitrile and 1 mL 2 M triethylammonium acetate (TEAA). The oligonucleotides were first washed with 1 mL of a mixture of 10% acetonitrile and 90% 2 M TEAA, and secondly with 1 mL H_2_O. For detritylation 2 mL of 2% trifluoroacetic acid (TFA) were added. After a second wash with 2 mL H_2_O, samples were eluted in 1 mL elution buffer (1 M NH_4_HCO_3_, 30% ACN). The elution fraction was dried again with a speed vac, lyophilised overnight, and diluted in water. All washes and elution were evaluated on a 20% denaturing PAGE gel.

### Ion-exchange HPLC purification

*In vitro* transcription and solid phase synthesis products were purified via anion-exchange HPLC. Samples were loaded on a DNAPac PA200 column (22×250 mm; Thermo Fisher, Thermo Fisher Ultimate 3000 system) at 75 °C under low salt conditions in buffer A (20 mm NaOAc pH 6.5, 20 mm NaClO4, 10% MeCN, *pH 6.5*), and eluted using an increasing sodium perchlorate concentration via buffer B (20 mm NaOAc pH 6.5, 600 mm NaClO4, 10% MeCN, *pH 6.5*). The gradient was optimised for each construct to achieve optimal separation. For *miR-34a*, a gradient of 20–26% buffer B in 10 column volumes at a flow rate of 8 mL/min was used. Purity of the HPLC fractions was monitored by absorbance at 260 nm and subsequently assessed by denaturing PAGE as described above. Fractions containing sufficiently pure RNA were pooled and desalted in water using Amicon Ultra centrifugal filters with a 3 kDa MWCO.

### Cloning and recombinant baculovirus production for AGO2 production

The *miR-34a*-AGO2 complex was prepared as reported by Banijamali et al. (2022)^29^. Briefly, the full-length human AGO2 was cloned into a pFastBac dual expression vector (Thermo Fisher) containing an N-terminal His-tag followed by a TEV cleavage site and transformed into chemically competent DH10EmBacY cells (Geneva Biotech). Blue/white screening was performed to confirm the presence of the recombinant Bacmid, followed by bacmid isolation and PCR analysis, according to the guidelines provided in the “Bac-to-Bac TOPO Expression System” user manual (Thermo Fisher, Version A, 15 December 2008 A10606). Fresh Sf9 cells were transfected with the recombinant AGO2 Bacmid. The transfection was carried out in 8×10^5^ Sf9 cells/plate (1.5×10^6^ cells/mL) and amplified through two subsequent rounds of transfection. Monitoring of baculovirus production was facilitated by internal GFP expression. 300 mL of the final baculovirus stock was filtered, and FBS was added to a final concentration of 2%. Successful overexpression of AGO2 was confirmed using Western Blot analysis with an anti-6X His tag antibody (Abcam, AB18184).

### AGO2 expression

10 mL AGO2 baculovirus stock was used to transfect each litre of Sf9 cells (1.5×106 cells/mL, passage 22, grown in Sf-900 II SFM media; Thermo Fisher). In total, 6 L were incubated for 72 hours at 27°C using an orbital shaker. The expression was monitored by observing the Bacmid internal GFP expression under polyhedrin promoter control. The cells were then centrifuged, and the resulting pellet (∼45g) was washed with PBS and resuspended in IMAC buffer A (50 mM Tris-HCl, 300 mM NaCl, 10 mM Imidazole, 1 mM TCEP, and 5% glycerol (v/v)) with 25x EDTA-free protease inhibitor cocktail (Merck). The cells were lysed using three cycles of freeze/thawing on dry ice, followed by sonication for 10 minutes at 30% amplitude with 10-second ON and 10-second OFF cycles to reduce the lysate viscosity. The lysate was centrifuged at 50,000 RCF for 1 hour at 4°C and the resulting supernatant was then filtered through a 0.22 µm sterile filter before being applied onto an IMAC buffer A-equilibrated HisTrap-Ni^2+^ column (Cytiva; HisTrap HP). A linear gradient was employed using buffer A (50 mM Tris-HCl, 300 mM NaCl, 10 mM Imidazole, 1 mM TCEP, 5% glycerol v/v) and buffer B (50 mM Tris-HCl, 300 mM NaCl, 300 mM Imidazole, 1 mM TCEP, 5% glycerol). All fractions containing the desired protein were combined and concentrated from 15 mL to 5 mL using a 30 kDa cut-off Amicon centrifugal filter unit (Sigma Aldrich).

### Loading of AGO2 with miR-34a

The AGO2 stock was incubated with crude *in vitro* transcribed *miR-34a* (3 mL of reaction concentrated to 600 μL) in a water bath at 37°C for 5 hours to facilitate guide RNA loading. The loaded protein solution was dialysed (Spectrum, 3000 MWCO) and 200 μL of TEV protease added (produced by Protein Science Facility at Karolinska Institute). The mixture was further dialysed against 2 L of IMAC buffer A overnight at 8°C. Following dialysis, any precipitate was removed by centrifugation, and the resulting supernatant was loaded onto a pre-equilibrated HisTrap-Ni^2+^ column to remove TEV protease and other impurities. The flow-through (5 mL) was collected and applied onto a pre-equilibrated size exclusion chromatography (SEC) column (Cytiva, HiLoad 16/600 Superdex 200 pg) using buffer C (20 mM HEPES, 100 mM KCl, 1 mM TCEP, 5% glycerol) (Figure S2A). All fractions from both the HisTrap-Ni^2+^ and SEC columns were analysed using SDS-PAGE (Invitrogen, NuPAGE 4 to 12%, Bis-Tris, 1.0 mm) to assess purity (Figure S2B). Protein concentration was determined using a Bradford assay (Thermo Fisher PierceTM Detergent Compatible Bradford Assay Kit) using BSA as a standard (Figure S2C). Absorbance measurements were performed at 595 nm using a Varioskan LUX multimode microplate reader (Thermo Fisher). The loading efficiency was estimated using a northern blot protocol optimised for detection of small RNAs^35^, with a 3’Cy3 labelled *miR-34a* probe for detection (Figure S2D).

### Slicing assay for AGO2 activity

To ascertain whether the loaded AGO2 sample was active, we performed a slicing assay on 3’-Cy3 labelled 34 nucleotide probe that was perfectly complementary to *miR-34a*, with a 6 nucleotide 5’ overhang and a 5 nucleotide 3’ overhang (AGO2 slicing probe; Table S1). Following incubation of *miR-34a* loaded AGO2 with the probe in slicing buffer^16^ (20 mM Tris-HCl pH 8.0, 150 mM NaCl, 2 mM MgCl_2_, 0.5 mM TCEP, 0.01 mg/ml baker’s yeast tRNA, 5% (v/v) glycerol), the formation of a fluorescently labelled 15 nucleotide cleaved product was confirmed by denaturing PAGE (Figure S2E).

### Binary binding assays (mRNA:miRNA)

To determine the apparent equilibrium dissociation constant (*K*_D,app_) of each mRNA:*miR- 34a* duplex, we adapted an EMSA protocol^36^ using fluorescent probes to monitor RNA:RNA binding. A 3’Cy5-labelled *miR-34a* (ordered from IDT; RNase-free HPLC purity) was titrated against a 1000-fold range of each mRNA target in EMSA buffer (15 mM NaH_2_PO_4_/Na_2_HPO_4_ pH 6.5, 25 mM NaCl, 2 mM MgCl_2_, 0.1 mM EDTA). The amount of Cy5-miR34a was fixed to 10 nM to give sufficient signal during detection. To promote bimolecular interaction during annealing, samples were heated to 95 °C, cooled slowly to 30 °C for 35 minutes, and incubated at 22 °C for 85 minutes. 10 µl of each sample was mixed with 4 µL native loading dye (1% Orange G (NEB) dissolved in 50% glycerol and 50% EMSA buffer) and run on a non-denaturing polyacrylamide gel (10% acrylamide-bis 19:1 in EMSA buffer) in 1X EMSA buffer at constant voltage (150 V for 40 min) using a Mini-PROTEAN® Tetra System (Bio-Rad). To prevent heating during electrophoresis, two 50 ml PEG cooling blocks were added to the electrophoresis chamber and the temperature was monitored to remain between 22-23 °C. In addition, the electrophoresis chamber was protected from light to prevent bleaching of Cy5. Each gel was imaged twice; first for Cy5 fluorescence detection, and subsequently following SYBR Gold staining (Invitrogen), using an ImageQuant LAS-4000 imager (GE healthcare). Fluorescence signals of the free and bound forms of Cy5-miR34a were quantified using Image Lab software 6.0.1 (Bio-Rad). To assess any potential impact of the fluorophore on the observed binding affinities, we conducted a reverse binding experiment. In this case, the fluorophore was placed on the target (*CD44*), while *miR-34a* remained unlabelled. We observed the same *K*_D,app_ within error for *CD44* (Figure S3B), providing evidence that the fluorophore itself did not significantly interfere with the binding process for this control target. Each experiment was repeated in triplicate and fitted to a sigmoidal binding curve (*Y* = *b* + *X* ∗ (*m* − *b*)/(*KD* + *X*)), where b = the baseline signal in the absence of binding and m = the maximum signal once the binding plateau is reached, using GraphPad Prism software version 9.0. Representative gel images are shown in Figure S3A and all data are presented in Supplementary File 2.

### Ternary binding assays (mRNA:miRNA-AGO2)

We adapted the RNA:RNA binding EMSA protocol described above to measure equilibrium dissociation constants for each mRNA:*miR-34a* interaction in the presence of AGO2. *miR-34a* was pre-loaded into AGO2 as described above and equilibrated with 5’ modified Cy5 target mRNAs (ordered from IDT; RNase-free HPLC purity) in EMSA buffer supplemented with 20 µM polyuridine (pU). Reactions were equilibrated at 37 °C in the dark for 1 hour, using a total reaction volume of 10 µL. Target RNAs were kept at a constant concentration of 10 nM, and were titrated against a 1000-fold range of *miR34a*- AGO. Following equilibration, samples were mixed with 4 µL native loading dye and electrophoresed on a non-denaturing polyacrylamide gel as described above. The entire experiment was run in triplicates for each mRNA target. Representative gel images are shown in Figure S4 and all data are presented in Supplementary File 3. Fluorescence of the bound and free forms was quantified according to the method described in Figure S5 to estimate the *K*_D,app_ for each interaction.

### Dual Luciferase reporter assays - cell culture and plasmid cloning

Reporter assays were carried out according to the protocol described by Kosek et al. (2023)^6^. HEK293T cells (purchased from ATCC) were cultured in Dulbecco’s modified essential medium (DMEM, Gibco) supplemented with 10% fetal bovine serum (FBS, Gibco). The cells were tested for mycoplasma contamination (Mycoplasmacheck, Eurofins) and confirmed to be mycoplasma-free. Target sites were cloned into a psiCHECK2 vector (Addgene plasmid #78258) between the XhoI and NotI restriction sites. The plasmid was cleaved by XhoI (NEB, #R0146) and NotI (NEB, #R3189) in CutSmart buffer (NEB, #B7204) with calf intestinal phosphatase (NEB, #M0290). Cleavage products were separated on a 1% agarose gel and the linearised plasmid was purified with a QIAquick Gel Extraction Kit (Qiagen, #28706). Complementary synthetic DNA oligos coding for *miR-34a* target sites (Table S1), with restriction site overhangs for XhoI and NotI, were purchased from IDT. Inserts were phosphorylated by incubation with T4 polynucleotide kinase (NEB, #M0201) in T4 DNA ligase buffer (#, NEB) at 37°C for 30 minutes, and then annealed by heating to 95°C for 5 minutes followed by incubation at room temperature. Annealed inserts were ligated into the vector with T4 DNA ligase (NEB, #0202) in T4 DNA ligase buffer for 20 minutes at room temperature. Ligated vectors were transformed into competent bacteria (One Shot TOP10 *E.* coli; Invitrogen) and grown overnight on ampicillin plates. Selected colonies were grown overnight in LB medium, and plasmids were purified using a PureLink Plasmid Miniprep Kit (Invitrogen, #K210011) or a Qiagen Plasmid Midi Kit (Qiagen, #12145). Correct insertions were verified by Sanger sequencing (Lightrunc, Eurofins).

### Dual Luciferase reporter assays - oligo preparation, transfection, and measurement

Synthetic *miR-34a* guide and passenger strands were purchased from IDT (PAGE purity), resuspended in Nuclease-Free Duplex Buffer (30 mM HEPES, pH 7.5; 100 mM potassium acetate; IDT, #11-01-03-01) and annealed by heating to 95°C for 2 minutes followed by incubation at room temperature.

HEK293T cells were seeded in 48-well plates 24 h prior to transfection. At 80-90% confluency, cells were transfected with 0.4 µg plasmid and either 0 pmol or 9 pmol *miR- 34a* duplex (to a final concentration of 0 nM or 30 nM miRNA), using Lipofectamine 2000 (Invitrogen) in Opti-MEM (Gibco) according to the manufacturer’s protocol. After 24 h incubation, cells were washed once with 0.1 mL phosphate-buffered saline (PBS) and luciferase activity was measured with Dual Luciferase Reporter Assay System (Promega, E1910) according to the manufacturer’s protocol. Measurements were made on a Promega GloMax 96 luminometer with 1 s delay and 10 s integration time. For each sample, the signal from *Renilla* luciferase (carrying the miRNA target site) was normalised to the signal from *Firefly* luciferase. The mean relative expression (expressed in terms of *Renilla* to *Firefly* luciferase - R/F - ratio) was obtained from three separate biological replicates in HEK293T cells. For each target site, the Renilla/Firefly Luciferase (R/F) ratio from the 30 nM *miR-34a* sample was then normalised to the 0 nM *miR-34a* sample. All raw and processed luciferase data are provided in Supplementary File 4.

To ascertain that the surrounding mRNA sequence did not influence repression efficacy, we assessed the predicted structural accessibility of each target site within the full length *Renilla* luciferase mRNA using RNApIfold^37^. This tool predicts the probability of a given region being unpaired, and results are outlined in Figure S8C.

### RNA:RNA binding by RABS

The RABS method was used to assess RNA:RNA and RNA:RNA-AGO2 interaction and structure, as previously described by Banijamali et al. (2022)^29^. Two RNA scaffolds were used for each mRNA target; i) a CIS-scaffold: RNA scaffold containing both mRNA target and miRNA sequence separated by a non-binding closing loop, and ii) a TRANS-scaffold: RNA scaffold containing the mRNA target sequence, to which free *miR-34a* or *miR-34a*- AGO2 could be bound (Figure 4A). The overall conformation of each scaffold was assessed using the RNAstructure package^38^ to ensure that the sequence of interest did not interact with the scaffold. Full scaffold sequence details are shown in supplementary information (Table S1). DNA templates containing a T7 polymerase promotor were generated via PCR assembly^39^ (using primers purchased at IDT) and the scaffolds were transcribed via *in vitro* transcription as described above. RNA products purified with RNAClean XP beads (Beckman Coulter).

The RNA scaffolds were folded in Na-HEPES buffer (pH 8): each RNA was heated to 95°C, snap-cooled on ice for 30 minutes, and incubated at room temperature for 30 minutes. For TRANS experiments, *miR-34a* or *miR-34a:*AGO2 was added at 1:2 and 1:1 ratios of mRNA:*miR-34a*(AGO2) respectively, after folding and prior to room temperature incubation of mRNA.

RNA was separated into X8 15 μL reactions (80 nM RNA, 67 mM Na- HEPES, 10 mM MgCl_2_) for treatment with the modifier 1-methyl-7-nitroisatoic anhydride (1M7^40^; Sigma- Aldrich). 1.72 μL of 1M7 (100 μM stock) or ddH_2_O was added per (X3) modified or (X5) unmodified reaction respectively. RNA was recovered using 9.8 μL recovery mix (0.25 M Na-MES (pH 6), 1.5 M NaCl, 0.006 μM 5-poly-dA-FAM- labeled primers (IDT), 1.5 μL poly-dT Dynabeads (Thermo Fisher)) per reaction. For *miR-34a:*AGO2-containing reactions, 45 μL of proteinase K (PK) mix (20 μg PK (Sigma Aldrich, PCR grade), 50 mM Tris-HCl pH 7.5, 75 mM NaCl, 6.25 mM EDTA, 1% SDS (w/v)) was added per reaction and incubated at 65 °C for 1 hour. After inactivating PK (120 μL 70% ethanol, followed by 5 minutes on ice), recovery mix (19.6 μL) was added to each reaction.

Of the five unmodified reactions, four were used as Sanger sequencing ladders, and one as an unmodified control. The three modified reactions served as technical replicates. The recovered RNA was then washed in 70 % ethanol and air dried, to which 2.5 μL of ddH_2_O and 2.5 μL of reverse transcription (RT) mix (1 μL 5X First strand buffer (Thermo Fisher), 0.01 M DTT, 1.6 mM dNTPs, 0.1 μL SuperScript III Reverse Transcriptase (Thermo Fisher)) were added. For the Sanger sequencing ladder reactions RT mixes were prepared using a 1:6 ratio of dNTPs:ddNTP. All reactions were incubated at 50 °C for 30 minutes. RNA was then hydrolysed using 5 μL NaOH (0.4 M) at 90 °C for 3 minutes, then cooled on ice for 3 minutes. To neutralise, 5 μL of Acid mix (1 volume of 1.25 M NaCl, 1 volume of 0.5 M HCl, 2 volumes of 1 M NaOAc) was added per reaction. Samples were then washed 3 times with 70% ethanol and air-dried. The cDNA was eluted from Dynabeads using 11 μL Formamide-ROX mix (1000 μl Hi-Di Formamide (Thermo Fisher), 8 μl of 350 ROX size standard (Thermo Fisher)) per reaction and incubating at room temperature for 15 minutes. All samples were prepared in two different dilutions: saturated (1:2 dilution of eluted cDNA with Formamide-ROX mix) and diluted (1:15 dilution of eluted cDNA with Formamide-ROX mix). Sites of modification (acyl adducts) were detected by primer extension and resolved by capillary electrophoresis via KIGene core facility at Karolinska Institute.

### RABS data processing

RABS data were aligned, corrected for signal decay, integrated, and normalised according to the HiTrace pipeline (High-Throughput Robust Analysis for Capillary Electrophoresis)^41^. To compare differing TRANS experimental conditions (free, +*miR-34a*, +*miR-34a*-AGO2) reactivity values of the nucleotides in the main loop were normalised to the averaged reactivity values of the surrounding buffer nucleotides (Table S1). Consistent with previous work^6,29,42^, reactivity values were limited to 0 – 1 to eliminate negative averaging artefacts and large positive values representing dynamic variation. Nucleotide positions with values greater than 0.5 were classified as unpaired or in flexible conformations while values ≤ 0.5 were classified as base paired or in inaccessible conformations. Each condition was tested with at least two biological replicates, each containing three technical replicates. Raw and processed data is shown in Supplementary File 5. To assess patterns in the reactivity data, we applied both KMeans and hierarchical clustering to the data (using one minus cosine similarity as a distance metric, linked by mean average,) and visualised the data in heatmaps via Morpheus by Broad Institute (RRID:SCR_017386).

### Secondary structure prediction

The CIS and TRANS reactivity data were then used to predict the secondary structures of each mRNA:*miR-34a* duplex. To achieve this, we employed the ViennaRNA package RNAcofold^37,43^. We explored several folding algorithms^42,44^, including those developed by Washietl et al. and Zarringhalam et al. and adjusted parameters, including the weighting and allowance of non-WC base pairs, to evaluate a wide range of possible structures based on the CIS data (comprising reactivity information from both the *miR-34a* and mRNA sides). All possible secondary structures are listed in Supplementary File 6. Subsequently, using the ViennaRNA Forna package^45^, we superimposed these structural predictions with TRANS + *miR34a*-AGO reactivity data (comprising reactivity data from the target side only). This combined approach allowed us to identify and select the most accurate predicted secondary structure for further analysis.

### Molecular Dynamics (MD) simulations

The *NOTCH1*:*miR-34a*-AGO2 ternary complex was built starting from the published crystal structure of target:*miR-122*-AGO2 (6N4O)^17^ using the Molecular Operating Environment (2022.02 Chemical Computing Group ULC). All existing RNA was removed except the seed region (near MID and L2 domains) which was mutated to that of *miR-34a* nucleotides 2-8. The remaining *miR-34a* and *NOTCH1* RNA (excluding the 11-nucleotide unbound *NOTCH1* 5’ overhang) was appended to the seed with secondary structure as depicted in Figures 5 and S9. The resulting structure was then used to create all-atom simulations in GROMACS 2019.4^46^ using the Amber99-Chen-Garcia RNA force field^47^ with backbone phosphate modifications^48^, and the TIP 4-point water model^49^ with Ewald optimisations and LJ parameters for alkali and halide ions^50^. The systems were solvated with ∼194,000 water molecules in a 10 nm cubic box, with salt conditions of 0.1 M KCl using 366 K^+^ and 364 Cl^-^ ions. The systems were energy-minimised using the steepest descent algorithm with a force tolerance of 1000 kJ mol^-1^nm^-1^. After energy minimisation, equilibration through N, V, T all-atom simulations in Gromacs 2019.4^46^ using the Amber99- Chen-Garcia RNA force field^47^ with backbone phosphate modifications^48^, and the TIP 4- point water model^49^ with Ewald optimisations and LJ parameters for alkali and halide ions^50^. The systems were solvated with ∼194,000 water molecules in a 10 nm cubic box, with salt conditions of 0.1 M KCl using 366 K^+^ and 364 Cl^-^ ions. The systems were energy minimised using the steepest descent algorithm and a force tolerance of 1000 kJ mol^-1^ nm^-1^. After energy minimisation, equilibration through N, V, T, and then N, P, T ensemble using the leap-frog integrator with a time step of 2 fs for 100 ps was applied. The LINCS algorithm^51^ was used to constrain bonds involving H atoms and the modified Berendsen thermostat^52^ was used for temperature coupling. The Verlet scheme was used for buffered neighbor searching with a cutoff of 1 nm. For N, V, T, a random seed was used for velocity generation with velocities assigned from a Maxwell distribution. For N, P, T, an isotropic Parrinello-Rahman barostat^53^ was used for pressure coupling with a time constant of 2.0 ps. The resulting system was then simulated at 350K for 10 ns in three, independent production trajectories, each with randomised starting velocities. This ensured that the complexes formed clash-free and remained stable, maintaining all RNA:protein seed region contacts throughout the course of all simulations (Figure S11). Simulations were visualised using Mol* Viewer^54^. The simulation trajectory file used for image generation is provided in Supplementary File 7.

## RESULTS

The sequence, structure, and biological features of 37 validated miR-34a targets were annotated and visualised via Principle Component Analysis (PCA), and targets belonging to each of the four identified clusters were selected (Supplementary Figure S1). Twelve mRNA targets of *miR-34a* were thus chosen to represent a diversity in biological function, sequence, and predicted structure (Figure 1A). Our determined mRNA target sequences varied in length from 20 – 35 nucleotides (Table S1), and exhibited diverse predicted secondary structures, varying from 12 – 21 predicted base pairs, and bulges ranging from 1 – 12 nucleotides in size (Figure 1B).

Samples of each of the 12 mRNA targets, as well as *miR-34a* and AGO2, were synthesised in-house for biophysical and biological characterisation. Binding affinity of the 12 targets for *miR-34a* was tested via electrophoretic mobility shift assay (EMSA), and a modified EMSA protocol was developed to test their affinity for *miR34a*-AGO2 (the ternary complex). The same 12 targets were then prepared for activity analyses by luciferase gene reporter assays, and for secondary structure analyses via structure probing (RNA:RNA binding by SHAPE (RABS))^29^. Secondary structures were calculated from the RABS data and structural features were compared with respect to their biochemical properties and biological activity to define distinct target classes.

### AGO2 constricts the range of affinities observed in RNA:RNA binding alone

To measure the affinity of each mRNA target binding to *miR-34a*, both within the binary complex (mRNA:*miR34a*) and the ternary complex (mRNA:*miR-34a*-AGO2), we optimised an RNA:RNA binding EMSA protocol to suit small RNA interactions. The protocol is loosely based on Bak et al. (2014)^36^, with major differences being use of a sodium phosphate buffering system supplemented with Mg^2+^ so as not to disturb weaker interactions, and fluorescently labelled probes. Original gel images and quantification are shown in supplementary Figures S3 and S4. All *K*_D,app_ values are shown in Supplementary Table 1, and represent the mean of three independent replicates.

For the binary complex measurements, unlabelled target RNAs were titrated against a constant concentration of Cy3-labelled *miR-34a*. Observed mRNA target affinities spanned four orders of magnitude; from low nanomolar (*SIRT1 K*_D,app_ = 1.27 ± 0.3 nM) to approximately 10 micromolar (*NOTCH2 K*_D,app_ = 6055 ± 3362 nM) (Figure 2A). To assess whether the target affinities were modulated upon binding within the AGO2 protein (or ternary complex), we adapted our assay to enable *miR-34a*-loaded AGO2 titration against a constant concentration of Cy5-labelled target mRNAs, and quantified the disappearance of free mRNA probe as shown in Figure S5A. Again, *SIRT1* was the tightest binder (*K*_D,app_ = 2.68 ± 0.3 nM) and *NOTCH2* the weakest (*K*_D,app_ = 496 ± 129 nM) (Figure 2), though affinity spanned only three orders of magnitude within the ternary dataset. The change in *K*_D,app_ between the binary and ternary complexes was within the margin of error for some targets (e.g., *HNF4α*: 36.7 vs. 38.9 nM and *BCL2*: 10.8 vs. 8.92 nM), but in other instances, such as *MTA2*, it differed by an order of magnitude (944.3 vs. 66.1 nM; Figure 3B). Interestingly, *MTA2* exhibited two *miR-34a*-bound bands due to alternative conformations (details see Figure S3A). For tight binders such as *SIRT1*, the affinity was considerably weakened (0.41 vs. 2.7 nM; Figure 3B). As such, AGO2 exerted a bi- directional effect on RNA:RNA affinity, strengthening the weak binders while weakening the tight binders.

**Figure 2.**
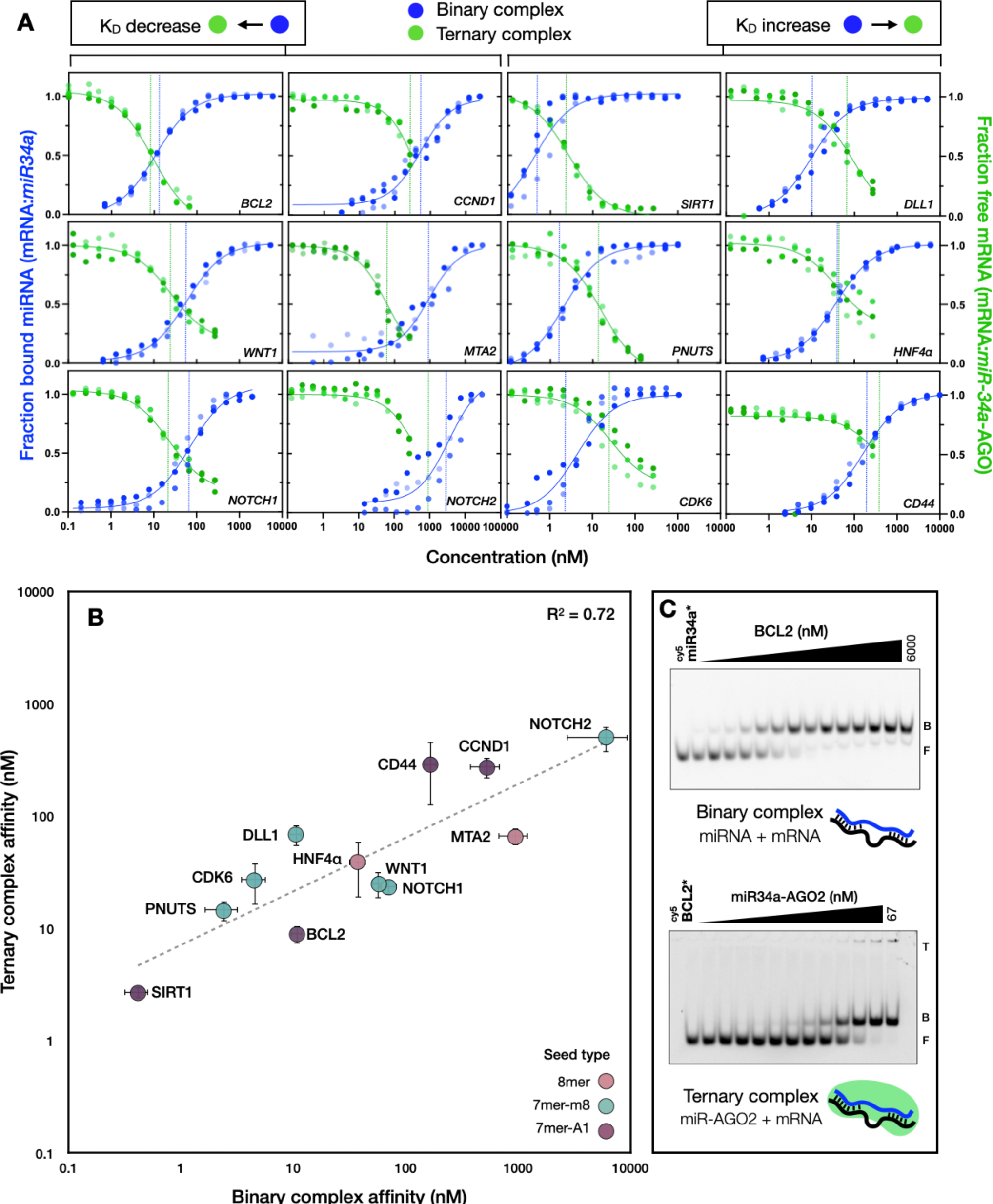
Ternary complex binding affinity can be predicted by binary complex binding. (A) Individual binding profiles for each of the 12 mRNA targets assessed by electrophoretic mobility assay (EMSA). Blue represents results for the binary complex, mRNA:*miR-34a*. Green represents results for the ternary complex, mRNA:*miR-34a*-AGO2. Dotted horizontal lines represent the *K*_D,app_ values. Note that the x-axis spans from 0.1 to 100,000 in *CCND1*, *MTA2* and *NOTCH2*, whereas the remaining targets span 0.1 to 10,000. A full list of *K*_D,app_ can be found in Supplementary Table S2. (B) Correlation of relative mRNA:*miR-34a* with mRNA:*miR-34a-* AGO2 binding affinities. No seed type correlation is observed, seeds coloured, where 8mer is pink, 7mer-m8 is turquoise, and 7-mer-A1 is mauve. The slope of the linear fit is 0.48, and intercept on the (log *y*)-axis is 7.11 (C) Example of *BCL2* mRNA binding assays for both binary (mRNA + miRNA) and ternary (mRNA + *miR-34a-* AGO2) complexes. The labelled species is indicated with (*). F indicates free labelled species (*miR34a* or mRNA), B indicates binary complex and T indicates ternary complex. Adjacent titrations points differ two-fold in concentration, with maximum concentrations 6000 nM *BCL2* or 67 nM *miR-34a*-AGO2. For the ternary complex duplex release from AGO2 was observed.

**Figure 3.**
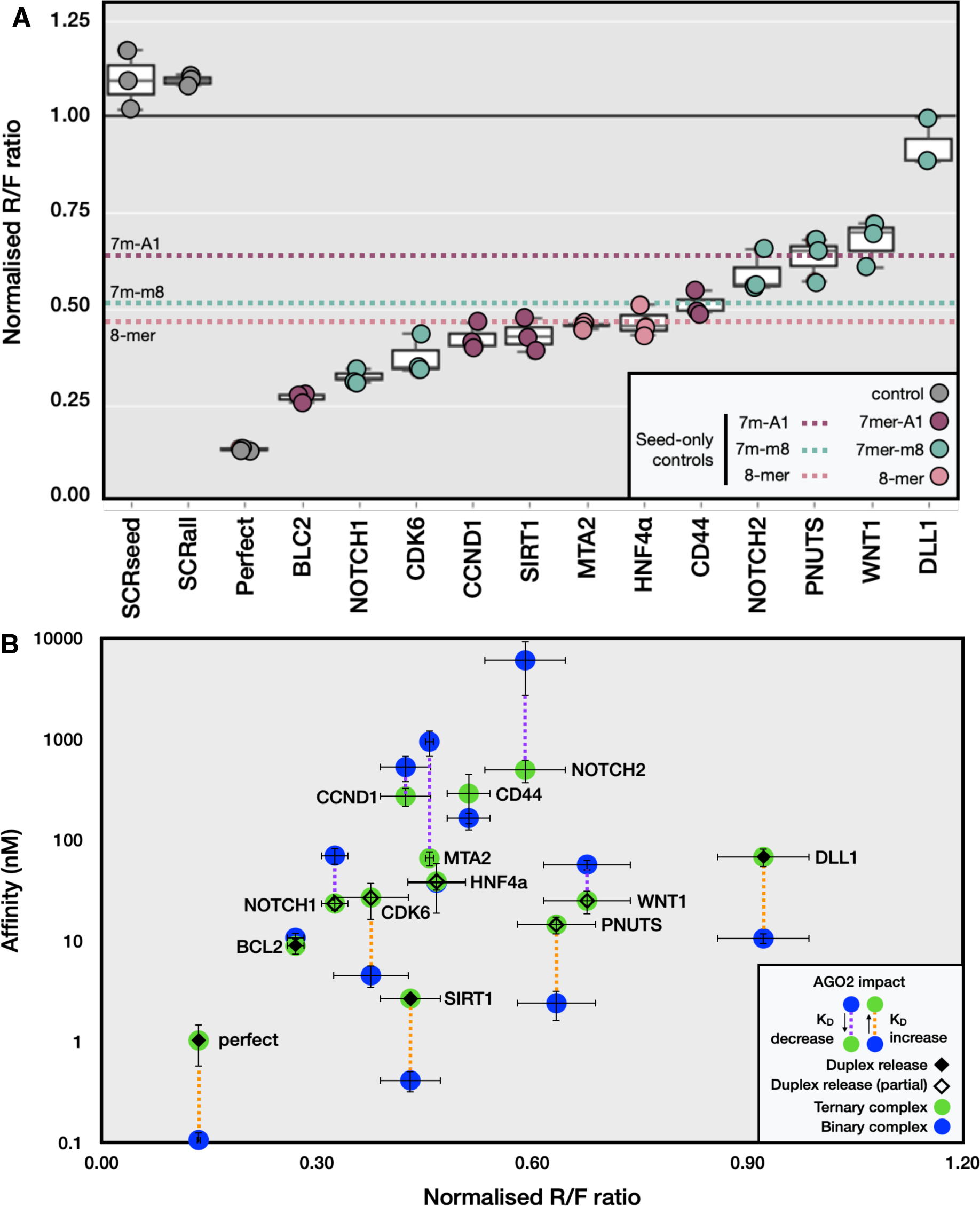
AGO2 moderates affinity by strengthening weak binders and weakening strong binders. (A) *miR-34a*-mediated repression of dual luciferase reporters fused to the 12 mRNA targeting sites. Luciferase activity from HEK293T cells co-transfected with each reporter construct, *miR-34a* was measured 24 hours following transfection and normalised to the *miR-34a*-negative transfection control. Each datapoint represents the R/F ratio for an independent experiment (n=3) with standard deviations indicated. SCRseed is a scrambled seed control, SCRall is a fully scrambled control, and Perfect is the perfect complement of *miR-34a*. Dotted horizontal lines represent the repression values for the 22-nucleotide seed-only controls^6^ for the respective seed types, in the absence of any other WC base pairing (7m = 7mer). (B) Comparison of relative target repression with relative affinity assessed by EMSA. Blue represents mRNA:*miR-34a* affinity (binary complex), while green represents mRNA:*miR-34a*-AGO2 affinity (ternary complex). Solid diamonds represent miRNA duplex release, open diamonds represent partial duplex release.

**Figure 4.**
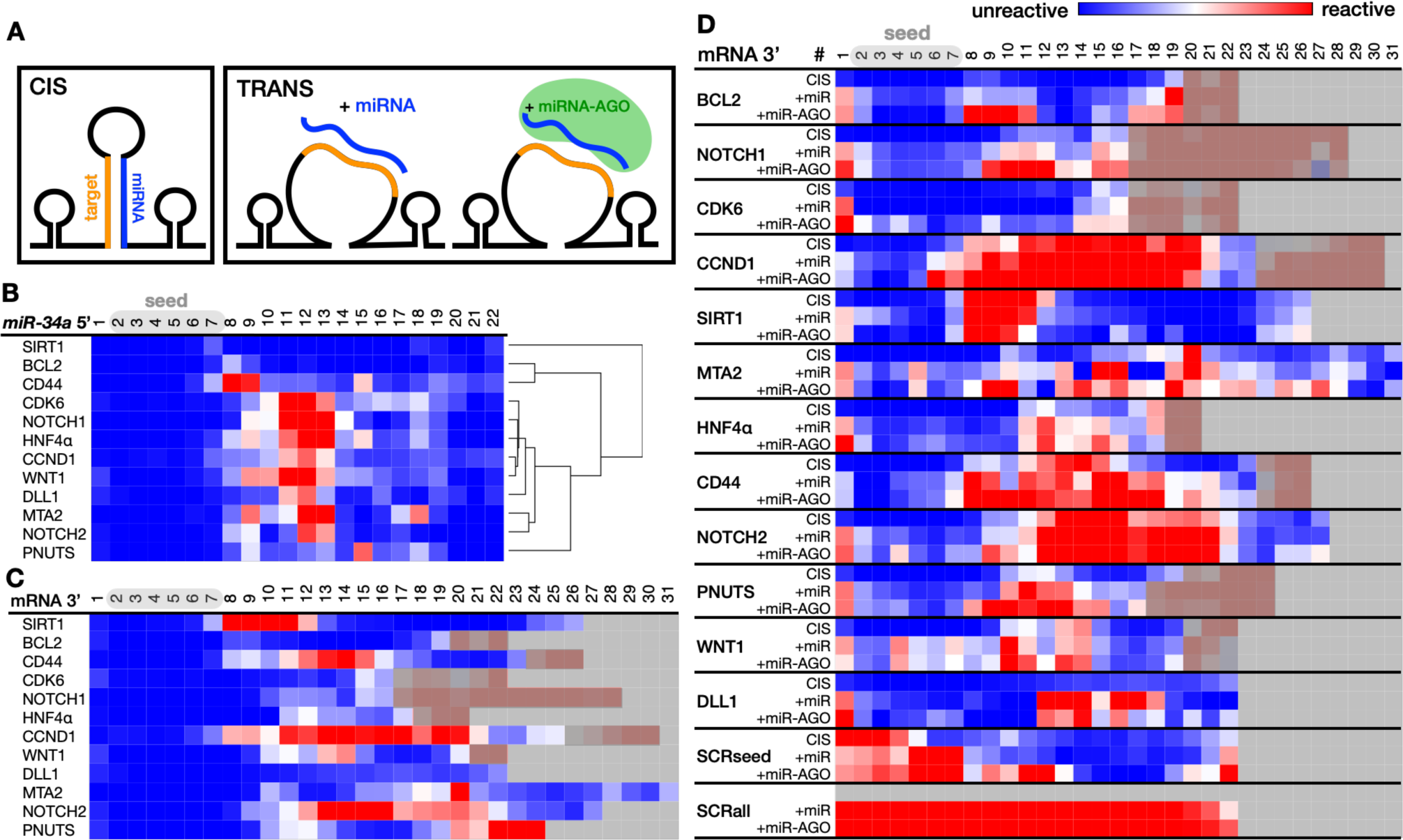
Structural probing by RABS reveals that structural features are established in the binary complex. (A) Illustration of RABS CIS and TRANS scaffolds in which mRNA targets binding to *miR-34a*. (B) CIS reactivity values of *miR-34a* nucleotide positions 1-22 when bound to each of the 12 target mRNAs. *miR- 34a* nucleotide positions are numbered from the 5‘ end (seed). Targets are clustered hierarchically according to their reactivity patterns, using one minus cosine similarity as a distance metric, and linked by mean average. (C) CIS reactivity values for each mRNA target when bound to *miR-34a*. mRNA targets are numbered from their 3’ end (seed) and are ordered hierarchically as in (B). (D) Comparison of reactivity values for each mRNA target when bound to *miR-34a* in CIS or TRANS (top two rows respectively), and to *miR-34a*-AGO in TRANS (lower row). The mRNA order along the *y*-axis corresponds to the degree of repression efficacy, with most repressed at the top. The values in panels B, C, and D are colour-coded to represent SHAPE reactivity levels: blue indicates low reactivity being base paired, red indicates high reactivity (i.e., no involvement in stable base pairing), and white marks the cut-off point for base pairing (at 0.5 reactivity). Grey boxes signify where mRNA constructs end. Red-grey regions in the mRNA tail are not involved with miRNA pairing but exist within the construct. The SCRseed (scrambled seed) control lacks a seed-match but contains a *miR-34a* complementary tail. SCRall lacks any stable base pair formation with *miR-34a*.

**Figure 5.**
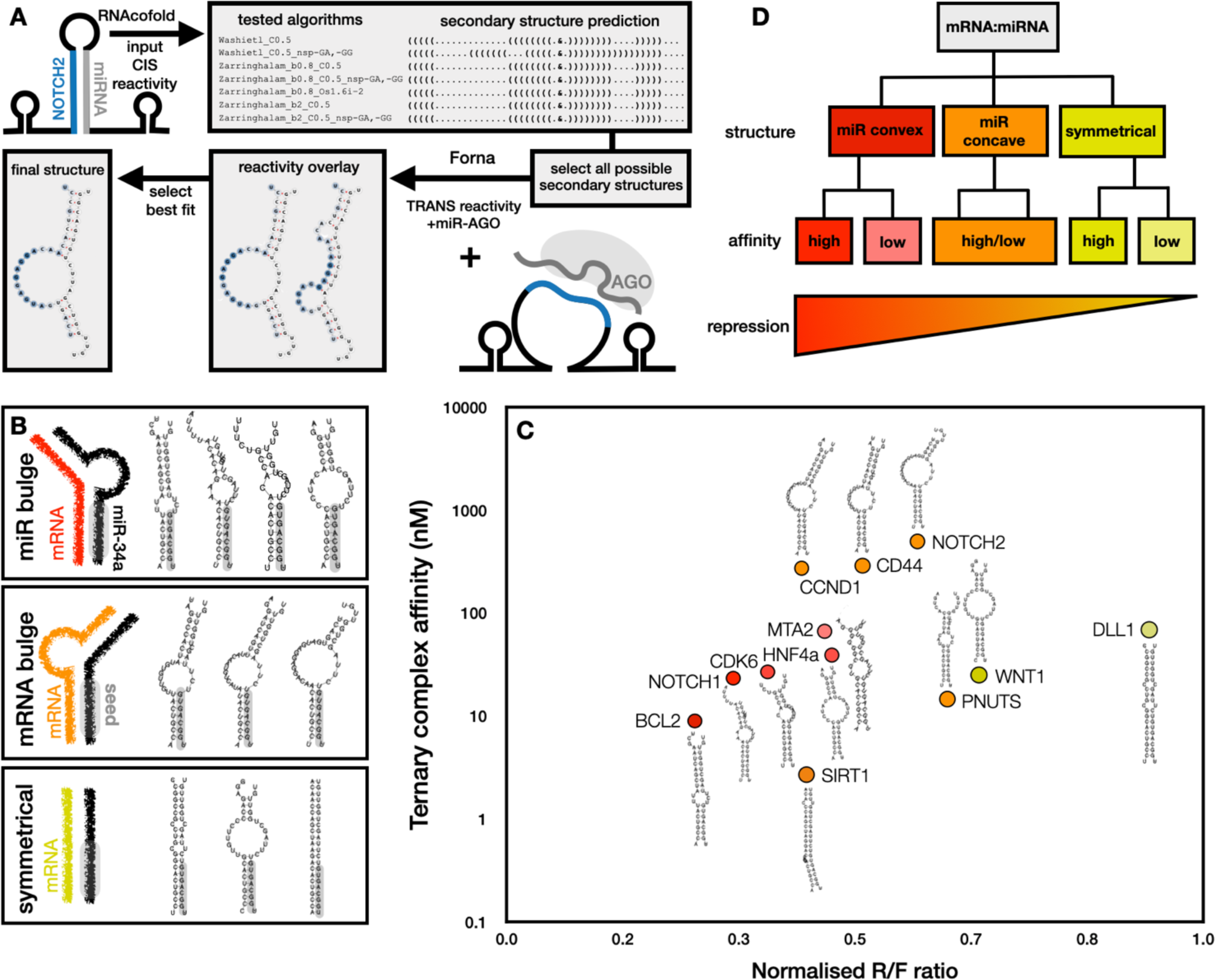
mRNA targets form three distinct structural groups. (A) Overview of the secondary structure prediction pipeline combining CIS and TRANS RABS, using the example of *NOTCH2*. The CIS binary complex (mRNA:*miR-34a*) reactivity was used to calculate possible secondary structures in RNAcofold, using a variety of algorithms^42,44^ as shown. The possible structures were then overlaid with TRANS ternary complex (+*miR- 34a*-AGO2) reactivity data to select the most representative structure. (B) Three groups of structures were revealed: bulged on the miRNA side, bulged on the target side, and symmetrical. (C) The three groups cluster together visually in relation to their ternary complex affinity and repression efficacy. The miRNA bulge group (red) had high repression efficacy and correlation to affinity (R^2^ = 0.85; Figure S7), while the mRNA bulge group (orange) had moderate repression with no correlation to affinity (R^2^ = 0.002; Figure S7), and the symmetrical group (dark yellow) appear to be disfavoured by *miR-34a*. (D) Observed trend relating structural class, affinity, and repression efficacy.

### Binary complex affinity dictates ternary complex affinity

A linear correlation was observed between the log-transformed binary and ternary complex affinities (R^2^ = 0.72; Figure 2B). Since the standard change in free energy for a reaction, ΔG°, is related to its equilibrium constant, *K* (in this case *K*_D_) by the equation ΔG° = -RTln*K*, comparison of the log transformed affinity values is proportional to ΔG. It follows that the ΔG of the binary and ternary complexes are highly correlated, and ΔG of the ternary complex is likely dictated by RNA:RNA interaction. The slope of the linear fit is 0.48, with (log *y*) intercept of 7.11 nM, thus there is an approximately ten-fold reduction in the range of affinities upon binding *miR-34a-*AGO compared to *miR-34a* only binding (Figure 2B). Interestingly, however, we did not observe a correlation between binary or ternary complex affinity and seed type.

In the ternary complex binding assays, a strong intermediate band emerged between the free and protein-bound bands when *miR-34a*-AGO2 was titrated against specific targets (*SIRT1*, *BCL2*, *DLL1*, and *CDK6,* as well as the Perfect and SCRseed controls). This intermediate band was only faintly or partially observed for *WNT1*, *HNF4α*, *NOTCH1*, and *PNUTS*, and it was entirely absent for *MTA2*, *CCND1*, *CD44*, and *NOTCH2* (Figure S4). To identify the nature of the intermediate band, we conducted a northern blot (Figure S5B). An *miR-34a* probe hybridised to the intermediate band and was observed at a similar height as a *miR-34a-SIRT1* duplex, indicating that the *miR-34a*:mRNA duplex is released from AGO2 (referred to as duplex release). Although this phenomenon has been previously observed^22,23,55^, its biological significance remains incompletely understood.

During development of our AGO2 binding mobility shift assays, we noticed that the presence of an RNA background (either polyuridine (pU) or total RNA extract) resulted in an increased specific AGO2 binding for low-affinity targets (Figure S6B-C). In the absence of this RNA background, all low-affinity targets (including the negative control SCRall) bound with similar affinity, at approximately 5 nM *miR-34a* loaded AGO2 (Figure S6A). Rather than displaying a typical sigmoidal binding curve, the targets instead bound with an abrupt ‘step’ at the 5 nM titration point, indicating that non-specific interaction with AGO2 was also occurring. Argonaute proteins have been observed to bind nucleic acid non-specifically, even in the absence of a guide^56^. Here, pU acts as a blocking agent to compete out non-specific interactions.

The mRNA:*miR-34a*-AGO2 assay had a limited titration range, reaching a maximum *miR- 34a*-AGO2 concentration of 268 nM due to a 5% loading efficiency (see Figure S2D for loading efficiency quantification). The total AGO2 concentration was 20-fold higher than the *miR-34a*-loaded portion, and further increase was hindered by protein precipitation. Weaker mRNA targets (*CD44*, *CCND1*, and *NOTCH2*) didn’t reach a saturated binding plateau within this range, leading to larger errors in their estimated *K*_D,app_ values. However, reasonable estimation of the *K*_D,app_ was possible by monitoring the disappearance of the free mRNA probe. To ensure AGO2 binding specificity despite low loading efficiency, a scrambled control was used (SCRall; lacking stable base pairing with *miR-34a* or other human miRNAs according to the miRBase database^57^). SCRall showed no interaction with *miR-34a*-AGO2 (Figure S4).

### Affinity as a contributing determinant of site effectiveness

Repression efficacy for the 12 mRNA targets by *miR-34a* was assessed through a dual luciferase reporter assay^6^. Expression values spanned a range from 73% downregulation compared to the *miR-34a*-negative control level (*BCL2*; mean R/F ratio = 0.27 ± 0.012; Figure 3A) to minimal repression (*DLL1*; mean R/F ratio = 0.922 ± 0.064; as shown in Figure 3A), with ‘perfect’ as the positive control indicating maximum possible down- regulation efficiency (mean R/F ratio = 0.135 ± 0.003) via the siRNA pathway. The repression efficiencies of the targets between these two extremities were evenly distributed, forming an optimal range for our analytical purposes. Notably, there was no discernible correlation between the seed type nor duplex release and the observed order of relative repression (Figure 3A). As seed type has been shown to be a major determinant of repression efficacy^7,58^, we were interested in differences that alter a target’s downregulation from the expected levels based on the seed alone. To this end, we compared target repression levels with 22 nucleotide seed-only control targets, each containing a seed match for the respective seed type but no complementary pairing outside of the seed^6^. The two 8-mer targets, *MTA2* and *HNF4α*, exhibited no significant variation in their repression levels compared to the corresponding 8-mer seed-only control. In contrast, all 7mer-m8 and 7mer-A1 targets demonstrated distinct differences from their respective seed-only controls (Figure 3A). 7mer-A1 targets *BCL2*, *CCND1*, *SIRT1* and *CD44* each exhibited more effective repression than the 7mer-A1 control. For the 7mer-m8 targets, *NOTCH1* and *CDK6* were more effective than their corresponding control, while *NOTCH2*, *PNUTS* and *WNT1* were less effectively repressed (Figure 3A). This result indicates that the impact of supplementary binding is greater for targets with weaker seeds.

We then compared the luciferase repression data with the target affinities (Figure 3B). While there was a stronger correlation to the ternary affinities than binary, a weak positive linear correlation emerged between repression level and the log of both binary and ternary affinities, accounting for 8 % and 23 % of variation within the repression dataset respectively (R^2^ = 0.08 and 0.23; Figure S7A-B). As previously observed in Figure 2B, there was a significant reduction, approximately tenfold, in the overall span of mRNA:*miR34a*-AGO2 affinities compared to the mRNA:*miR34a* range. This effect is further highlighted in Figure 3B, where it is apparent that strong binary complex affinities weaken upon ternary binding, whereas weak binary complex affinities strengthen. This observation demonstrates that AGO2 exhibits a bidirectional capacity to modulate binding affinity.

### Duplex release occurs in targets with strong binary complex affinity

Our affinity analysis also provided insight into the miRNA duplex release phenomenon. It is evident that tighter binders (*K*_D,app_ less than 100 nM) result in duplex release, whereas weaker binders do not exhibit this behaviour (Figure 3B). This trend was observed to varying degrees: tight ternary complex binders such as Perfect, *BCL2*, *SIRT1*, and SCRseed demonstrate strong duplex release, while moderate binders *NOTCH1*, *CDK6*, *HNF4α*, *PNUTS* and *WNT1* all exhibited partial duplex release (Figure S4 and Table S2). An intriguing exception to this trend was *MTA2* (ternary complex *K*_D,app_ = 66.1 ± 11.2 nM), which did not exhibit duplex release despite sharing a similar ternary affinity to *DLL1* (*K*_D,app_ = 69.2 ± 14.1 nM), which exhibited distinct duplex release. Notably, *DLL1* had a stronger binary complex affinity (*K*_D,app_ = 10.6 ± 1.13 nM) whereas *MTA2* has a significantly weaker binary affinity (*K*_D,app_ = 944.3 ± 274), suggesting that duplex release might be governed by binary affinity rather than ternary affinity. In the case of SCRseed control, it is notable that the seed region binding does not appear to be necessary for duplex release.

Evidently, even when considering only the isolated binding site and surrounding sequence factors are excluded from consideration, affinity effects only provide a partial explanation for targeting outcomes. This led us to investigate whether structural characteristics within the targeting site could serve as contributing determinants to the efficacy of repression.

### Structural determinants of mRNA:miRNA-AGO2 binding

#### miRNA bound at seed and tail

To infer base pairing patterns and secondary structure for each of the 12 mRNA:*miR-34a* pairs, we used the RABS technique^29^ (Figure 4A). All individual reactivity traces are shown in Figure S9. Reactivity of each of the 22 *miR-34a* nucleotides was assessed upon binding to each of the 12 mRNA targets within a CIS construct, containing both *miR-34a* and the mRNA target site that can bind within the same RNA scaffold. *miR-34a* was unreactive between nucleotide positions 1-7 and 19-22 (blue regions; Figure 4A), indicating both the *miR-34a* seed and tail regions are strongly bound, regardless of the target. The central and supplementary regions between *miR-34a* positions 8-18 exhibited variable reactivity across the targets. Hierarchical clustering of the targets according to their reactivity values revealed that positions 11-13 (equivalent to an offset central region) strongly distinguished two target groups. Targets *CDK6*, *NOTCH1*, *HNF4α*, *CCND1*, *WNT1*, *DLL1*, *MTA2* and *NOTCH2* were reactive between positions 11 - 13, while *SIRT1, BCL2*, *CD44* and *PNUTS* were not (Figure 4A). The latter four targets, which showed no reactivity in positions 11- 13 signifying strong binding to *miR-34a* across the seed and central regions, displayed no apparent correlation with their relative repression efficacy. Notably, *BCL2* and *SIRT1* were highly repressed, while *CD44* and *PNUTS* were only weakly repressed. Moreover, the miRNA supplementary region (typically spanning positions 13-16), did not exhibit distinct binding (Figure 4A). Instead, an offset supplementary equivalent spanning *miR-34a* positions 14–18 displayed substantial binding for all targets, though not as strongly as seen in the tail positions (19-22).

#### Asymmetric target binding

The mRNA:*miR-34a* pairs exhibited diverse reactivity on the mRNA target side (Figure 4B). Unlike the miRNA side, no distinct reactive central region was observed, and the reactivity patterns did not mirror those observed on the miRNA side, indicating an asymmetry in the duplex structure. The total nucleotides engaged in *miR-34a* interaction varied, highlighting that mRNA target sites are not confined to the commonly assumed 22- nucleotide region for target prediction. The variability in reactivity across mRNA target sites underscores the absence of strict rules for binding positions in the seed, central, supplementary, and tail regions. Instead, diverse sequence positions contribute to these interactions, indicating the presence of structural features enabling such binding arrangements.

#### AGO increases flexibility, but does not alter binding pattern

To investigate whether AGO2 alters the structure established between miRNA and mRNA target, we employed TRANS constructs in our RABS analysis. This allowed us to examine how the target’s structure changes upon introduction of a binding partner (either *miR-34a* or *miR-34a*-AGO2). In TRANS, the target site is located within an RNA scaffold and only target reactivity is observable (as illustrated in Figure 4A). Comparison between CIS and TRANS revealed that the underlying reactivity patterns resulting from mRNA:*miR-34a* binding were preserved upon mRNA:*miR-34a*-AGO2 binding (Figure 4C). As the reactivity in the ternary complex matches that in binary, we can exclude protein occlusion effects from our RNA structure analysis. It follows that the structure imparted via direct RNA:RNA interaction remains intact within AGO2, highlighting the role of the RNA as the structural determinator.

Comparison of CIS and TRANS reactivity patterns revealed a stabilising effect of the CIS construct. There was a tendency for the reactive regions to increase in magnitude in TRANS, attributed to an energetically favourable contribution of the connecting loop^59^ in CIS, resulting in overall reduced CIS reactivity values. In TRANS, an increase in reactivity was observed upon + *miR-34a*-AGO2 binding, indicating that AGO2 imparts greater flexibility while preserving the structural features formed in the binary complex. *SIRT1* and *DLL1* were minor exceptions, exhibiting slightly decreased reactivity within their central reactive regions when bound to *miR-34a*-AGO2 compared to *miR-34a* alone (Figure 4C). *MTA2* presented multiple binding conformations, evident from the multiple bands on native gels (Figure S3 and S4), likely leading to the incoherent reactivity patterns observed (Figures 4C and S9).

### Three distinct structural groups form upon mRNA:*miR-34a* binding

To decipher the reactivity patterns observed in RABS, we combined both CIS and TRANS reactivity data to predict the secondary structures of each mRNA:*miR-34a* duplex. Firstly, we evaluated a range of possible structures based on the CIS data (comprising reactivity data from both the *miR-34a* and mRNA sides in the binary complex) using the Vienna RNA tool RNAcofold^37^. Subsequently, we superimposed these structural possibilities with TRANS +*miR-34a*-AGO2 reactivity data (ternary complex) to ultimately select the most representative predicted secondary structure (Figure 5A). All RABS reactivity traces and corresponding secondary structures are shown in Figure S9.

Varied structural features were observed across the mRNA:*miR-34a* pairs, forming three distinct groups: a bulge present on the *miR-34a* side, a bulge on the mRNA side, and symmetrical internal loops (Figure 5B). Each of the 12 targets exhibited one of these three structural types upon binding within the ternary complex. The bulge was typically positioned after the seed, commencing between position 8 and 11 on the *miR-34a* side, considered the central region (Figure 5C and S9). To compare bulge size, we considered asymmetrically unpaired nucleotides, i.e., symmetrical mismatches were not counted. On the *miR-34a* bulge side, asymmetry varied from 1 nucleotide (*BCL2*) to 5 nucleotides (*CDK6*). On the target side, asymmetry ranged from a single nucleotide (*PNUTS*) up to 9 nucleotides (*NOTCH2*).

We hypothesised that the orientation of the bulge and the bend of the duplex might play a role in how the duplex aligns within AGO2, ultimately affecting the level of repression achieved. Notably, the targets *BCL2*, *NOTCH1*, *CDK6*, *HNF4α*, and *MTA2* all exhibit a miRNA bulge structure, and had high relative repression efficacy (Figure 5C). Further, the miRNA bulge group had a strong positive correlation between repression and ternary complex affinity (R^2^ = 0.85; Figure S7C). It follows that a bulge positioned on the miRNA side could confer an advantage, resulting in effective repression, with differences in affinity ‘fine tuning’ the repression level. Conversely, when a bulge is situated on the target side, reduced repression may result. This was evident in the case of *SIRT1*, which was a tight binder to *miR-34a*-AGO2 (*K*_D,app_ = 2.7 ± 0.4 nM). However, *SIRT1* may face a ’penalty’ due to its mRNA bulge, resulting in similar repression levels to miRNA bulge counterparts with tenfold weaker affinities (*CDK6*, and *HNF4α*; Figure 5B). This penalty may be due to potential formation of a bend in the RNA duplex in the opposite direction compared to the miRNA bulge group. Interestingly, all targets exhibiting an mRNA bulge conformation were similarly repressed (R/F ∼ 0.5), despite a large spread in affinity values. As such, the mRNA bulge group lacked any correlation between repression and affinity (R^2^ = 0.002; Figure S7C). These combined observations emphasise the significance of structure, and specifically bulge location, in contributing to repression efficacy.

The final group of targets comprised those forming symmetrical duplexes, exemplified by *WNT1* and *DLL1*. These targets exhibited only modest repression (with R/F ratios of 0.68 and 0.92 respectively; Figure 3A and Table S2) despite possessing relative *miR-34a*- AGO2 affinities comparable to *CDK6* (*K*_D,app_ = 26.9 ± 1.4 nM). *CDK6* however, belonging to the mRNA bulge group, was much more effectively repressed (R/F = 0.37). It appears that the rigidity of the structure is also important; *WNT1* has a larger internal loop consisting of 7 mismatches leading to larger potential flexibility, while *DLL1* is more rigid with two single mismatches only. Moreover, *PNUTS* presented a near symmetrical configuration (mRNA bulge with 1 nucleotide asymmetry) and displayed relatively poor repression compared to its ternary complex affinity (R/F = 0.63 ± 0.06, *K*_D,app_ = 14.5 ± 2.9 nM), indicating a potential role of rigidity in this case. It should be noted that the ’Perfect’ control, although symmetrical, can undergo repression through the siRNA pathway (via slicing), rendering it incomparable in this context.

The structural orientation of the *miR-34a*:mRNA duplex may not only be important in determining repression efficacy, but also in determining whether duplex release occurs. Notably, the four targets that did not exhibit duplex release (*CD44*, *CCND1*, *NOTCH2*, and *MTA2*) had secondary structures belonging to the mRNA bulge category with asymmetry of at least 5 nucleotides, potentially creating a larger bend in the duplex (Figure S10A). Note that *MTA2* is expected to exist in multiple *miR-34a*-bound confirmations based on native gel band patterns (Figure S3A and S4) and incoherent RABS reactivity patterns (Figure 4C). Thus, *MTA2* had two plausible secondary structures: an mRNA bulge structure with 12-nucleotide asymmetry, and a miRNA bulge structure with 2- nucleotide asymmetry. Altogether, it is possible that larger asymmetrical bulges (>4 nucleotides) on the mRNA side prevent release of the mRNA:*miR-34a* duplex.

The structural characteristics of each *miR-34a*:mRNA duplex exhibit a notable correlation with affinity, repression efficacy, and duplex release activity. This observation underscores the previously underappreciated significance of RNA:RNA structure in the context of miRNA targeting. Recognising the potential of RNA structure in deciphering miRNA targeting outcomes, we subsequently explored whether these structural attributes held relevance in the three-dimensional configuration of a mRNA:*miR-34a* duplex within AGO2. While there is ample evidence that AGO2 can readily accommodate mRNA:miRNA complexes containing bulges on the target strand^6,10,17,18,26,60^, our results show that AGO2 is able to bind complexes with a pronounced bulge on the miRNA strand as well. This is somewhat counterintuitive, as existing structures would suggest miRNA bulges to be sterically incompatible within the AGO2 nucleic acid binding cleft. To ascertain the structural plausibility of a stably formed ternary complex with a large miRNA-side bulge, we built a 3D structural model of the *NOTCH1*:*miR-34a*-AGO2 ternary complex. Starting with the existing crystal structure of *miR-122*-AGO2 bound to a seed and supplementary paired target^17^, the structure was mutated in-place to match the sequence of the *NOTCH1*:*miR-34a* complex while maintaining the secondary structure depicted in Figure 5B, and equilibrated in multiple, independent unrestrained molecular dynamics simulations. We find that the pronounced miRNA-side bulge is readily accommodated in a stably formed ternary complex over the course of multiple, independent simulations. This is due to the ability of the bulged region to loop out in a highly compact manner and form contacts with the PAZ domain (Figure 6B), while stably maintaining contacts between the 5’ seed region in the AGO2 seed binding cleft (Figure S11). The 3’ supplementary region flexibly bends and interacts with the N-terminal domain, reminiscent of the TDMD-competent state with 3’ tail release from the PAZ domain^19^ (Figure 6C), and aligning with the observation that *NOTCH1*:*miR-34a* releases from AGO2 as seen via EMSA (Figure S4). The stability of this model in explicit solvent MD simulations suggests that AGO2 can accommodate a wider range of mRNA:miRNA conformations than previously assumed.

**Figure 6.**
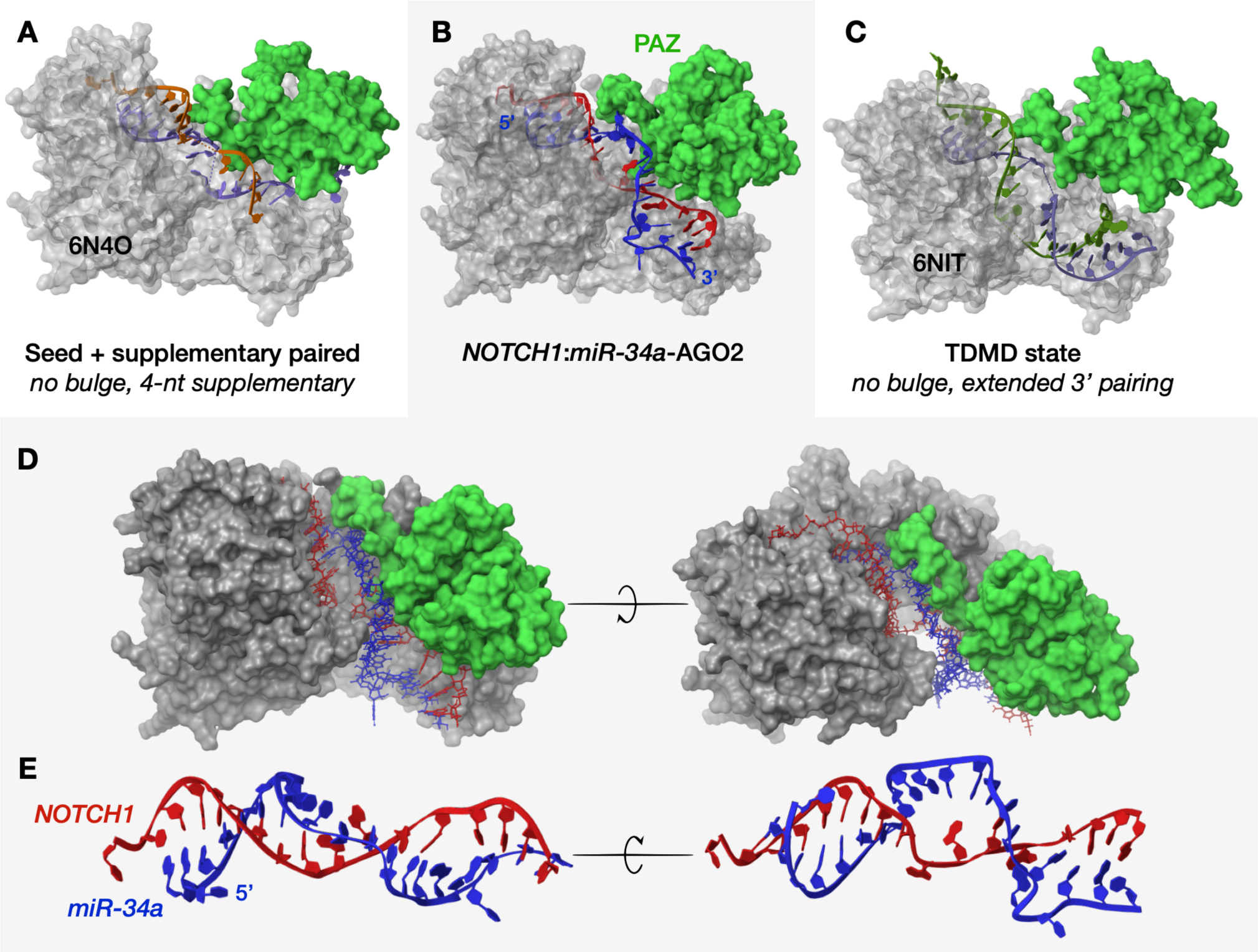
The miRNA-bulge structure is readily accommodated by AGO2 as shown by molecular dynamics simulation. (A) Crystal structure of *miR-122* (purple) paired via the seed and 4 nucleotides of supplementary pairing to its target (orange), PDB-ID 6N4O^17^. (B) All-atom MD simulation of *miR-34a* (blue) and *NOTCH1* (red) in AGO2. The MD simulation details are provided in Supplementary File 7. (C) Crystal structure of a *miR-122* (purple) and target (olive green) with extended 3’ pairing (TDMD-competent state), PDB-ID 6NIT^19^. AGO2 is shown in grey with PAZ domain in green for all structures. The RNA duplexes in (A) and (C) have a symmetrical 4-nucleotide internal loops whereas (B) features an asymmetrical miRNA-bulge, containing 5 unpaired nucleotides (*miR-34a* positions 10-14), with 3-nucleotide asymmetry. (D) The *NOTCH1*:*miR-34a*- AGO2 structure is rotated to show the RNA binding cleft and accommodation of the miRNA bulge (left) and stable binding of the seed region (right). (E) The *NOTCH1*:*miR-34a* duplex with AGO2 removed for clarity.

## DISCUSSION

Three decades after the discovery of the first miRNA, current prediction algorithms can only account for approximately half of the variability in mRNA repression caused by miRNAs^11^. This indicates the presence of as-yet-unidentified factors that impact targeting effectiveness and remain unaccounted for in predictive models. Here, we explored additional factors influencing miRNA targeting efficacy by isolating and examining the binding site itself, excluding surrounding sequence factors from consideration.

Using the tumour suppressor *miR-34a* as a subject, we studied the interaction of twelve mRNA targets in high detail. We determined binding affinities of each target for both *miR- 34a* and *miR-34a*-AGO2 via EMSA and revealed that AGO2 has a bidirectional capacity to modulate affinity, decreasing the affinity of strong mRNA:*miR-34a* pairs while increasing the affinity of weaker mRNA:*miR-34a* pairs (Figure 3B). Using RABS to determine secondary structure and luciferase reporter assays to determine repression levels, we revealed that three distinct structural groups are formed upon binding of the targets to *miR-34a*, and that both structure and affinity play a role in determining target downregulation (Figure 5C) and duplex release (Figures 3B and S10A). We further observed that interaction of a target within the ternary complex appears to be controlled at the RNA level, as mRNA:*miR34a* and mRNA:*miR34a*-AGO affinities are highly correlated (Figure 2B), and reactivity patterns observed upon mRNA:*miR34a*-AGO2 binding follow those established upon mRNA:*miR34a* binding (Figure 4C). In addition, structural probing revealed flexibility of the 3’-supplementary location and underscored importance of the tail, confirming our earlier findings on the *SIRT1*:*miR-34a* interaction^6^. Our results suggest that structural characteristics of a mRNA:miRNA binding site could serve as contributing determinants to the efficacy of repression.

### RNA:RNA binding assays provide valuable predictive information

An earlier examination of mRNA:miRNA binding thermodynamics found that mouse and fly AGO2 reduce the affinity of a guide RNA for its target^61^. Our data indicate that the range of binary complex affinities is instead constricted by human AGO2 in the ternary complex – strengthening weak binders while weakening strong binders. Notably, binary complex affinity was found to be a strong predictor of ternary affinity (Figure 2B, R^2^ = 0.72). As such, the affinity of target binding by *miR-34a*-AGO2 binding can thus be approximated via direct RNA:RNA hybridisation strength. Employing RNA:RNA binding assays offers the advantages of simplicity and enhanced reliability in experimentation. TargetScan predicts biological targets of miRNAs by recognising conserved 8mer, 7mer, and 6mer sites matching the seed region of each miRNA^4^. However, our affinity dataset shows no significant linear correlation with TargetScan predicted relative K_D_ for the 12 studied *miR-34a* targets (Figure S8B). While Gibbs free energy (G) is often included in targeting prediction models as a measure of stability of the miRNA:mRNA pair^12,62^, our *K*_D,app_ values display only modest correlations with the free energy values derived from the minimum free energy (MFE) structures predicted by RNAcofold (R^2^ = 0.13 and 0.2 for binary and ternary complexes respectively; Figure S8A). Disparities in K_D_ values may arise from the different target lengths used in our study compared to Targetscan entries, which consider only a 23-nucleotide target site. Notably, TargetScan predicted relative K_D_ values do correlate with our luciferase dataset (R^2^ = 0.43; Figure S8B), indicating consideration of parameters beyond *in vitro* binding affinity in the TargetScan algorithm. The lack of correlation between predicted and observed values may stem from inaccurate representation of the true binding mode in predicted secondary structures, leading to imprecise energy values and poor predictions of miRNA targeting efficiency. Investigating the actual binding mode thus is crucial for understanding the true energetic contribution to binding, but given the challenge of determining it *a priori*, using RNA:RNA binding affinity as a proxy for ternary complex affinity offers a potential solution, and minimises resource investment.

### Three distinct structures are observed upon target binding to miR-34a

Recent literature highlights that binding to both the supplementary and tail regions, but not the central region of miRNA, enhances target repression^6,17^. The supplementary region has typically been defined as nucleotides 13-16 of the miRNA, the tail region nucleotides 17-22, and the central region nucleotides 9-12. Our RABS results suggest that these regions should instead be defined as successive base-paired blocks but not by strict nucleotide numbering systems. We observed that the reactive central region of *miR- 34a* varied between positions 9-13, most commonly involving nucleotides 11-13. This deviates from the common definition of the central region containing mismatches in positions 9-11^17,63^. Interestingly, this reactivity was not mirrored symmetrically on the mRNA side, indicating the existence of asymmetrical structural features within the duplex (Figure 4). Additionally, we observed that the nonreactive nucleotides following the central region, indicative of supplementary binding, varied in position both on the *miR-34a* and mRNA sides. In contrast, the *miR-34a* nucleotides 20-22 were unreactive for all target interactions (Figure 4A), despite differences in corresponding mRNA length and SHAPE reactivity. The *miR-34a* tail nucleotides 20-22 were the only positions outside of the seed that were consistently unreactive. Taken together, these results indicate that aside from the miRNA seed and tail, binding regions within an mRNA:miRNA pair are highly variable and cannot be predicted through sequence alignment alone (Figure S10B).

The asymmetrical nature of the reactivity patterns can be explained by secondary structure features. Prior studies have revealed that the miRNA central region (nucleotides 9-11) is unpaired within AGO2^17^, and that AGO2 can accommodate a bulge or loop on the mRNA target up to 15 nucleotides in length, located between the seed and supplementary regions^6,17,64^. While our structural data align with the existence of bulges on the target, we also observed that bulges can be accommodated on the miRNA side (Figure 5). We provide structural evidence to demonstrate that miRNA bulges can exist within native sequences in the ternary complex (Figures 5 & 6) and have observed bulges with up to 5 nucleotide asymmetry (for example *CDK6*). This finding is consistent with a recent publication where siRNA modified with one or two nucleotide bulges on the guide strand were shown to be compatible with gene repression *in vivo*^65^.

We also observed the existence of a third structural group that can form upon mRNA:*miR- 34a* binding: symmetrical pairs. These three groups (here named miRNA bulge, mRNA bulge, and symmetrical) differed not only in terms of bulge location, but also in terms of relative repression efficacy (Figure 6C&D). Specifically, the miRNA bulge group was most effectively repressed and correlated strongly with ternary complex affinity (R^2^ = 0.85), the mRNA bulge group less effectively repressed and did not correlate with ternary complex affinity (R^2^ = 0.002), and the symmetrical group had poor relative repression (Figure 5C). We propose that the bulge location and degree of asymmetry may determine the mRNA:miRNA duplex angularity and overall 3D structure within AGO2, leading to the observed differences in repression efficacy.

#### mRNA:miRNA duplex release

Certain mRNA targets have been shown to induce mRNA:miRNA release from Argonaute, however, the rules governing the process remain elusive. Our results revealed a general trend that *miR-34a* targets with higher affinity exhibited duplex release, while those with moderate affinity exhibited partial release (Figure 3B), aligning with a previous report showing that highly complementary targets lead to guide RNA release^22^. More specifically, our findings indicate that the binary complex affinity may be more important in determining duplex release than the ternary complex, as tighter binary complexes can release despite weaker ternary complex affinity (Figure 3). We show that all canonical seed-matched targets (8mer, 7mer-m8 and 7mer-A1) can trigger duplex release, however, the seed is not necessary as a scrambled seed control (SCRseed; lacking a seed-match but containing a *miR-34a* complementary tail) also exhibited clear duplex release (Figure S5)). We observed a relationship to the structural class of the mRNA:miRNA duplex, where mRNA bulges with high asymmetry (>4 nucleotides) appear to impede duplex release (Figure S10A), consequently diminishing AGO’s recycling capacity. These findings highlight the importance of miRNA:mRNA interactions and the structural nuances at play. Recognising that secondary structure provides limited insight into overall geometry, further studies should consider 3D structural aspects for a more comprehensive understanding of base pair patterns and their relevance in protein binding. Redefinition of the miRNA seed, central, supplementary, and tail regions by their successive order and structural roles, rather than by numbered nucleotide positions, warrants consideration. Furthermore, RNA:RNA interactions may be more important in guiding biological outcomes within AGO than previously appreciated.

#### Impact on miRNA targeting prediction and drug design

Current bioinformatic strategies fall short in accurately predicting miRNA-mediated repression efficiency. We propose that the structural characteristics of miRNA targeting play a pivotal role, and integrating structural insights into targeting prediction algorithms has the potential to enhance the accuracy of existing models. Using structural insights may also prove critical in the successful design of miRNA-based therapeutics. Until now, endeavours to employ modified *miR-34a* as a therapeutic agent have encountered setbacks in clinical trials^66^. These failures have been attributed to issues of non-specificity and high drug dosages, which can trigger immunogenic responses^67^. By tailoring miRNA-based drugs based on known structural information, we may have the potential to create more effective treatments characterised by a reduced immunogenic profile and a more selective targeting approach. This strategy could also broaden the range of molecules that can be targeted for therapeutic purposes.

## DATA AVAILABILITY

Raw data from binding assays, luciferase reporter assays, structural probing, secondary structure predictions, analysis code, and molecular dynamics simulations are included as supplementary files.

## AUTHOR CONTRIBUTIONS

L.S.: Conceptualisation, Methodology, Investigation, Data curation, Formal analysis, Validation, Visualisation, Writing – draft, review, and editing. D.K.: Methodology, Investigation. E.B., W.B., J.M., C.K. and A.C.: Investigation. L.B. and I.G.: Conceptualisation. D.S.: Software. E.R.A and K.P. Conceptualisation, Supervision, Project administration, Writing – review & editing.

## ACKNOWLEDGEMENTS

1. J. Weidhaas for the psiCHECK2 vector gift from (Addgene plasmid 78258), The Protein Science Facility at Karolinska Institute for production of T7 polymerase and inorganic pyrophosphatase, the Petzold and Andersson Labs for impactful discussion.

