## Supplementary Information for "Biophysics of microRNA-34a targeting and its influence on down-regulation"

General information

All standard chemicals and solvents were purchased from the commercial suppliers Sigma-Aldrich, Thermo Fisher, or VWR unless otherwise specified.

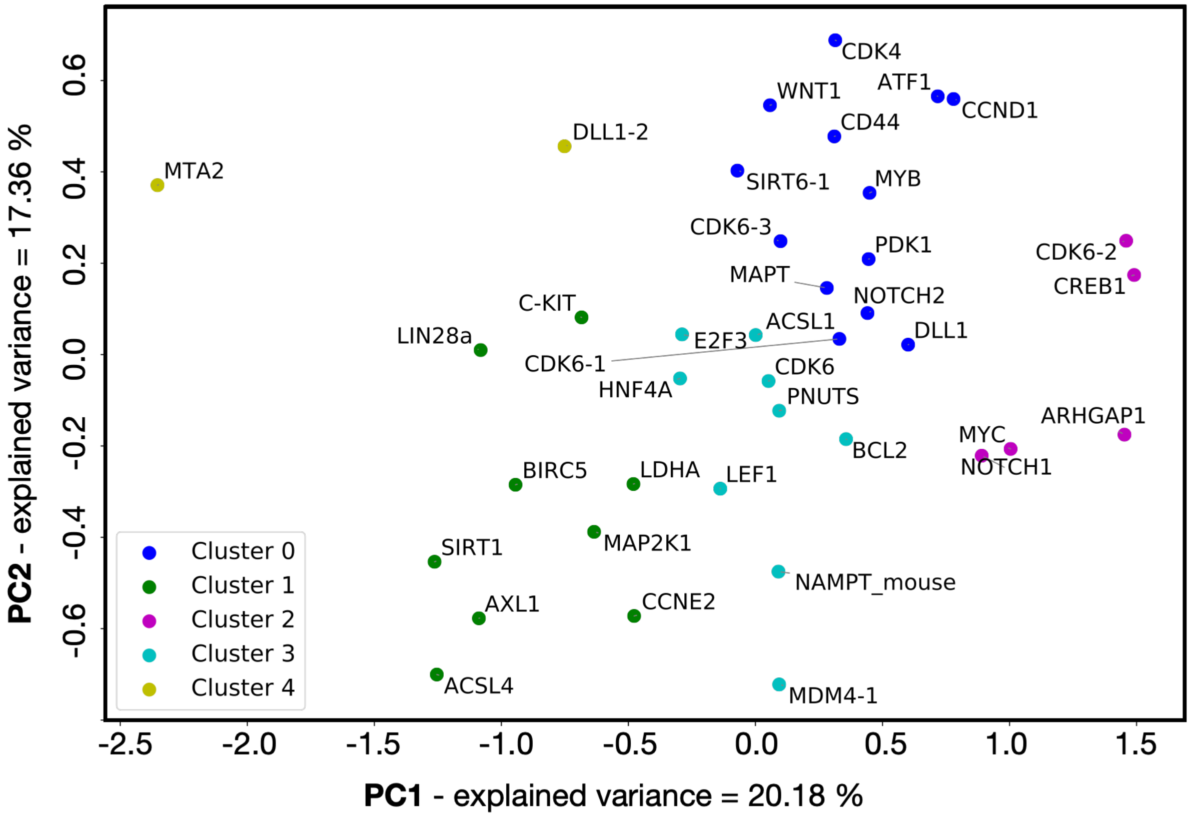


**Figure S1. Selection of *miR-34a* targets for analysis**. Principle Component Analysis (PCA) with first and second principle components (PC1 and PC2), followed by K-means clustering of various *miR-34a* mRNA targets. 12 targets were chosen for this study to represent a diversity in sequence, function, and predicted structural features.

**Table S1:** Sequences of all RNAs used in the study.

| Name | Sequence (uppercase = RNA, lowercase = DNA, m: 2´-O-methylation, phos = 5‘ phosphate) | Length (nt) |
|---|---|---|
| *miR-34a* guide | 5’- phos - UGGCAGUGUCUUAGCUGGUUGU -3’ | 22 |
| *miR-34a*-passenger | 5’- CAAUCAGCAAGUAUACUGCCCU-3’ | 22 |
| mRNA-*BCL2* | 5’- UCGAAUCAGCUAUUUACUGCCA -3’ | 22 |
| mRNA-*CCND1* | 5’- AGGAUCAGUUUACAAUGUCAUAUACUGCCA -3’ | 30 |
| mRNA-*CDK6* | 5’- UACUUUCUGCCACACACUGCCU -3’ | 22 |
| mRNA-*CD44* | 5’- UAGGCCACUAUGUGUUGUUACUGCCA -3’ | 26 |
| mRNA-*DLL1* | 5’- CCGGCCGCCUGCGGCACUGCCU -3’ | 22 |
| mRNA- *HNF4α* | 5’- AGGGCCACAUCCCACUGCCA -3’ | 20 |
| mRNA-*NOTCH1* | 5’- ACUUUUAUUUUACACAGAAACACUGCCU -3’ | 28 |
| mRNA-*NOTCH2* | 5’- UCAGUGAUGAGGAGGACAACACUGCCU -3’ | 27 |
| mRNA-*MTA2* | 5’- GGUACCUGGUUGGGGGAGGGGGGCGUGCACUGCCA -3’ | 35 |
| mRNA-*PNUTS* | 5’-AAAGUCACACUACAUGCACUGCCU -3’ | 24 |
| mRNA-*SIRT1* | 5’- ACACCCAGCUAGGACCAUUACUGCCA -3’ | 26 |
| mRNA-*WNT1* | 5’- GGAGACCCCUUGUUGCACUGCCC -3’ | 23 |
| PERFECT | 5’- ACAACCAGCUAAGACACUGCCA -3’ | 22 |
| Polyuridine | 5´- UUUUUUUUUUUUUUUUUUUUUU -3´ | 22 |
| allSCRAMBLED | 5’- UAUCAACAAAUUUUAUAAAAAU -3´ | 22 |
| seedSCRAMBLED | 5’- ACAACCAGCUAUCACACACGGA -3’ | 22 |
| AGO2 slicing probe | 5’-GGGCAACACAACCAGCUAAGACACUGCCAACACC -Cy3 | 34 |
| *CIS scaffold (all targets)* | 5'- GGGAGAACAAGCAGGGAGUACCUGC*AAACAACAAA*GGCG<br>CGGCGCUCAU**NNNNN**UUGUUCGCUU**UGGCAGUGUCUUAGC**<br>**UGGUUGU**UCUCGCGCCGCGCC*AAAACAAA*GGACGGAGUA<br>CGUCCAACAAAAGAAACAACAACAACAAC -3’ | |
| *TRANS scaffold* | 5’- GGGAGAACAAGCAGGGAGUACCUGC*AAACAACAAA*GGCG<br>CGCGUGCUGCCUCGCGGGCAAAACAAAAAACAA**NNNNN**AAA<br>ACAC*AAAACAAA*CCCGCGGGGCUGCGCGAGCGCC*AAAACAAA*<br>GGACGGAGUACGUCCAACAAAAGAAACAACAACAACAAC -3’ | |
| *TRANS scaffold WNT1 and CD44* | 5’- GGGAGAACAAGCAGGGAGUACCUGC*AAACAACAAA*GGCG<br>CGCGUGCUGCCUCGCGGGCUACAUAUUCUUAUAUAA**NNNNN**<br>AUUUUAAACCCGCGGGGCUGCGCGAGCGCC*AAAACAAA*GGA<br>CGGAGUACGUCCAACAAAAGAAACAACAACAACAAC -3’ | |

* **NNNNN** = mRNA target sequence or *miR-34a* sequence
** Underlined = reference hairpins GCAGGGAGUACCUGC and GGACGGAGUACGUCC
*** Italicised = buffer nucleotides *AAACAACAAA* and *AAAACAAA*

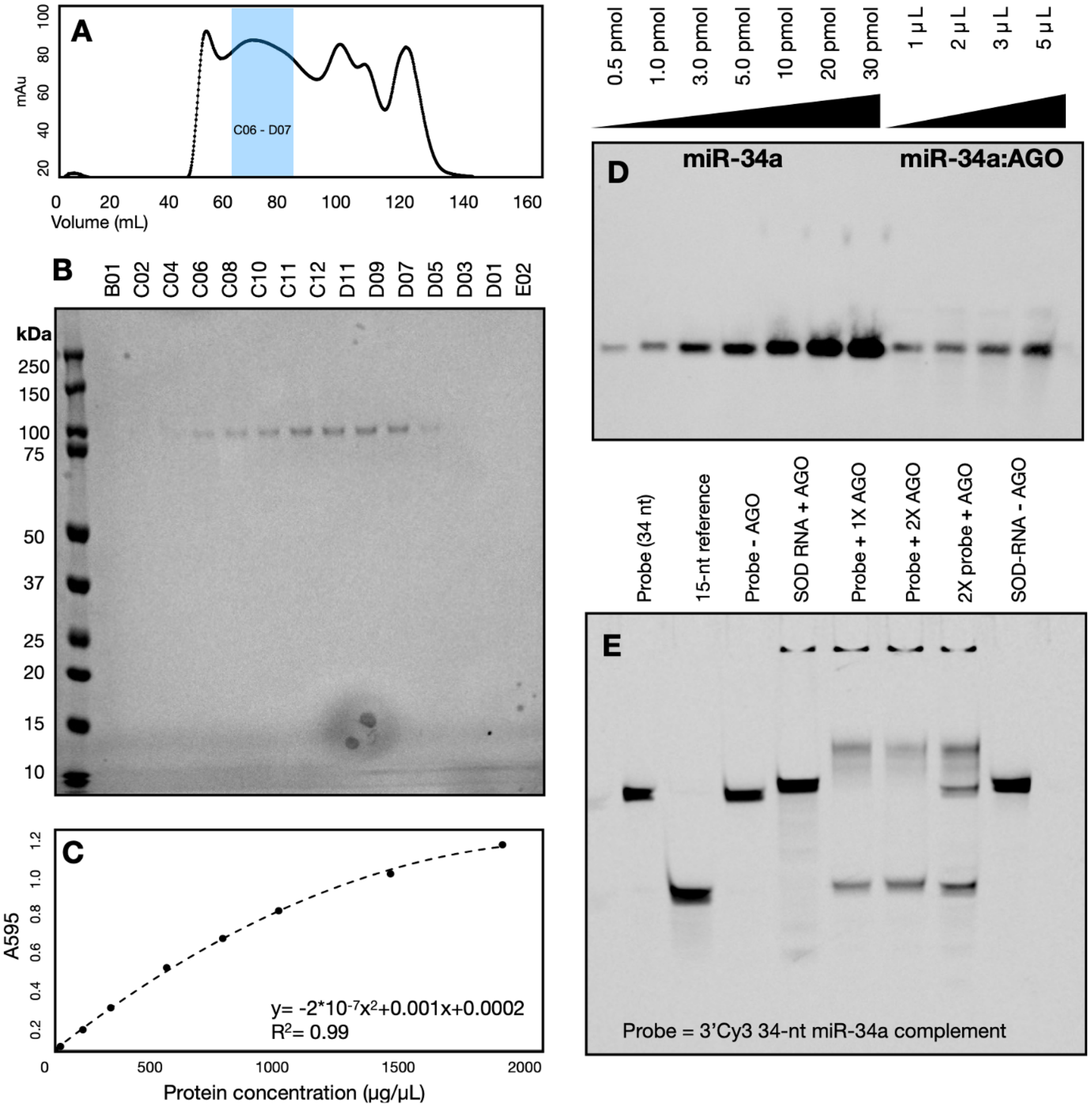


**Figure S2. AGO2 purification and quantification**, as reported by Banijamali et al. (2022). (A) Size exclusion chromatogram of *miR-34a* loaded AGO2 after reverse IMAC monitored at 280 nm wavelength. *miR-34a*-AGO2 eluted within the marked blue region (65 – 90 mL), indicating selected fractions. Peaks at ~ 100-120 ml represent unbound RNA. (B) Analysis by Bis-Tris NuPAGE (4-12%, 1.0 mm) of the SEC purified *miR-34a*-AGO2. To the left, molecular weight ladder in kDa, and AGO2 is visible near the 100 kDa band without any impurities. Lanes C06 to D07 were pooled and concentrated to ~ 2 ml. (C) Bradford assay revealed a 14.8 µM protein concentration based on the BSA standard curve fitted to a quadratic equation. (D) The *miR-34a* loading efficiency in AGO2 was assessed via northern blot with a complementary 3'Cy3-labelled *miR-34a* probe. Pure *miR-34a* (left) was used for standard, and *miR34a*:AGO2 was protease K digested following SEC purification and prior to loading on a 15% denaturing PAGE gel. Loading efficiency was estimated at 5%, and the protein was stored in SEC buffer + 5% glycerol at -80 °C. Note that this image is adapted from Banijamali et al. (2022) SI Figure S4b using the same procedure and sample. (E) The prepared *miR34a*:AGO2 is catalytically active, as shown by slicing assay. Following incubation of *miR34a*:AGO2 with a 34-nt 3'Cy3-labelled *miR34a* complementary probe for 1h at 37 °C, formation of a labelled 15-nt cleaved product was monitored by denaturing 20 % PAGE. The 3'Cy3-labelled SOD mRNA negative control did not cleave.

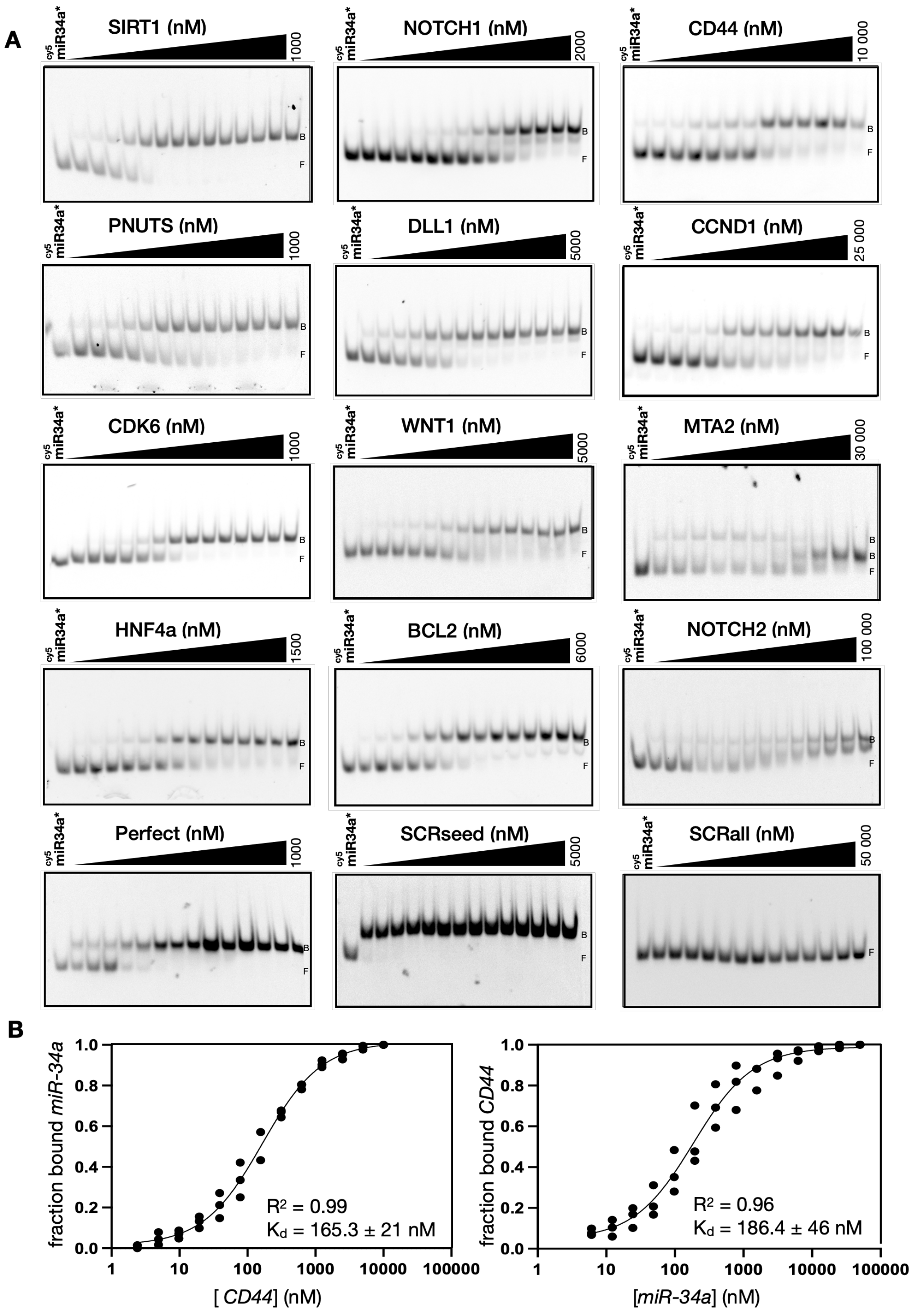


**Figure S3. Exemplary mRNA:miRNA (binary) binding assays by EMSA.** (A) Each mRNA target was titrated against 3'Cy5 labelled *miR-34a* and the fraction of *miR-34a* bound was monitored by fluorescence. 'B' denotes binary complex and 'F' denotes free cy5-*miR-34a*. *MTA2* was observed to have two unique *miR-34a*-bound bands, indicating that it adopts a unique conformation when bound to *miR-34a* at higher concentrations, favouring a more tightly folded structure, leading to faster migration. For *NOTCH2*, *miR-34a* migrated towards the upper bound position, yet it did not fully vanish as anticipated. This may be attributed to weak and thus incomplete binding. Triplicates were measured and fitted for each mRNA target. Here a single representative gel image is shown. Data for all individual replicates are presented in Supplementary File 2. (B) Fluorophore placement does not affect observed $K_{D,app}$. In the direct *CD44* binding assay (left), 3'Cy5 fluorophore was located on *miR-34a*, and unlabelled *CD44* was titrated against a constant concentration of *miR-34a*. In the reverse assay (right), 5'Cy5 fluorophore was located on *CD44*, and unlabelled *miR-34a* was titrated against constant *CD44*. Similar $K_{D,app}$ values were observed within error, providing evidence that the fluorophore itself did not significantly interfere with the binding process for this control target.

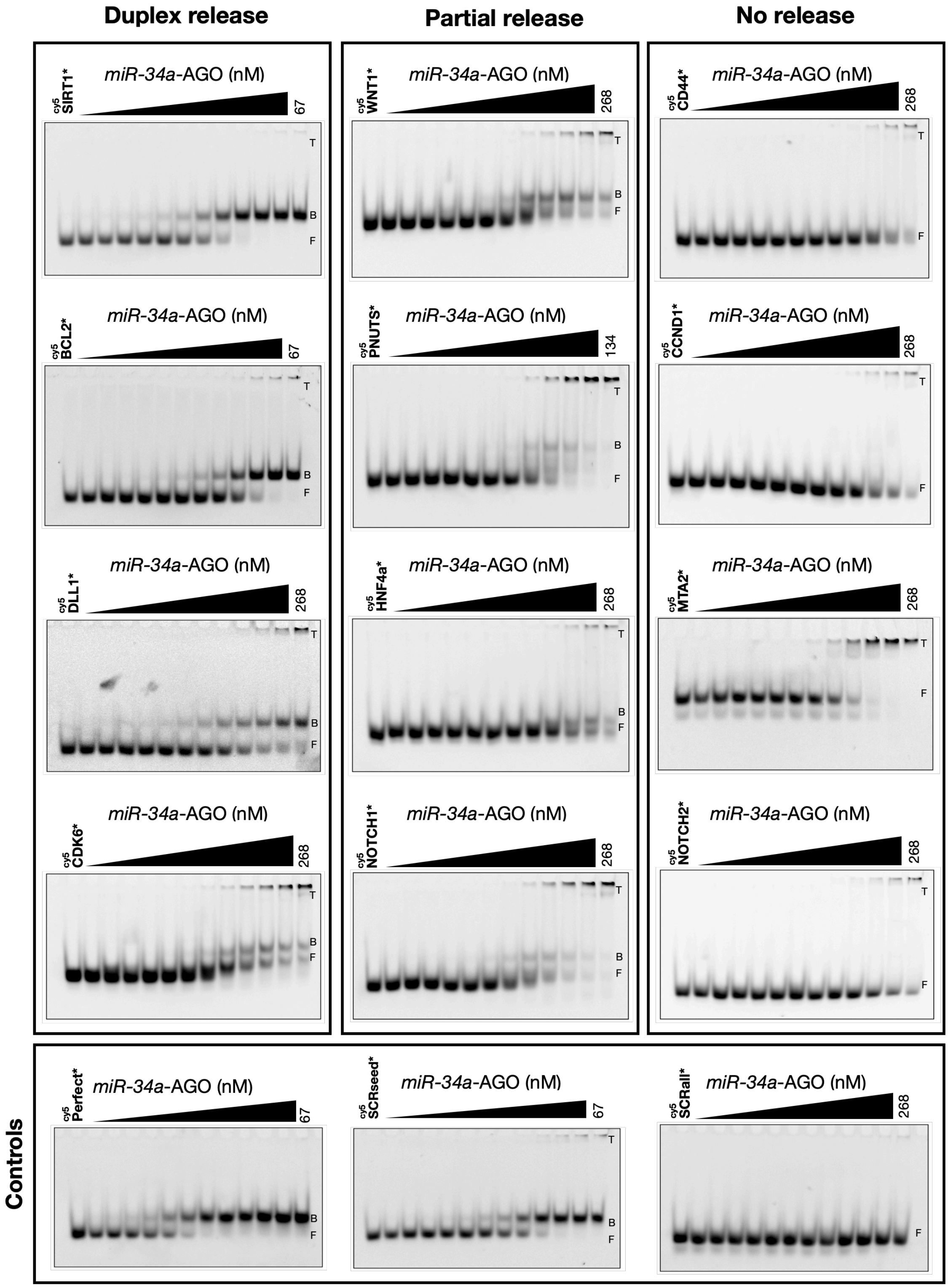


**Figure S4. Exemplary mRNA:miRNA-AGO2 (ternary) binding affinity assessed by EMSA.** One representative gel replicate for each target is shown. Target mRNAs are labelled with 5'Cy5 are indicated by an asterisk (*). The free mRNA is marked with F, the binary complex with B, and ternary complex with T. Cy5 signal is monitored by fluorescence at 647 nm. Data for all individual replicates are presented in Supplementary File 3. See quantification in Figure S5A.

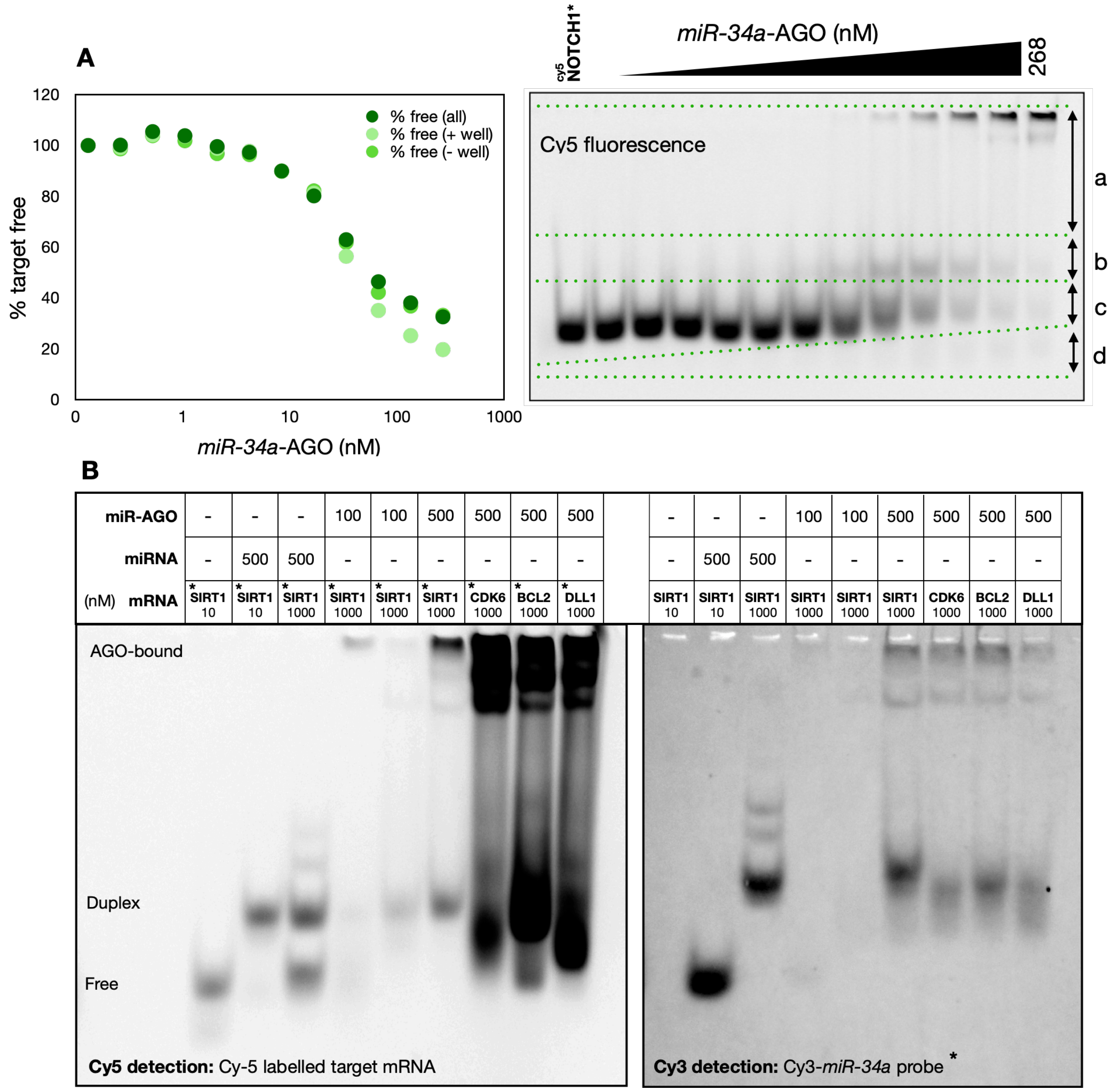


**Figure S5. mRNA:miRNA-AGO2 EMSA quantification.** Binding was assessed my monitoring disappearance of the free probe (Cy5-labelled target mRNA) upon titration of *miR-34a*-AGO2. The amount of free probe was plotted against increasing *miR-34a*-AGO2 concentration and fitted to a sigmoidal binding curve. Several quantification approaches were tested to find the most consistent method. As the background signal differed across each gel, inclusion of the entire lane length (a+b+c+d) was found to produce a consistently sigmoidal shape. Shown above, the trace '% free (all)' values were calculated by (c + d)/(a + b + c + d), i.e. tracking the amount of free target over the total amount of target per lane. The values were normalised to the control lane, here containing Cy5-*NOTCH1* to represent 100 % unbound. The '% free (all)' calculation was used throughout our dataset.
The '% free (+ well)' trace was calculated as (c)/(a + b + c), i.e. including the fluorescent signal in the well (upper) but not including the shadow below. The '% free (- well)' trace was calculated as (c)/(b + c). We chose to include all fluorescent signal observed on the gel in our analysis and found that none of the methods tested significantly altered the observed $K_{D,app}$.
(B) Northern blot of *miR-34a*-AGO2 binding to *SIRT1*, *CDK6*, *BCL2* and *DLL1* using a Cy3-labelled *miR-34a* probe (right), and corresponding EMSA before membrane transfer imaged with a Cy5 filter (left). Various ratios of Cy-5 labelled target mRNA, unlabelled *miR-34a*, and *miR34a*-loaded AGO2 were annealed with slow-cooling and assessed via EMSA (native 10 % polyacrylamide gel in EMSA buffer). asterisk (*) represents the labelled entity being traced. The samples were then transferred to a nylon+ membrane and detected via Northern Blotting using a Cy3-labelled probe to detect *miR-34a*. *miR-34a* was detected in the free form, the duplex, and the uppermost AGO2-bound bands, indicating that *miR-34a* dissociates from AGO2 to form a mRNA:*miR-34a* duplex.

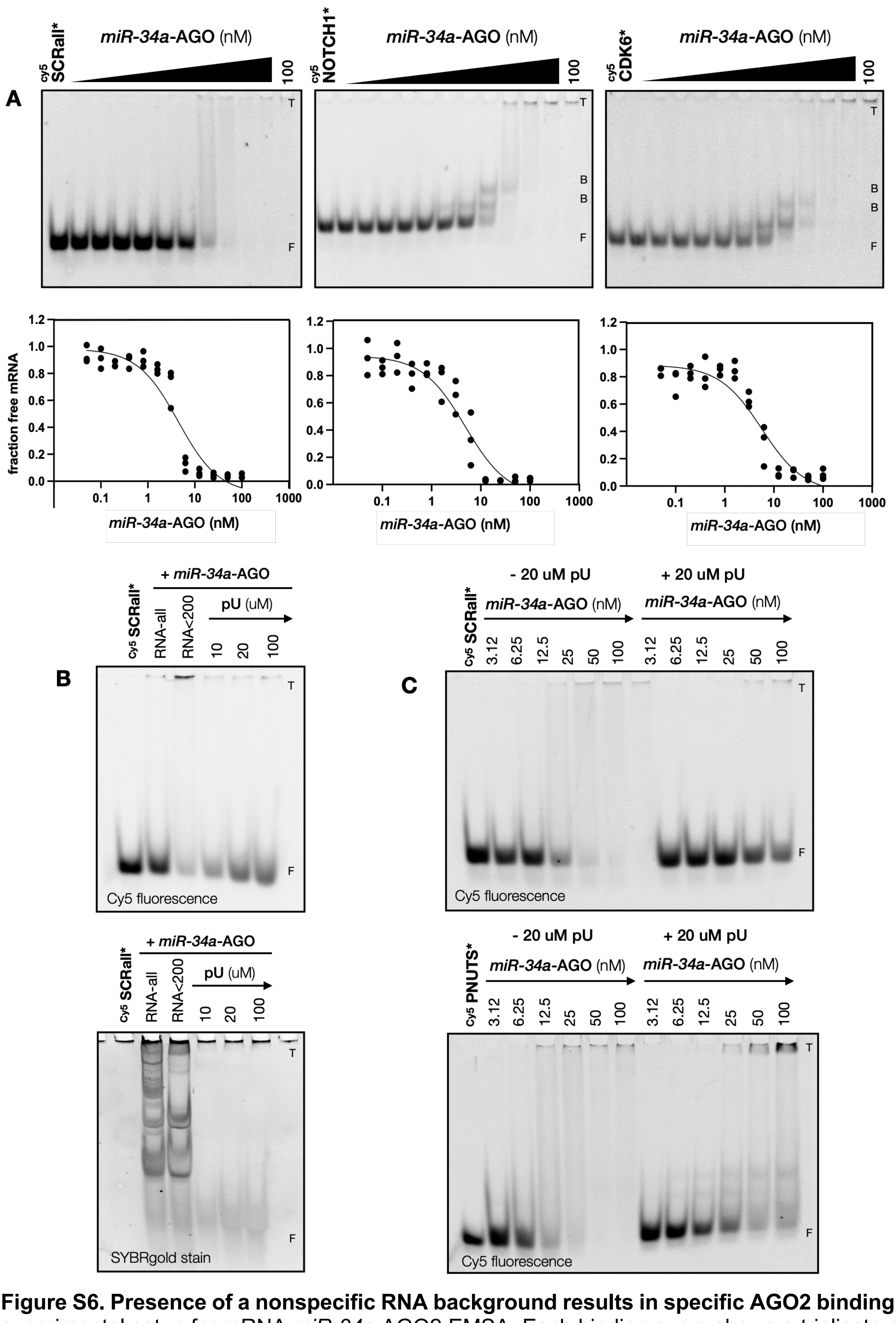


**Figure S6. Presence of a nonspecific RNA background results in specific AGO2 binding** . (A) Original experimental setup for mRNA:*miR-34a*-AGO2 EMSA. Each binding curve shows a triplicate measurement for the target mRNA (*PNUTS*, *WNT1*, or *NOTCH1*). Each target was observed to bind at approximately 5 nM [*miR-34a*-AGO2], with a rapid decrease in free target at that point, indicating the presence of non-specific binding. (B) Cy5-labelled SCRall (negative control) was equilibrated with *miR-34a*-AGO2 (100 nM) in EMSA buffer supplemented with either 4000 ng total RNA extract (RNA-all), 4000 ng RNA extract smaller than 200 kDa (RNA<200), or pU in increasing concentrations as labelled. Imaged via Cy5 fluorescence (top) or SYBRgold staining (lower). (C) Titrations with increasing *miR-34a*-AGO2 concentration against Cy5-labelled SCRall (top) or *PNUTS* (lower) comparing the absence and presence of 20 µM pU during equilibration. Target mRNAs labelled with 5'Cy5 are indicated by an asterisk (*). F indicates the free Cy5-labelled mRNA probe, B indicates binary complex, and T indicates ternary complex. All final mRNA:*miR-34a*-AGO2 EMSAs were carried out in the presence of 20 µM pU.

**Table S2.** Apparent relative binding affinities ($K_{D,app}$) and efficacy (monitored by luciferase activity) of mRNA targets. Binding affinities are listed for interaction of each target with both *miR-34a* and *miR-34a*-AGO2. Targets are sorted by ascending luciferase mean R/F.

| Target | mRNA:*miR-34a* | | mRNA:*miR-34a*-AGO2 | | Luciferase activity | |
|---|---|---|---|---|---|---|
| | $K_{D,app}$ (nM) | SD | $K_{D,app}$ (nM) | SD | Mean R/F | SD |
| PERFECT | 0.91 | 2.0 | 1.03 | 0.46 | 0.135 | 0.003 |
| BCL2 | 10.8 | 1.2 | 8.9 | 1.6 | 0.270 | 0.012 |
| NOTCH1 | 70.21 | 13.4 | 23.5 | 3.6 | 0.325 | 0.019 |
| CDK6 | 4.55 | 1.2 | 26.9 | 10.7 | 0.375 | 0.053 |
| CCND1 | 529.3 | 156.6 | 272 | 58 | 0.423 | 0.036 |
| SIRT1 | 0.41 | 0.1 | 2.7 | 0.3 | 0.430 | 0.043 |
| MTA2 | 944.3 | 274.5 | 66.1 | 11.2 | 0.457 | 0.007 |
| HNF4α | 37.6 | 6.3 | 38.9 | 20.2 | 0.467 | 0.041 |
| CD44 | 165.3 | 21.3 | 289 | 165 | 0.511 | 0.031 |
| NOTCH2 | 6055 | 3362 | 496 | 129 | 0.590 | 0.057 |
| PNUTS | 2.41 | 0.8 | 14.5 | 2.9 | 0.634 | 0.055 |
| WNT1 | 57.2 | 6.9 | 25 | 6.6 | 0.676 | 0.061 |
| DLL1 | 10.6 | 1.3 | 68.1 | 14.1 | 0.922 | 0.064 |
| SCRall | - | - | - | - | 1.094 | 0.014 |
| SCRseed | - | - | 3.5 | 0.11 | 1.094 | 0.077 |

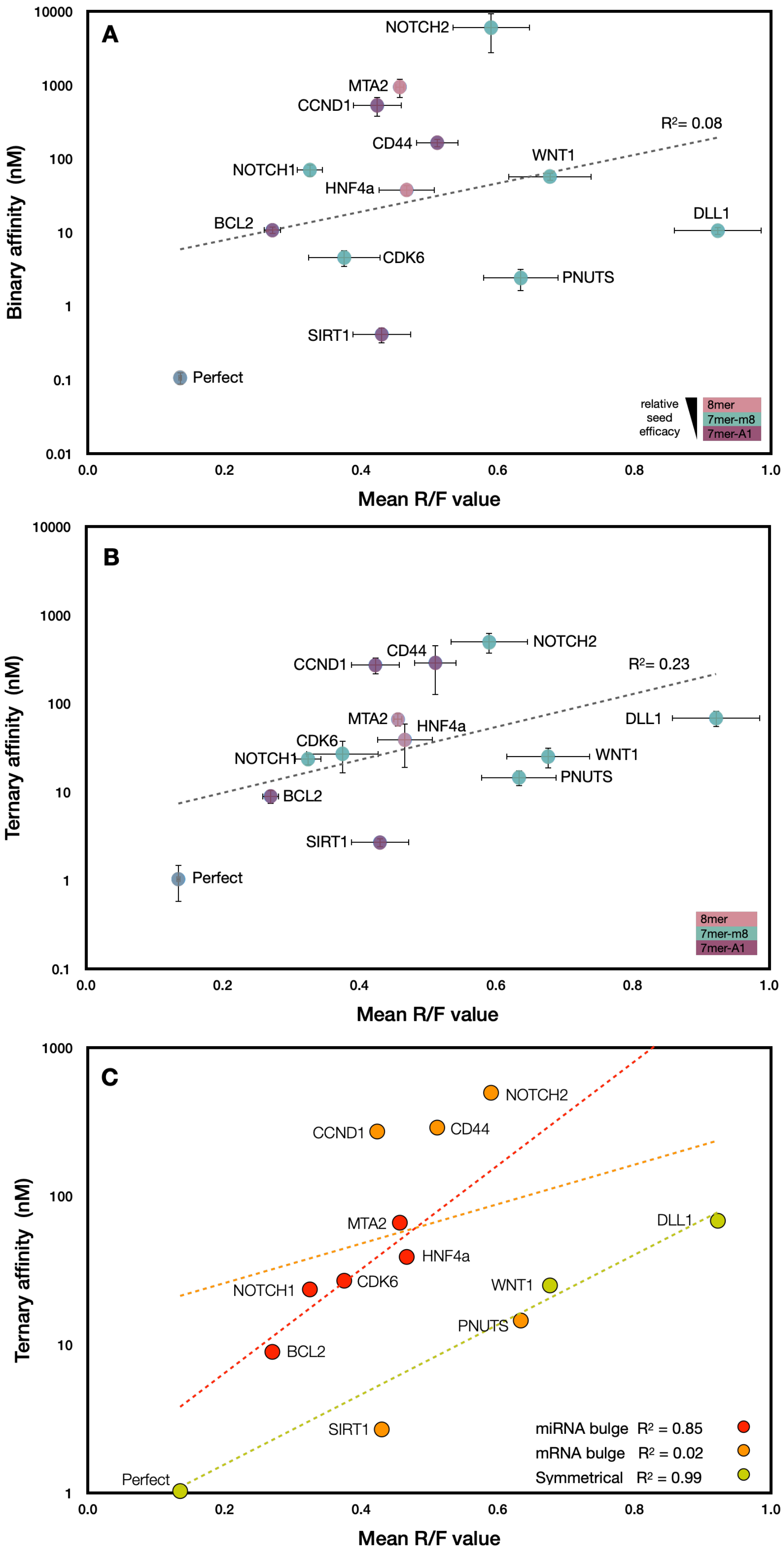


**Figure S7. Relative affinities of the 12 *miR-34a* targets versus their relative repression efficiency**. (A) Binary (mRNA:*miR-34a*, $R^2$ = 0.08) or (B) ternary (mRNA:*miR-34a*-AGO2, $R^2$ = 0.23) affinity as measured by EMSA vs repression measured by luciferase reporter assays. The mRNA targets are coloured by seed type where 8mer is orange, 7mer-m8 is purple, and 7mer-A1 is cyan. (C) Ternary affinity versus repression, grouped according to structural class, where miRNA bulge is red ($R^2$ = 0.85), mRNA bulge is orange ($R^2$ = 0.02), and symmetrical is yellow ($R^2$ = 0.99).

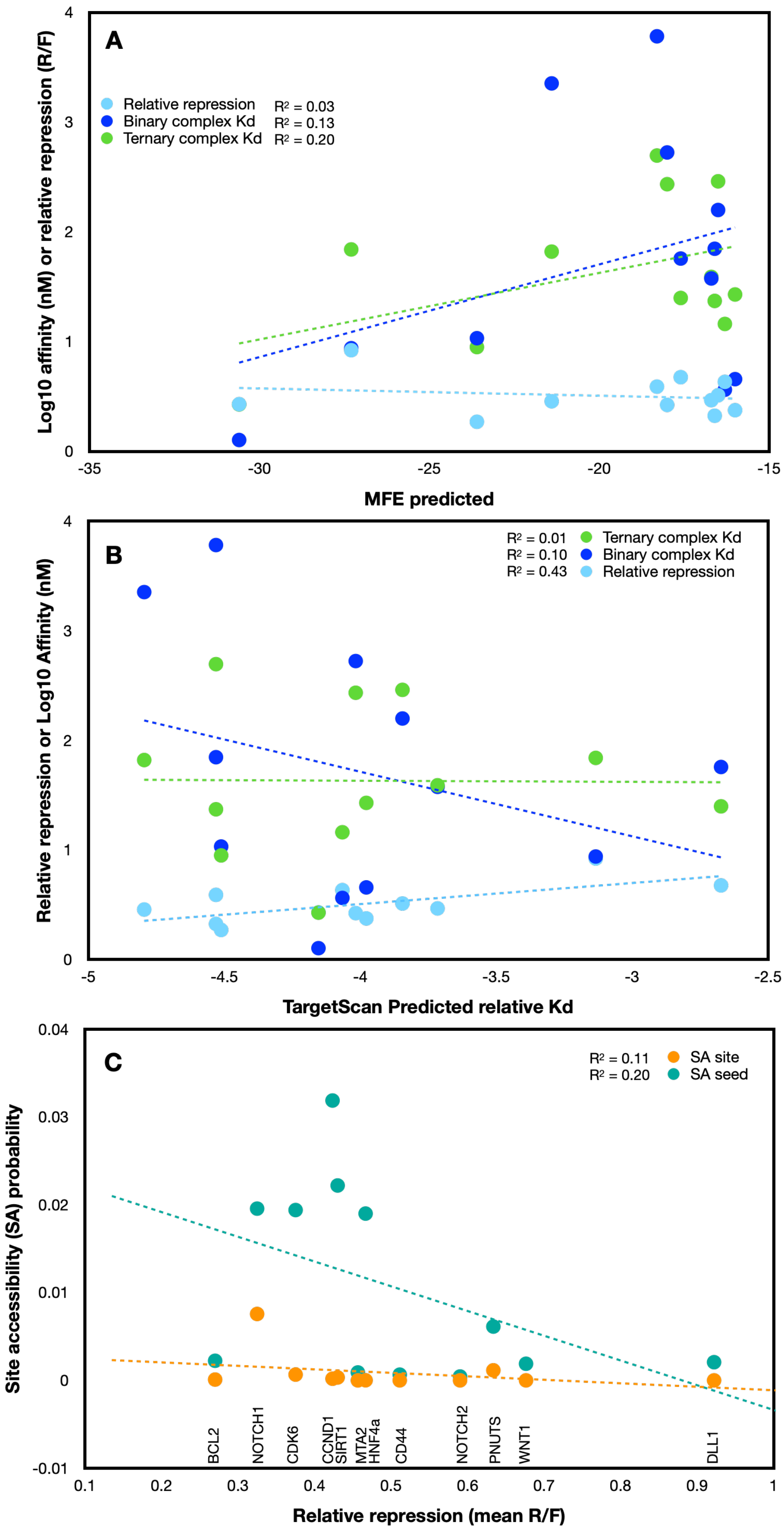


**Figure S8. Comparison of experimental results with bioinformatically predicted scores.** (A) Minimum Free Energy (MFE) of the predicted lowest free energy secondary structures for each target was computed using RNAcofold, binary and ternary $K_{D,app}$ values were obtained via EMSA, and relative repression scores were obtained via luciferase assay. (B) Comparison of predicted relative $K_D$ (obtained from TargetScan database) with $K_{D,app}$ obtained via EMSA, and relative repression. (C) Comparison of site accessibility (SA) of the luciferase constructs (assessed by RNAplfold) with relative repression. Binary complex denotes mRNA:*miR-34a* and ternary denotes mRNA:*miR34a*-AGO2.

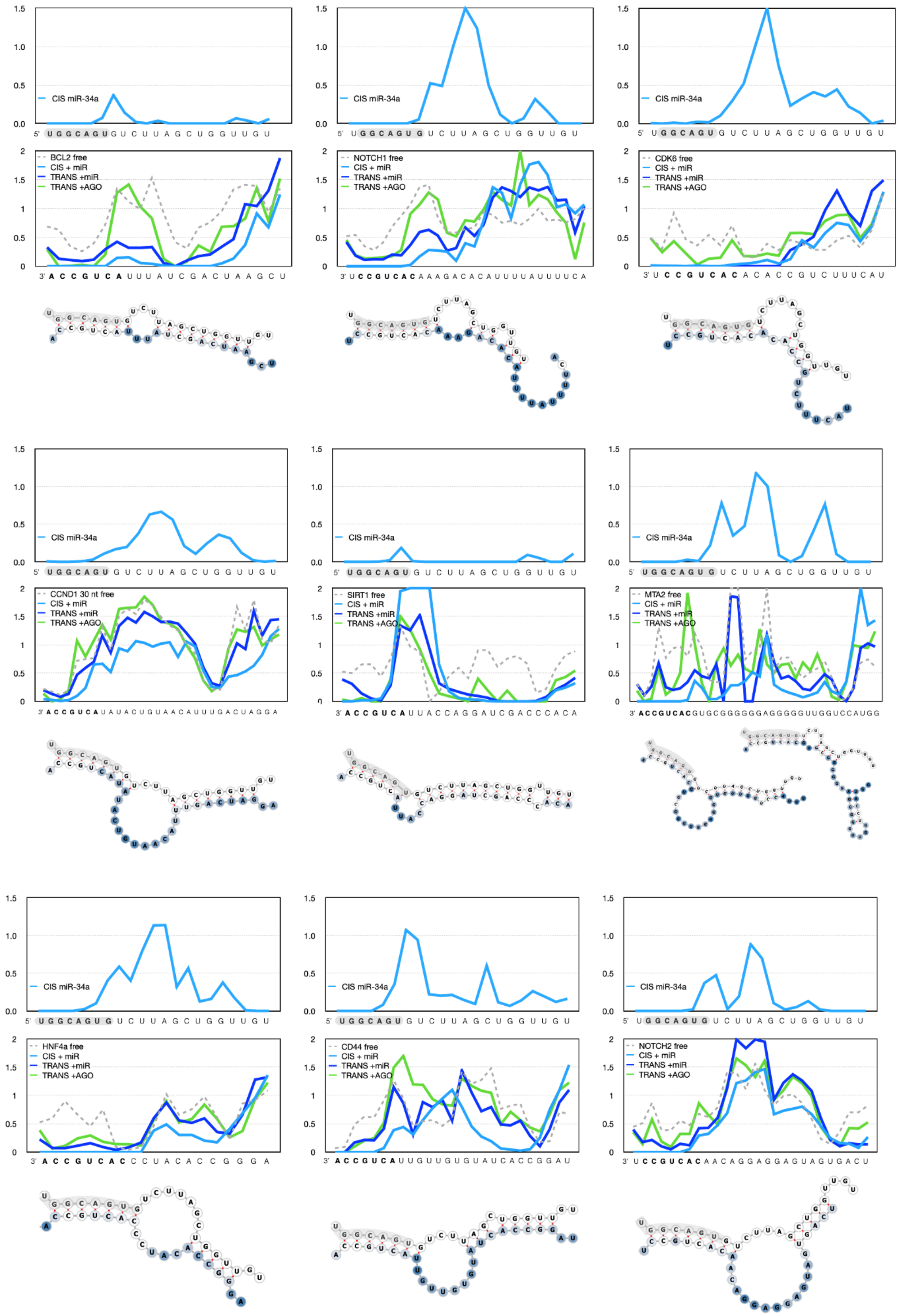

CIS miR-34a
BCL2 free
CIS + miR
TRANS +miR
TRANS +AGO
NOTCH1 free
CDK6 free
CCND1 30 nt free
SIRT1 free
MTA2 free
HNF4a free
CD44 free
NOTCH2 free

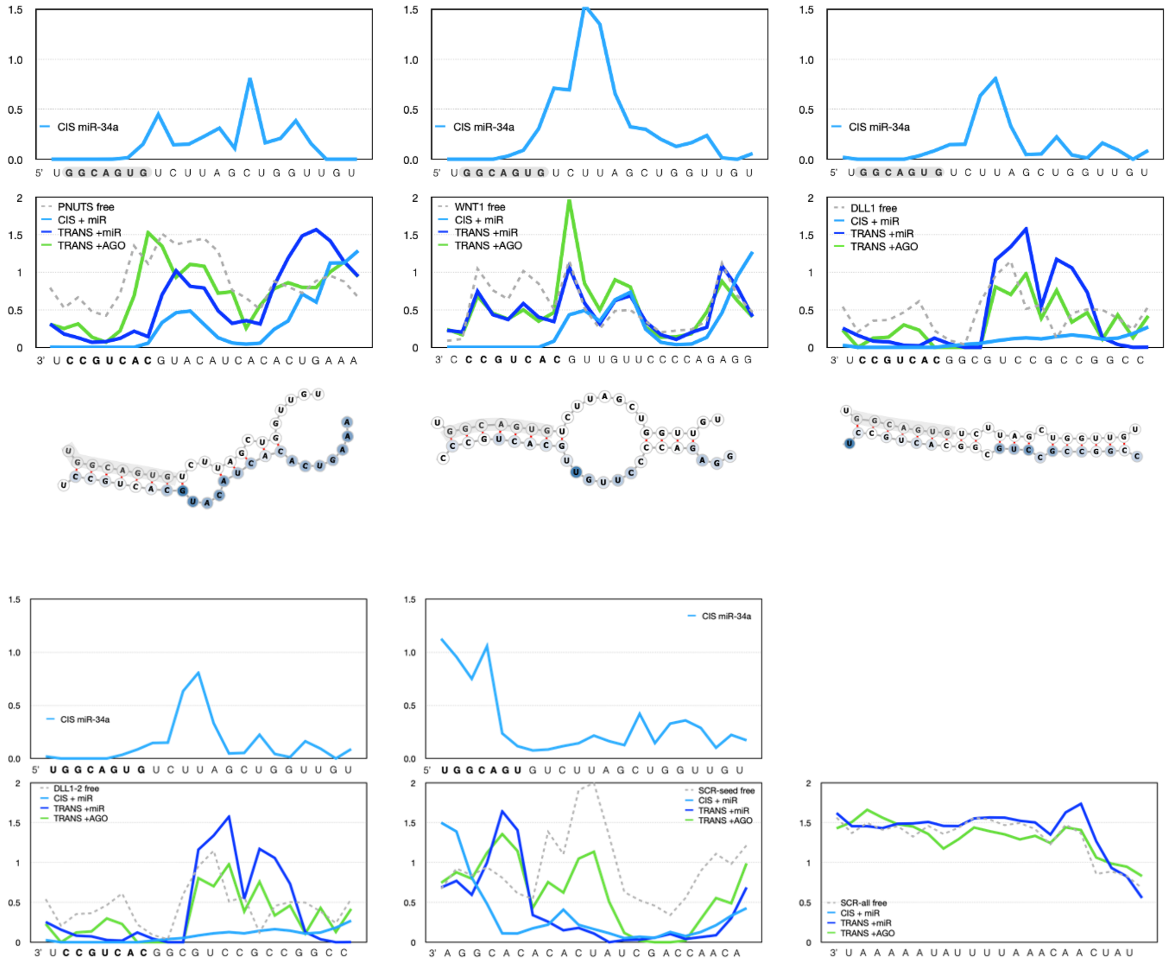


**Figure S9. RABS reactivity profiles of the 12 *miR-34a* targets.** CIS reactivity profiles, where *miR-34a* is bound to its mRNA target site within a single RNA (light blue), and TRANS reactivity profiles where the mRNA target site within an RNA scaffold and is bound to either *miR-34a* (dark blue) or *miR-34a*-AGO2 (green) are shown for each target. The predicted secondary structure, using a combination of both CIS and TRANS reactivity is shown below, with TRANS +*miR-34a*-AGO reactivity pattern shown on the structure. Seed regions are marked in bold and grey highlight. Structures were generated using RNAforna. Two plausible structures for *MTA2* were predicted, consistent with the two bands seen via native gel (Figure S3). Values larger than 0.5 indicates high reactivity (i.e., no involvement in stable base pairing). Note that the range in reactivity values is greater on the mRNA than the miRNA, although we are unable to interpret meaningful differences in values significantly greater than 0.5***.***

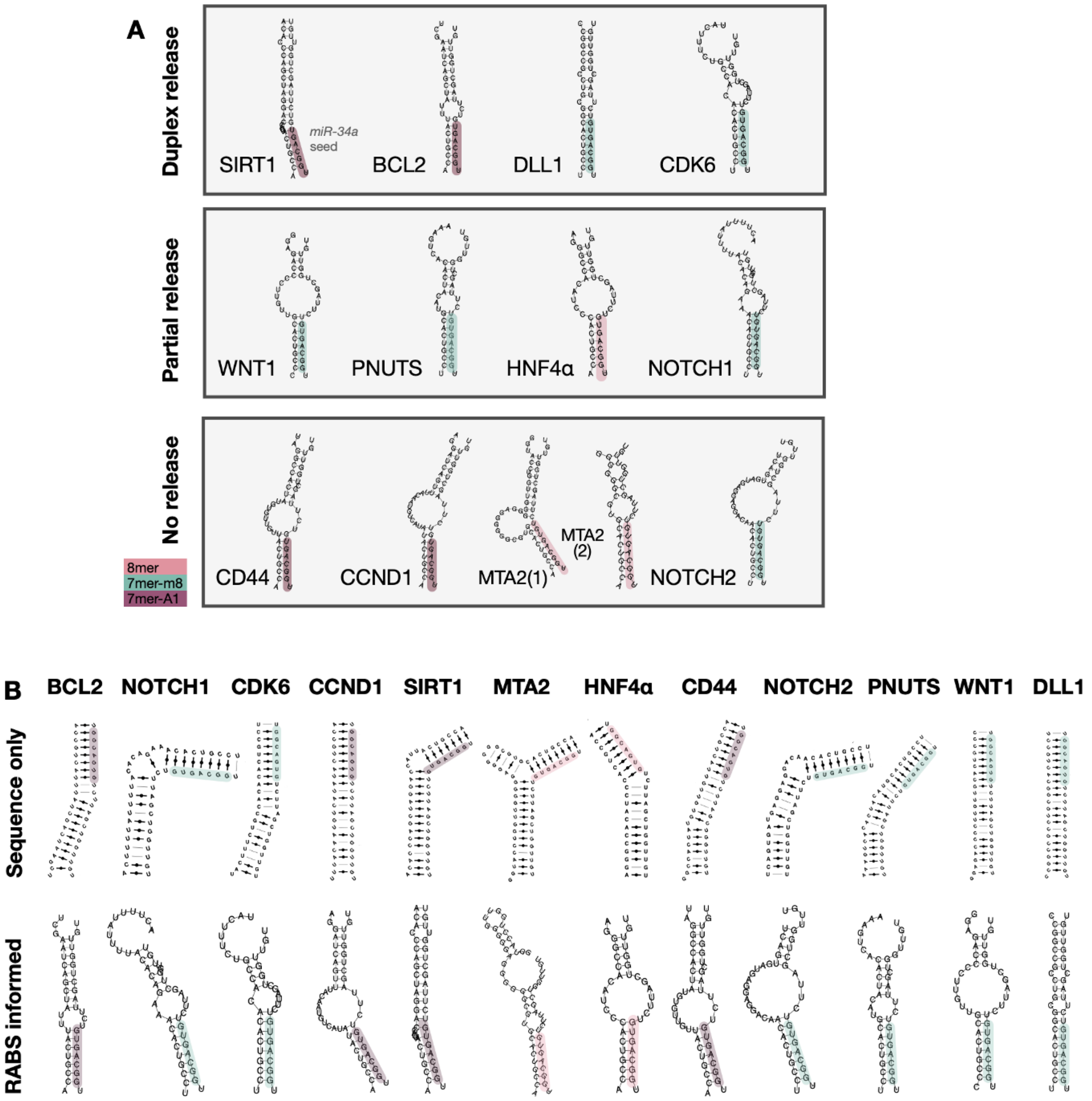


**Figure S10. Assessment of structural parameters relating to duplex release and prediction accuracy.** (A) mRNA:*miR-34a* duplex structures grouped by miRNA duplex release. Titration of *miR-34a*-AGO resulted in formation of an intermediate band (an mRNA:*miR-34a* duplex) in the case of *SIRT1*, *BCL2*, *DLL1*, *CDK6*. *WNT1*, *PNUTS*, *HNF4α,* and *NOTCH1* produced fainter intermediate bands and were thus classified as partial release. No release was observed for *CD44*, *CCND1*, *MTA2* and *NOTCH2* (Figure S4). Notably, the targets which did not exhibit duplex release all had mRNA bulge structures with large bulge sizes (>4 nucleotides of asymmetrical base pairing). Note that *MTA2* is exhibited two distinct conformations in solution, as previously discussed and exemplified in Figure S3 and S9. No correlation with seed type was observed in the dataset. (B) Advanced secondary structure prediction using sequence information only (upper) and RABS reactivity data (lower). The sequence-only structures were predicted using MCfold and include all possible non-WC pairs. The RABS-informed structures were derived using a combination of sequence information and reactivity input from both CIS and TRANS RABS experiments and were calculated via RNAprobing. Construct length was optimised via EMSA and RABS; if the construct was not observed to bind, varying mRNA lengths were tested (resulting in elongations for *CCND1* and *CD44*). Other constructs were observed to bind shorter target site lengths than predicted (*NOTCH1*, *CDK6*, *MTA2* and *PNUTS*). This result highlights that it is difficult to predict miRNA binding sites via sequence alone. The *miR-34a* seed is highlighted according to seed type.

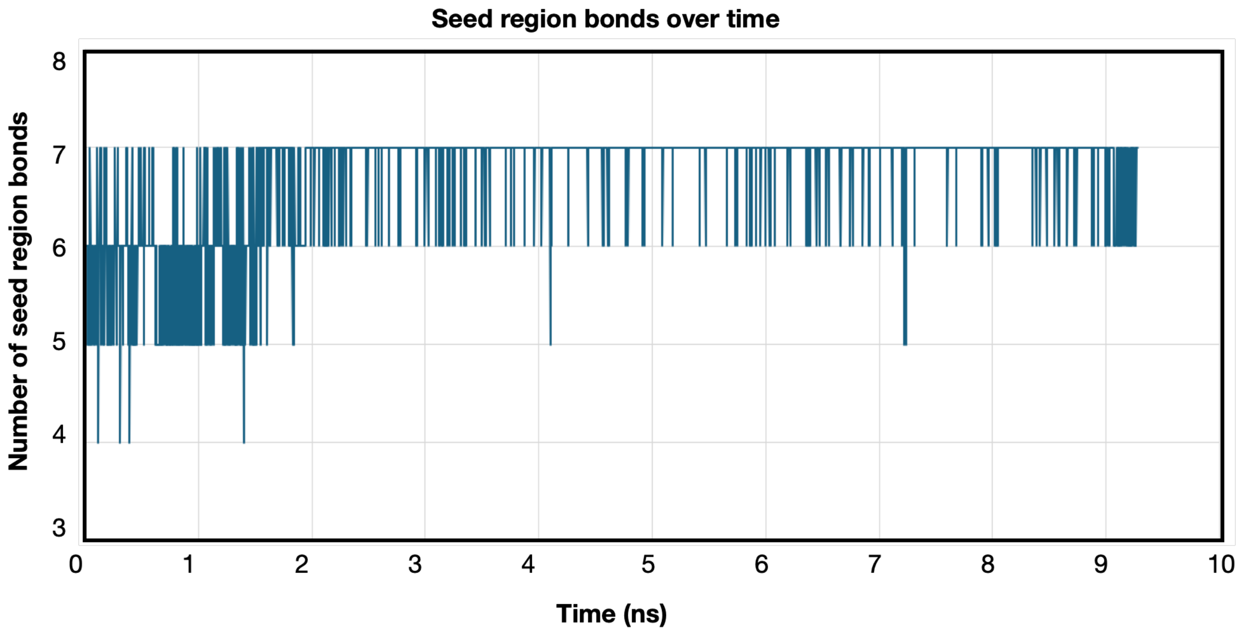


**Figure S11. Seed region contacts between the mRNA:*miR-34a* duplex and AGO2 during MD simulation.** Contacts were defined as <0.35 nm from RNA backbone to protein and were tracked over the 10 ns simulation time. Contacts to all 7 seed bases formed within 2 ns and were stable throughout the simulation.
