## Supplementary File 1 for "Biophysics of microRNA-34a targeting and its influence on down-regulation": analysis.py

```
#!/usr/bin/env python
# -*- coding: utf-8 -*-

import re
import numpy as np
import scipy as sp
import pandas as pd
from scipy.cluster.vq import kmeans2
# from sklearn.cluster import KMeans
import matplotlib.pyplot as plt
from matplotlib import colors as mcolors
# import networkx as nx
from mpl_toolkits.mplot3d import Axes3D
from mpl_toolkits.mplot3d import proj3d
from matplotlib.patches import FancyArrowPatch
from mpl_toolkits.mplot3d.proj3d import proj_transform
from matplotlib.text import Annotation
# from mpl_toolkits.mplot3d.art3d import Line3DCollection
# from adjustText import adjust_text
from sklearn.feature_selection import VarianceThreshold
import operator

# plt.style.use('ggplot')


colors = dict(mcolors.BASE_COLORS, **mcolors.CSS4_COLORS)


class objdict(dict):

    def __getattr__(self, name):
        if name in self:
            return self[name]
        else:
            raise AttributeError("No such attribute: " + name)

    def __setattr__(self, name, value):
        self[name] = value

    def __delattr__(self, name):
        if name in self:
            del self[name]
        else:
            raise AttributeError("No such attribute: " + name)


Colors = objdict(colors)


# def repel_labels(ax, x, y, z, labels, k=0.01):
#     G = nx.DiGraph()
#     data_nodes = []
#     init_pos = {}
#     for xi, yi, zi, label in zip(x, y, z, labels):
#         data_str = 'data_{0}'.format(label)
#         G.add_node(data_str)
#         G.add_node(label)
#         G.add_edge(label, data_str)
#         data_nodes.append(data_str)
#         init_pos[data_str] = (xi, yi, zi)
#         init_pos[label] = (xi, yi, zi)
#
#     pos = nx.spring_layout(G, dim=3, pos=init_pos, fixed=data_nodes, k=k)
#
#     # undo spring_layout's rescaling
#     pos_after = np.vstack([pos[d] for d in data_nodes])
#     pos_before = np.vstack([init_pos[d] for d in data_nodes])
#     scale_x, shift_x = np.polyfit(pos_after[:, 0], pos_before[:, 0], 1)
#     scale_y, shift_y = np.polyfit(pos_after[:, 1], pos_before[:, 1], 1)
#     scale_z, shift_z = np.polyfit(pos_after[:, 2], pos_before[:, 2], 1)
#     shift = np.array([shift_x, shift_y, shift_z])
#     scale = np.array([scale_x, scale_y, scale_z])
#     for key, val in pos.items():
#         pos[key] = (val * scale) + shift
#
#     for label, data_str in G.edges():
#         # ax.text3D(Xred[i, 0], Xred[i, 1], Xred[i, 2], label, zdir='x')
#         annotate3D(ax, s=label,
#                     xyz=pos[data_str],
# 					xycoords='data',
#                     xytext=pos[label][:2],
# 					textcoords='data',
# 					ha='right',
# 					va='bottom',
#                     arrowprops=dict(arrowstyle="->",
#                                     shrinkA=0, shrinkB=0,
#                                     connectionstyle="arc3",
#                                     color=Colors.gray),
# 					)
#     # expand limits
#     all_pos = np.vstack(pos.values())
#     x_span, y_span, z_span = np.ptp(all_pos, axis=0)
#     mins = np.min(all_pos - x_span * 0.15, 0)
#     maxs = np.max(all_pos + y_span * 0.15, 0)
#     ax.set_xlim([mins[0], maxs[0]])
#     ax.set_ylim([mins[1], maxs[1]])
#     ax.set_zlim([mins[2], maxs[2]])


class Annotation3D(Annotation):
    '''Annotate the point xyz with text s'''

    def __init__(self, s, xyz, *args, **kwargs):
        Annotation.__init__(self, s, xy=(0, 0), *args, **kwargs)
        self._verts3d = xyz

    def draw(self, renderer):
        xs3d, ys3d, zs3d = self._verts3d
        xs, ys, zs = proj_transform(xs3d, ys3d, zs3d, renderer.M)
        self.xy = (xs, ys)
        Annotation.draw(self, renderer)


def annotate3D(ax, s, *args, **kwargs):
    '''add anotation text s to to Axes3d ax'''

    tag = Annotation3D(s, *args, **kwargs)
    ax.add_artist(tag)


def dim_red_pca(X, d=0, corr=False, norm=None):
    r"""
    Performs principal component analysis.

    Parameters
    ----------
    X : array, (n, d)
        Original observations (n observations, d features)

    d : int
        Number of principal components (default is ``0`` => all components).

    corr : bool
        If true, the PCA is performed based on the correlation matrix.

    norm : None or string
        Possible values: mean_centre, z_score, min_max

    Notes
    -----
    Always all eigenvalues and eigenvectors are returned,
    independently of the desired number of components ``d``.

    Returns
    -------
    Xred : array, (n, m or d)
        Reduced data matrix

    e_values : array, (m)
        The eigenvalues, sorted in descending manner.

    e_vectors : array, (n, m)
        The eigenvectors, sorted corresponding to eigenvalues.

    """
    # Normalise
    if norm is None:
        X_ = X
    elif norm == 'mean_centre':
        pass
        # Each feature will now have mean ~= 0
        X_ = X - X.mean(0)
    elif norm == 'z_score':
        # Each feature will now have mean ~= 0 and std ~= 1
        X_ = (X - X.mean(0)) / X.std(0)
    elif norm == 'min_max':
        # Each feature will now have min ~= 0 and max ~= 1
        X_ = (X - X.min(0)) / X.ptp(0)
    else:
        raise ValueError(("Please specify a valid normalisation procedure.\n"
                          "Possible values: mean_centre, z_score, min_max.\n"))
    print('Normalisation:\t{}'.format(norm))
    # Compute correlation / covarianz matrix
    if corr:
        CO = np.corrcoef(X_.T)
        print('Method:\t\t{}'.format('correlation'))
    else:
        CO = np.cov(X_.T)
        print('Method:\t\t{}'.format('covariance'))
    # Compute eigenvalues and eigenvectors
    e_values, e_vectors = sp.linalg.eigh(CO)

    # Sort the eigenvalues and the eigenvectors descending
    idx = np.argsort(e_values)[::-1]
    e_vectors = e_vectors[:, idx]
    e_values = e_values[idx]
    # Get the number of desired dimensions
    d_e_vecs = e_vectors
    if d > 0:
        d_e_vecs = e_vectors[:, :d]
    else:
        d = None
    # Map principal components to original data
    LIN = np.dot(d_e_vecs, np.dot(d_e_vecs.T, X_.T)).T
    return LIN[:, :d], e_values, e_vectors


def remove_correlated(data_frame, threshold):
    '''
    Given two or more features that are correlated with value higher than
    threshold, keep only the first. Returns a new data frame with correlated
    features removed
    '''
    corr = data_frame.corr()
    features = data_frame.columns.values[:]
    blacklisted = list()
    for f in features:
        if f not in blacklisted:
            correlated_features = [i for i in features if abs(corr.loc[f, i]) >= threshold]
            blacklisted += correlated_features[1:]
    whitelisted = [i for i in features if i not in blacklisted]
    return data_frame[whitelisted]


# Load and clean up data.
# IN_FILE = 'features'
IN_FILE = 'out'
df = pd.read_csv(IN_FILE, index_col=[0], sep=r"\s+", engine='python')
df.fillna(value=0.0, inplace=True)
# eliminate each and every constant feature.
# df_no_zero = df.loc[:, (df != 0).any(axis=0)]
# df = df_no_zero

# eliminate junciton features
# df = df[[i for i in df.columns if not i.startswith('junc')]]

# eliminate features with low variance:
sel = VarianceThreshold(threshold=(.9 * (1 - .9)))

SN_variance = sel.fit_transform(df.values)
df_cleaned = df[df.columns[sel.get_support()]]
eliminated_features = df.columns[map(operator.not_, sel.get_support())]
print('{} features eliminated due to low variance:'.format(len(eliminated_features)))
print('')
print('\n'.join(eliminated_features))
print('')

df_cleaned_cleaned = remove_correlated(df_cleaned, threshold=0.9)
df_cleaned = df_cleaned_cleaned

SN = df_cleaned.values

# PCA
# If corr=True, the normalisation has no effect on the evalues
Xred, evals, evecs = dim_red_pca(SN, 3, norm='z_score', corr=False)
print('First 4 evlues: {}'.format([round(i, 2) for i in evals[:4]]))

# All eigenvectors must be of unit norm.
for ev in evecs:
    np.testing.assert_array_almost_equal(1.0, np.linalg.norm(ev), decimal=6)

# Plot explained variance by different principal components.
tot = sum(evals)
var_exp = [(i / tot) * 100 for i in sorted(evals, reverse=True)]
cum_var_exp = np.cumsum(var_exp)

N_components = 10
objects = list(['PC %s' % i for i in range(1, N_components)])
y_pos = np.arange(len(objects))
height = var_exp[:N_components - 1]
plt.bar(y_pos, height, align='center', color=Colors.slateblue)
plt.plot(y_pos, cum_var_exp[:N_components - 1], marker='.', color=Colors.coral, label='cumulative')
plt.xticks(y_pos, objects)
plt.gca().xaxis.set_ticks_position('none')
plt.gca().yaxis.set_ticks_position('none')
plt.ylabel('Explained variance in percent')
plt.title('Explained variance by different principal components')
plt.ylim(0, 100)
plt.gca().yaxis.grid(True)
for spine in plt.gca().spines.values():
    spine.set_visible(False)
plt.legend()
plt.savefig('analysis_v4_explained_variance.pdf', ftype='pdf')
# plt.show()


# Clustering
clust, labels_ = kmeans2(data=Xred, k=5, iter=100, thresh=1.0000000000000001e-05, minit='random', missing='warn')
# quit()

# labels_ = [1, 4, 1, 4, 2, 2, 3, 2, 3, 0, 0, 1, 1, 1, 1, 2, 3, 1, 0, 2, 1, 1, 3, 4, 4, 1, 2, 3, 4, 1, 1, 1, 1, 4, 0, 3, 0]

# kmeans = KMeans(n_clusters=4, n_init=10, max_iter=500, tol=1.0000000000000001e-05, init='random')
# kmeans.fit(Xred)
# clusters = kmeans.cluster_centers_
# labels_ = kmeans.labels_

# determine variance portion
xlab = '1. PC - ExpVar = {:.2f} %'.format(evals[0] / sum(evals) * 100)
ylab = '2. PC - ExpVar = {:.2f} %'.format(evals[1] / sum(evals) * 100)
zlab = '3. PC - ExpVar = {:.2f} %'.format(evals[2] / sum(evals) * 100)

# Scatter plot with clustering of transformed data in 3D
mean_x = np.mean(Xred[:, 0])
mean_y = np.mean(Xred[:, 1])
mean_z = np.mean(Xred[:, 2])

print('Shape transformed: {}'.format(Xred.shape))


class Arrow3D(FancyArrowPatch):

    def __init__(self, xs, ys, zs, *args, **kwargs):
        FancyArrowPatch.__init__(self, (0, 0), (0, 0), *args, **kwargs)
        self._verts3d = xs, ys, zs

    def draw(self, renderer):
        xs3d, ys3d, zs3d = self._verts3d
        xs, ys, zs = proj3d.proj_transform(xs3d, ys3d, zs3d, renderer.M)
        self.set_positions((xs[0], ys[0]), (xs[1], ys[1]))
        FancyArrowPatch.draw(self, renderer)


plt.close()
fig = plt.figure(figsize=(15, 10))
ax = Axes3D(fig)
scatterHs = []
clr = ['b', 'g', 'm', 'c', 'y', 'darkviolet', 'k']
for cluster in set(labels_):
    scatterHs.append(ax.scatter(Xred[labels_ == cluster, 0], Xred[labels_ == cluster, 1], Xred[labels_ == cluster, 2], color=clr[cluster], label='Cluster {}'.format(cluster)))

scatter_labels = [k.split('.')[0] for k in df_cleaned.index.values]
for i, txt in enumerate(scatter_labels):
    ax.text3D(Xred[i, 0], Xred[i, 1], Xred[i, 2], txt, zdir='x')
    # repel_labels(ax, Xred[:, 0], Xred[:, 1], Xred[:, 2], labels=scatter_labels, k=0.1)
# for v in evecs.T[:3]:
#     a = Arrow3D([mean_x, v[0]], [mean_y, v[1]], [mean_z, v[2]], mutation_scale=10, lw=3, arrowstyle="-|>", color="r")
#     ax.add_artist(a)
ax.set_xlabel(xlab)
ax.set_ylabel(ylab)
ax.set_zlabel(zlab)
# ax.set_xlim(-2.5, 2.5)
# ax.set_ylim(-2.5, 2.5)
# ax.set_zlim(-2.5, 2.5)
plt.legend(handles=scatterHs, loc=1)
plt.title('First three principle components')
# ax.view_init(elev=10, azim=16)
ax.view_init(elev=19, azim=47)
# plt.savefig('analysis_v4_scatter.pdf', ftype='pdf', bbox_inches='tight', transparent=True)
plt.show()


def plot_evectors(list_evectors, x_tick_labels, title):
    '''
    Plot the eigenvectors for deriving the influence of each feature
    '''
    n_evectors = len(list_evectors)
    n_features = len(list_evectors[0])
    first = 0
    last = n_evectors - 1

    plt.close()
    fig, ax = plt.subplots(max(n_evectors, 2), 1, figsize=(35, 10))
    for i, vec in enumerate(list_evectors):
        ax[i].bar(range(n_features), list_evectors[i])
        ax[i].set_xticks(range(n_features))
        ax[i].grid('on', linestyle='--')
        ax[i].set_ylim(-0.6, 0.6)
        ax[i].set_xlim(-2, n_features + 1)
        ax[i].axhline(0, color=Colors.gray)
        plt.setp(ax[i].yaxis.get_major_ticks()[0], visible=False)
        plt.setp(ax[i], ylabel='Weight')

        if i == first:
            plt.setp(ax[i], title=title)
        if not i == last:
            plt.setp(ax[i].get_yticklabels()[0], visible=False)
            ax[i].set_xticklabels([])
            ax[i].xaxis.set_ticks_position('none')
        if i == last:
            ax[i].set_xticklabels(x_tick_labels, rotation=-50, ha='left')

    if first == last:
        plt.delaxes(ax[1])

    fig.subplots_adjust(wspace=0, hspace=0.01)
    out_name = re.sub('_+', '_', re.sub(r'[\W]', '_', title))
    plt.savefig(out_name + '.pdf', ftype='pdf', bbox_inches='tight')


plot_evectors(evecs[0:3], x_tick_labels=df_cleaned.columns.values, title='Eigenvectors: 1, 2, 3')
plot_evectors(evecs[3:6], x_tick_labels=df_cleaned.columns.values, title='Eigenvectors: 4, 5, 6')
plot_evectors(evecs[6:9], x_tick_labels=df_cleaned.columns.values, title='Eigenvectors: 7, 8, 9')
```
