## Supplementary File 1 for "Biophysics of microRNA-34a targeting and its influence on down-regulation": characterise_mRNA_all.py

```
#!/usr/bin/env python3
# -*- coding: utf-8 -*-
'''
Usage:
    characterise_mRNA_all.py  (-h | --help)
    characterise_mRNA_all.py  <mcff_directory>


Purpose:
    Examine all the mcff files in mcff_directory and characterise the therein
    contained mRNA primary sequence according to the type of the seed region
    and the residue content in positions 13 to 16.
'''


def main(mcff_directory):
    for f in os.listdir(mcff_directory):
        mcff = rna.parse_mcff_file(os.path.join(mcff_directory, f))
        # mcff_unique = rna.return_unique_mcff(mcff_raw)
        # mcff = rna.return_fixed_len_mcff(mcff_raw, max_number_of_secondaries=1)
        try:
            rna.validate_primary(mcff['primary'])
            mcff_fasta = rna.mcff_to_fasta(mcff, index=0)
            FF = rna.FastaFeatures(mcff_fasta)
            mRNA_primary = FF.mRNA_primary[::-1]
            # top_fasta = M.mcff_fastas[0]
            # _name, _primary, _secondary = rna.unpack_mcff_fasta(top_fasta)
            _name = mcff['name'].split('.')[0]
            _seed_type = rna.return_seed_type(mRNA_primary, reverse=False)
            _residue_slice = rna.return_13_to_16(mRNA_primary)
            print("{:20}\t{}\t{}\t{}".format(_name, _seed_type, _residue_slice, mRNA_primary))

        except ValueError as e:
            print(e)


if __name__ == '__main__':
    from docopt import docopt
    import os
    # import sys
    # sys.path.append('/home/dimitri/code/mRNA_miRNA/feature_extraction')
    import rna_utils as rna

    arguments = docopt(__doc__)
    _path = arguments['<mcff_directory>']
    main(_path)
```
