## Supplementary File 1 for "Biophysics of microRNA-34a targeting and its influence on down-regulation": classify_seed.py

```
#!/usr/bin/env python3
# -*- coding: utf-8 -*-
'''
Usage:
    classify_seed.py  (-h | --help)
    classify_seed.py  <mRNA_sequence>
'''


def return_seed_type(mRNA_primary):
    '''mRNA seed classification
    Assign seed type to mRNA seed sequence.

    Args:
        mRNA_primary <str> mRNA sequence

    Returns:
        mRNA_seed_type <str>
    '''
    reg_8mer = r'ACCGUCAC'
    reg_7mer_m8 = r'[^A]CCGUCAC'
    reg_7mer_a1 = r'ACCGUCA[^C]'
    reg_6mer = r'[^A]CCGUCA[^C]'
    reg_6mer_m8 = r'[^A][^C]CGUCAC'
    reg_6mer_a1 = r'ACCGUC[^A][^C]'
    key_value_list = [(reg_8mer, '8mer'), (reg_7mer_m8, '7mer-m8'),
                      (reg_7mer_a1, '7mer-A1'), (reg_6mer, '6mer'),
                      (reg_6mer_m8, '6mer-m8'), (reg_6mer_a1, '6mer-A1')]
    for key, value in key_value_list:
        if re.match(key, mRNA_primary):
            return value
    return 'Other'


if __name__ == '__main__':
    import re
    from docopt import docopt
    arguments = docopt(__doc__)
    mRNA_primary = arguments['<mRNA_sequence>']
    seed_type = return_seed_type(mRNA_primary)
    print(seed_type)
```
