## Supplementary File 1 for "Biophysics of microRNA-34a targeting and its influence on down-regulation": generate_feature_table.py

```
#!/usr/bin/env python3.6
# -*- coding: utf-8 -*-
'''
Usage:
    generate_feature_table.py   (-h | --help)
    generate_feature_table.py   <mcff_dir>

Arguments:
    <mcff_dir>  A directory containing mcff files (generated by MCFold).
    Typically there will be just one file for each gene, e.g. for the file
    corresponding to the chosen mRNA length. It is assumed that the chosen
    prediction is always the mfe.
'''

import os
import collections
import itertools
import numpy as np

# sys.path.append('/home/dimitri/code/mRNA_miRNA/feature_extraction')
import rna_utils as rna

__author__ = 'Dimitri Schritt'
__copyright__ = 'Copyright (C) Karolinska Institutet - All Rights Reserved 2017'


# PATH = '/home/dimitri/projects/mRNA_miRNA/materials/mcff_latest_from_kibox'
# Path to mcff files

def main():
    lenOD = collections.OrderedDict()
    lenOD_c = padded_values(lenOD, PATH, pre_run=True)
    padded_values(lenOD_c, PATH, pre_run=False)


def padded_values(lenOD, PATH, pre_run=False):
    _is_first = True
    fill_value = np.nan
    mcff_lengths = list()
    # loop through files once to set the max_mcff_length. This parameter is
    # determined by the mcff file that has the least number of unique secondary
    # structures.
    for f in os.listdir(PATH):
        mcff_raw = rna.parse_mcff_file(os.path.join(PATH, f))
        mcff_unique = rna.return_unique_mcff(mcff_raw)
        mcff_lengths.append(len(mcff_unique['secondaries']))
        print('{}: {}'.format(len(mcff_unique['secondaries']), f))
    max_mcff_length = min(mcff_lengths)
    print('Features will be generated based upon the top {} unique secondary structures.'.format(max_mcff_length))
    for f in os.listdir(PATH):
        mcff_raw = rna.parse_mcff_file(os.path.join(PATH, f))
        mcff_unique = rna.return_unique_mcff(mcff_raw)
        mcff = rna.return_fixed_len_mcff(mcff_unique, max_mcff_length)
        try:
            rna.validate_mcff(mcff)
            rna.validate_primary(mcff['primary'])
            M = rna.McffFeatures(mcff)
            top_fasta = M.mcff_fastas[0]
            _name, _primary, _secondary = rna.unpack_mcff_fasta(top_fasta)
            F = rna.FastaFeatures(top_fasta)

            MFeatures = M.Features
            FFeatures = F.Features
            if BLACKLISTED_FEATURES:
                MFeatures = _filter_by_blacklist(MFeatures, BLACKLISTED_FEATURES)
                FFeatures = _filter_by_blacklist(FFeatures, BLACKLISTED_FEATURES)

            width = 25
            _headers = '{:{align}{width}}'.format('complex_name', width=width, align='<')
            _values = '{:{align}{width}}'.format(_name, width=width, align='<')

            # for key, value in M.Features.items():
            for key, value in FFeatures.items():
                if pre_run:
                    if isinstance(value, list):
                        lenOD[key] = max(len(value), lenOD.get(key))
                    else:
                        lenOD[key] = 1
                else:
                    width = max(len(key), len(str(value))) + 2
                    _headers += '{:{align}{width}}'.format(key, width=width, align='>')
                    _values += '{:{align}{width}}'.format(str(value), width=width, align='>')

            for key, value in FFeatures.items():

                # preprocessing: unpacking lists and tuples
                if isinstance(value, list):
                    _value = rna.flatten(value)
                    if len(_value) > 0:
                        if isinstance(_value[0], tuple):
                            _value = list(itertools.chain.from_iterable(_value))
                elif isinstance(value, tuple):
                    _value = list(value)
                else:
                    _value = value
                if not _value:
                    _value = fill_value
                elif isinstance(_value, list) and len(_value) == 1:
                    _value = _value[0]

                if isinstance(_value, (int, float)) or (_value is fill_value):
                    if pre_run:
                        lenOD[key] = max(1, lenOD.get(key))
                    else:
                        if lenOD[key] > 1:
                            _key = '{}_{}'.format(key, 0)
                            width = max(len(_key), len(str(_value))) + 2
                            _headers += '{:{align}{width}}'.format(_key, width=width, align='>')
                            _values += '{:{align}{width}}'.format(_value, width=width, align='>')

                            for ii in range(1, lenOD[key]):
                                _key = '{}_{}'.format(key, ii)
                                width = max(len(_key), len(str(_value))) + 2
                                _headers += '{:{align}{width}}'.format(_key, width=width, align='>')
                                _values += '{:{align}{width}}'.format(fill_value, width=width, align='>')
                        else:
                            width = max(len(key), len(str(_value))) + 2
                            _headers += '{:{align}{width}}'.format(key, width=width, align='>')
                            _values += '{:{align}{width}}'.format(_value, width=width, align='>')

                elif isinstance(_value, list):
                    if pre_run:
                        lenOD[key] = max(len(_value), lenOD.get(key))
                    else:
                        for _i, _v in enumerate(_value):
                            # _key = '{}_{}'.format(key, _i).replace('AUCG_0', 'A').replace('AUCG_1', 'U').replace('AUCG_2', 'C').replace('AUCG_3', 'G')
                            _key = '{}_{}'.format(key, _i)
                            width = max(len(_key), len(str(_v))) + 2
                            _headers += '{:{align}{width}}'.format(_key, width=width, align='>')
                            _values += '{:{align}{width}}'.format(_v, width=width, align='>')

                        max_len = lenOD[key]
                        if max_len > len(_value):
                            for ii in range(len(_value), max_len):
                                _key = '{}_{}'.format(key, ii)
                                width = max(len(_key), len(str(_v))) + 2
                                _headers += '{:{align}{width}}'.format(_key, width=width, align='>')
                                _values += '{:{align}{width}}'.format(fill_value, width=width, align='>')

            if _is_first and not pre_run:
                print(_headers)
                rna.write_line_to_file('features_all.txt', '\t'.join(['complex_name'] + list(M.Features.keys()) + list(F.Features.keys())), flag='w')
                rna.write_line_to_file('features.txt', _headers, flag='w')
                _is_first = False
            if not pre_run:
                print(_values)
            rna.write_line_to_file('features_all.txt', '\t'.join(['{}'.format(i) for i in [_name] + list(M.Features.values()) + list(F.Features.values())]))
            rna.write_line_to_file('features.txt', _values)
            _values = ''
            _headers = ''

        except ValueError as e:
            print(e)
    return lenOD


def _filter_by_blacklist(ordered_dict, blacklist):
    _del_keys = list()
    for _key in ordered_dict.keys():
        if _key in blacklist:
            _del_keys.append(_key)
    for _key in _del_keys:
        del ordered_dict[_key]
    return ordered_dict


def main_to_txt(PATH):
    _is_first = True
    mcff_lengths = list()
    for f in os.listdir(PATH):
        mcff_raw = rna.parse_mcff_file(os.path.join(PATH, f))
        mcff_unique = rna.return_unique_mcff(mcff_raw)
        mcff_lengths.append(len(mcff_unique['secondaries']))
    max_mcff_length = min(mcff_lengths)
    print('Features will be generated based upon the top {} unique secondary structures.'.format(max_mcff_length))
    for f in os.listdir(PATH):
        mcff_raw = rna.parse_mcff_file(os.path.join(PATH, f))
        mcff_unique = rna.return_unique_mcff(mcff_raw)
        mcff = rna.return_fixed_len_mcff(mcff_unique, max_mcff_length)
        try:
            rna.validate_mcff(mcff)
            rna.validate_primary(mcff['primary'])
            M = rna.McffFeatures(mcff)
            top_fasta = M.mcff_fastas[0]
            _name, _primary, _secondary = rna.unpack_mcff_fasta(top_fasta)
            F = rna.FastaFeatures(top_fasta)
            if _is_first:
                if os.path.exists('features.txt'):
                    os.remove('features.txt')
                rna.write_line_to_file('features.txt', '\t'.join([_name] + list(M.Features.keys()) + list(F.Features.keys())))
                _is_first = False
            rna.write_line_to_file('features.txt', '\t'.join(['{}'.format(i) for i in [_name] + list(M.Features.values()) + list(F.Features.values())]))

        except ValueError as e:
            print(e)


if __name__ == '__main__':
    from docopt import docopt
    arguments = docopt(__doc__)
    PATH = arguments['<mcff_dir>']

    # Try to load file of blacklisted features
    BLIST_FILE = '/home/dimitri/code/mRNA_miRNA/feature_extraction/FEATURE_BLACKLIST'
    WHITELISTED_FEATURES = list()
    BLACKLISTED_FEATURES = list()
    if os.path.isfile(BLIST_FILE):
        print('>>> Found blacklist file: {}'.format(BLIST_FILE))
        with open(BLIST_FILE, 'r') as blf:
            for line in blf:
                line = line.strip()
                if line:
                    if line.startswith('#'):
                        BLACKLISTED_FEATURES.append(line.strip('# '))
                    else:
                        WHITELISTED_FEATURES.append(line)

    if BLACKLISTED_FEATURES:
        print('>>> The following features have been blacklisted:')
        for _feature in BLACKLISTED_FEATURES:
            print('>>> ' + _feature)
        print('>>> NOTE: This affects std out and features.txt. The file features_all.txt will contain all features.')

    main()
    # if you want to convert the txt formatted feature table into xlsx format:
    # import pandas as pd
    # df = pd.read_txt('features.txt', sep='\t', index_col=[0])
    # df.to_excel('features.xlsx')
```
