## Supplementary File 1 for "Biophysics of microRNA-34a targeting and its influence on down-regulation": README.md

Usage:
===============================

Command line: generate feature files
-------------------------------
```
./generate_feature_table.py | awk '/En_first/{y=1}y' > features
```

Python
-------------------------------
```
import rna_utils
```


Definitions:
===============================

seed region
-------------------------------
The seed region is identified by counting up to 8 consecutive base pairs
starting nearest to 24 nucleotides from end of the primary sequence and
excluding the UG base pair of the artificial UUCG hairpin. In order to be a
valid seed region, it must form no fewer than 3 base pairs. By definition,
under the here employed numbering convention, the starting indices of the seed
region are (-1, 1). 

position
-------------------------------
Positions of primary and secondary features are defined with respect to the
start of the seed region. Negative indices demarcate positions of mRNA
nucleotides. Positive indices demarcate positions of miRNA nucleotides.

stem
-------------------------------
An unbroken sequence of base pairs. The boundaries of a stem are secondary
features such as loops, bulges, hairpins, and junctions. Mismatches and wobbles
are considered base pairs in the context of a stem.

bulge
-------------------------------
A sequence of unpaired nucleotides occurring exclusively in the mRNA or in the
miRNA demarcated on both sides by a stem.

internal loop
-------------------------------
A sequence of unpaired nucleotides demarcated on both sides by a stem. An
internal loop may be symmetric or asymmetric depending on the relative number
of unpaired nucleotides of the mRNA and the miRNA. If these are equal, the
internal loop is symmetric. 

hairpin loop (hairpin)
-------------------------------
A sequence of unpaired nucleotides demarcated on only one side by a stem.

junction (multiloop)
-------------------------------
The joining of two or more stems.


Features:
===============================

complex_name
-------------------------------
The name of the mRNA-miRNA complex.

En_first
-------------------------------
The lowest binding energy among the energies of predicted secondary structures.

En_first-second
-------------------------------
The absolute value of the difference in binding energies between the lowest and
the second lowest binding energy. The energies here considered are those of
secondary structure predictions that differ by more than a simple
pairing/unpairing of any number of nucleotides.

En_first-last
-------------------------------
The absolute value of the difference in binding energies between the lowest and
the highest binding energy. Uniqueness is as defined as under En_first-second.
When the feature is computed in the context of a collection of mRNA-miRNA
pairs, the number of unique secondary structure that are considered is the same
for all mRNA-miRNA pairs and is determined by the mRNA-miRNA pair with the
fewest unique secondary secondary structure predictions.

mRNA_length
-------------------------------
The number of nucleotides that comprise the mRNA.

mRNA_A_count
-------------------------------
The number of adenines within the mRNA.

mRNA_U_count
-------------------------------
The number of uracils within the mRNA.

mRNA_C_count
-------------------------------
The number of cytosines within the mRNA.

mRNA_G_count
-------------------------------
The number of guanines within the mRNA.

seed_WC_count
-------------------------------
The number of Watson-Crick base pairs within the seed region.

seed_GU_count
-------------------------------
The number of GU wobbles within the seed region.

seed_MM_count
-------------------------------
The number of mismatches within the seed region.

WC_count
-------------------------------
The number of Watson-Crick base pairs.

GU_count
-------------------------------
The number of GU wobbles.

GU_positions
-------------------------------
An exhaustive list of position indices at which instances of GU wobbles are found.

MM_count
-------------------------------
The number of mismatches.

MM_positions
-------------------------------
An exhaustive list of position indices at which instances of mismatches are found.

GGGG_content
-------------------------------
For each and every instance of 4 or more consecutive guanines, return the length of this pattern.

GGGG_count
-------------------------------
The number of instance of 4 or more consecutive guanines.

hairpin_loop_count
-------------------------------
The number of hairpin loops.

hairpin_loop_positions
-------------------------------
For each and every instance of a hairpin loop, the two starting indices of the hairpin loop.

hairpin_loop_lengths
-------------------------------
For each and every instance of a hairpin loop, the number of nucleotides that comprise the hairpin loop.

hairpin_loop_AUCG_counts
-------------------------------
For each and every instance of a hairpin, a set of 4 numbers corresponding to the number As, Us, Cs, and Gs within the hairpin that form base pairs.

internal_loop_count
-------------------------------
The number of internal loops.

internal_loop_positions
-------------------------------
For each and every instance of an internal loop, the starting indices of the internal loop.

internal_loop_lengths
-------------------------------
For each and every instance of a internal loop, the number of nucleotides that comprise the internal loop.

internal_loop_AUCG_counts
-------------------------------
For each and every instance of an internal loop, a set of 4 numbers corresponding to the number As, Us, Cs, and Gs within the internal loop that form base pairs.

bulge_count
-------------------------------
The number of bulges.

bulge_positions
-------------------------------
For each and every instance of a bulge, the starting index of the bulge.

bulge_lengths
-------------------------------
For each and every instance of a bulge, the number nucleotides that comprise the bulge.

bulge_AUCG_counts
-------------------------------
For each and every instance of a bulge, a set of 4 numbers corresponding to the number As, Us, Cs, and Gs within the bulge that form base pairs.

stem_count
-------------------------------
The number of stems.

stem_positions
-------------------------------
For each and every instance of a stem, the two starting indices of the stem.

stem_lengths
-------------------------------
For each and every instance of a stem, the number of base pairs that comprise the stem.

junction_count
-------------------------------
The number of junctions.

junction_positions
-------------------------------
For each and every instance of a junction, the index on the side of the mRNA corresponding to the start of the junction.

junction_unpaired_AUCG_counts
-------------------------------
For each and every instance of a junction, a set of 4 numbers corresponding to the number As, Us, Cs, and Gs within the junction that do not form base pairs.

junction_types
-------------------------------
An integer greater than or equal to 2 corresponding to the number of stems that comprise the junction. 

complementarity_X
-------------------------------
The degree of complementarity for a given nucleotide $`X \in \lbrace A, U, C, G\rbrace`$ is given by the following formula

```math
100 * \frac{the\,number\,of\,X\,within\,the\,mRNA\,that\,form\,WC\,base\,pairs\,with\,the\,miRNA}{the\,total\,number\,of\,X\,within\,the\,mRNA}
```


Dependencies:
===============================
```
re
os
sys
collections
itertools
numpy
forgi
msa_secondary_utils
logging_exceptions
```
