## Supplementary File 1 for "Biophysics of microRNA-34a targeting and its influence on down-regulation": rna_utils.py

```
#!/usr/bin/env python3.6
# -*- coding: utf-8 -*-
import re
import collections
import numpy as np
import forgi.graph.bulge_graph as fgb
from context import msa_secondary_utils as msa


__author__ = 'Dimitri Schritt'
__copyright__ = 'Copyright (C) Karolinska Institutet - All Rights Reserved 2017'

'''
This file contains functions and classes for feature extraction from mcff
formatted data. In the present context, mcff data refers to a dictionary
containing the following four objects: a name <string>, a sequence <string>, a
list of secondary structures (dot bracket notation) <list of strings>, and a
list of energies <list of floats>. The lists are of length >= 2.
'''

MISMATCHES = ['AA', 'CC', 'GG', 'UU', 'AG', 'GA', 'AC', 'CA', 'CU', 'UC']
WOBBLES = ['GU', 'UG']


class Colour(object):
    '''
    Helper class for setting colour in print statements.
    '''
    BLUE = '\033[94m'
    GREEN = '\033[92m'
    WARNING = '\033[93m'
    RED = '\033[91m'
    ENDC = '\033[0m'


def parse_mcff_file(file_path):
    '''
    Read mcff file and return OrderedDict object. This object consists of key
    value pairs.
    The values are as follows:
        name <string>,
        sequence <string>,
        list of secondary structures (dot bracket notation) <list of strings>,
        list of energies <float>,
        list of indices <int>.
    The only checking that is performed at this level is for the existence of
    the above.
    '''
    with open(file_path, 'r') as f:
        _name = None
        _primary = None
        _secondaries = list()
        _energies = list()
        # shorthand notation for secondary structure
        # indicating the number of loops.
        _shorthand = list()
        for line in f:
            line = line.strip()
            if re.match(r'>', line) and not _name:
                _name = line[1:]
            elif re.match(r'[A-Z\d]+$', line):
                _primary = line
            elif re.match(r'\(', line) or re.match(r'\.', line):
                line = line.split()
                if len(line) == 3:
                    _secondaries.append(line[0])
                    _energies.append(float(line[1]))
                    _shorthand.append(line[2])
            # else:
            #     print('OTHER: {}'.format(line))
    if not (_name and _primary and _secondaries and _energies):
        raise ValueError('encountered incomplete mcff file')
    kv_list = [
        ('name', _name), ('primary', _primary), ('secondaries', _secondaries),
        ('energies', _energies), ('indices', list(range(len(_secondaries))))]
    return collections.OrderedDict(kv_list)


def validate_mcff(mcff):
    '''
    Check consistency of mcff format.
    '''
    # each and every secondary structure has the same length as the
    # primary structure.
    if not mcff['primary']:
        print(mcff)
    length_p = len(mcff['primary'])
    for secondary in mcff['secondaries']:
        length_s = len(secondary)
        if not length_p == length_s:
            raise ValueError(('Invalid mcff: Primary and secondary structure '
                             'were found to of unequal length!\n len(primary):'
                              '{}\n len(secondary): {}'.format(length_p, length_s)))

    # there must be one energy value for each and every secondary structure
    len_secs = len(mcff['secondaries'])
    len_ens = len(mcff['energies'])
    if not len_secs == len_ens:
        raise ValueError(('Invalid mcff: The number of secondary structures '
                         'was found to differ from the number of energies! '
                          '\nlen(secondaries): {}\nlen(energies):{}'.format(len_secs, len_ens)))

    # the secondary structures ought to be sorted from energetically
    # most favourable to least favourable.
    if not sorted(mcff['energies']) == mcff['energies']:
        raise ValueError(('Invalid mcff: The secondary structures appear to be '
                         'out of order. It is expected for them to be sorted '
                          'by energetic favourability with the lowest energy '
                          'structure at the top.'))

    # there should be more than one secondary structure in the mcff. Without
    # this the energy features involving differences between different
    # secondary structure predictions become meaningless.
    if len_secs < 2:
        raise ValueError(('Invalid mcff: The mcff must contain at least two '
                         'secondary structure predictions. '
                          '\n{} predictions were found.'.format(len_secs)))


def validate_primary(primary):
    # usually there is one and only one instance of the pattern UUCG.
    # issue warning if there is more than one.
    # raise exception is there is less than one.
    _uucg_count = len(re.findall('UUCG', primary))
    if _uucg_count == 0:
        raise ValueError(
            'Encountered incomplete primary sequence. UUCG not found.')
    if _uucg_count > 1:
        print('WARNING: Encountered pimary sequence with {} instances of UUCG.'.format(_uucg_count))

    # the 'centre' of the UUCG sequence should be located at index -24
    _uucg_index_centers = [np.mean([match.start(), match.end()]) for match in re.finditer('UUCG', primary)]
    _uucg_index_centers_shifted = [len(primary) - i for i in _uucg_index_centers]
    if 24 not in _uucg_index_centers_shifted:
        print(_uucg_index_centers_shifted)
        raise ValueError('UUCG not found to be centred on index -24. Please check primary sequence.')


# TODO: ADD TO TESTS. Add to input validation function
def validate_secondary(secondary, primary):
    '''
    Check that there are no known anomalies in the secondary structure.
    Check that the secondary structure is consistent with the primary
    structure.
    '''
    # The secondary structure of UUCG must be a hairpin: e.g. either (..) or
    # ....
    ids = [(match.start(), match.end()) for match in re.finditer('UUCG', primary)][0]
    corresp_second = secondary[ids[0]: ids[1]]
    if corresp_second not in ['(..)', '....']:
        raise ValueError('Encountered invalid secondary structure: UUCG is not part of a hairpin.')
    if not len(re.findall(r'\(\.\.\.\)', secondary)) == 0:
        raise ValueError('Encountered invalid secondary structure: found instance of (...).')


def return_fixed_len_mcff(mcff, max_number_of_secondaries):
    '''
    Return a new mcff containing at most max_number_of_secondaries.
    '''
    _name = mcff['name']
    _primary = mcff['primary']
    _secondaries = mcff['secondaries'][: max_number_of_secondaries]
    _energies = mcff['energies'][: max_number_of_secondaries]
    _indices = mcff['indices'][: max_number_of_secondaries]
    kv_list = [
        ('name', _name), ('primary', _primary), ('secondaries', _secondaries),
        ('energies', _energies), ('indices', _indices)]
    return collections.OrderedDict(kv_list)


def return_unique_mcff(mcff):
    '''
    Return a new mcff containing only unique secondary structures. A pair of
    secondary structures is considered degenerate if it differs only by an
    opening or closing of a bracket. For instance, (....) and ((..)) are
    considered degenerate.
    '''
    _name = mcff['name']
    _primary = mcff['primary']
    _secondaries = list()
    _energies = list()
    _indices = list()
    for secondary, energy, ind in zip(mcff['secondaries'], mcff['energies'], mcff['indices']):
        centre = len(secondary) - 24
        if not close_brackets(secondary, centre) in _secondaries:
            _secondaries.append(secondary)
            _energies.append(energy)
            _indices.append(ind)

    kv_list = [
        ('name', _name), ('primary', _primary), ('secondaries', _secondaries),
        ('energies', _energies), ('indices', _indices)]
    return collections.OrderedDict(kv_list)


def mcff_to_fasta(mcff, index):
    '''
    Return an mcff fasta by index from mcff.
    '''
    index_max = len(mcff['secondaries']) - 1
    index_min = -(index_max + 1)
    if (index > index_max) or (index < index_min):
        raise IndexError(('List index out of range! You have requested an '
                          'entry with index {}. The accepted range is {} to {}, '
                          'and is given by the number of secondary '
                          'structurs in mcff.'.format(index, index_min, index_max)))
    _name = mcff['name']
    _primary = mcff['primary']
    _secondary = mcff['secondaries'][index]
    _mcff_fasta = construct_mcff_fasta(_name, _primary, _secondary)
    return _mcff_fasta


def close_brackets(secondary, centre):
    '''
    In:  (((((((((.((((((.....((.((((....)))).)))))))))))))))))
    Out: (((((((((.((((((.....((((((((..)))))))))))))))))))))))
    '''
    list_one = list(secondary[: centre - 1])
    list_two = list(secondary[centre + 1:])
    secondary_new = '..'
    one_sec = list_one.pop(-1)
    two_sec = list_two.pop(0)

    while (len(list_one) > 0) and (len(list_two) > 0):
        if one_sec == '(' and two_sec == ')':
            secondary_new = one_sec + secondary_new + two_sec
            one_sec = list_one.pop(-1)
            two_sec = list_two.pop(0)

        elif one_sec == '.' and two_sec == ')':
            secondary_new = one_sec + secondary_new
            one_sec = list_one.pop(-1)

        elif one_sec == '(' and two_sec == '.':
            secondary_new = secondary_new + two_sec
            two_sec = list_two.pop(0)

        elif one_sec == '.' and two_sec == '.':
            secondary_new = '(' + secondary_new + ')'
            one_sec = list_one.pop(-1)
            two_sec = list_two.pop(0)
        else:
            break
    if one_sec == '.' and two_sec == '.':
        secondary_new = '(' + secondary_new + ')'
    else:
        secondary_new = one_sec + secondary_new + two_sec
    secondary_new = ''.join(list_one) + secondary_new + ''.join(list_two)
    assert len(secondary) == len(secondary_new)
    return secondary_new


def return_hairpin_centres(secondary):
    '''
    For each and every occurrence of the pattern '(.+)', determine the index
    corresponding to its 'centre'. Return these in the for of a list.
    Strictly speaking this is actually the index of the first base to the right
    of centre. Given that every hairpin consists of an even number of bases,
    there is no base at the centre of the symmetrical structure.
    E.g. return_hairpin_centres('((..))') == [3]
    '''
    hairpin_ids = list()
    # The number of hairpin residues is always even.
    for match in re.finditer(r'\((\.\.)+\)', secondary):
        start = match.start()
        end = match.end()
        hairpin_ids.append([start, end])
    centres = [int(np.mean(i)) for i in hairpin_ids]
    return centres


def return_stem(primary, secondary, centre):
    '''
    Return stem e.g. the longest uninterrupted sequence of paired nucleotides
    from a given position given by centre.  centre is an integer that
    corresponds to the location of the centre of a pattern such as (..)
    '''
    ps_list = [(i, j) for i, j in zip(primary, secondary)]

    list_one = ps_list[: centre]
    list_two = ps_list[centre:]

    bp_list = list()

    one_pri, one_sec = list_one.pop(-1)
    two_pri, two_sec = list_two.pop(0)

    while (one_sec == two_sec == '.' or (one_sec == '(' and two_sec == ')')):

        # internal symmetrical loop, or hairpin
        if one_sec == two_sec == '.':
            # print('{} {}\t{} {} unpaired'.format(one_pri, two_pri, one_sec, two_sec))
            pass
        # paired
        elif one_sec == '(' and two_sec == ')':
            # print('{} {}\t{} {} paired'.format(one_pri, two_pri, one_sec, two_sec))
            bp_list.append(one_pri + two_pri)
        else:
            break

        if (len(list_one) > 0) and (len(list_two) > 0):
            one_pri, one_sec = list_one.pop(-1)
            two_pri, two_sec = list_two.pop(0)
        else:
            break

    return bp_list


# TODO: add to tests
def return_seed_type(mRNA_primary, reverse=False):
    '''mRNA seed classification
    Assign seed type to mRNA seed sequence.

    Args:
        mRNA_primary <str> mRNA sequence
        reverse <bool> This must be set to True if th emRNA sequence is in the
        form 5' to 3', [default: False]

    Returns:
        mRNA_seed_type <str>
    '''
    sequence = mRNA_primary[:]
    if reverse is True:
        sequence = mRNA_primary[::-1]
    reg_8mer = r'ACCGUCAC'
    reg_7mer_m8 = r'[^A]CCGUCAC'
    reg_7mer_a1 = r'ACCGUCA[^C]'
    reg_6mer = r'[^A]CCGUCA[^C]'
    reg_6mer_m8 = r'[^A][^C]CGUCAC'
    reg_6mer_a1 = r'ACCGUC[^A][^C]'
    key_value_list = [(reg_8mer, '8mer'), (reg_7mer_m8, '7mer-m8'),
                      (reg_7mer_a1, '7mer-A1'), (reg_6mer, '6mer'),
                      (reg_6mer_m8, '6mer-m8'), (reg_6mer_a1, '6mer-A1')]
    for key, value in key_value_list:
        if re.match(key, sequence):
            return value
    return 'Other'


# TODO: add to tests
def return_13_to_16(mRNA_primary):
    '''Return the residues in positions 13 to 16 (inclusive).

    If mRNA primary is shorter than 13 bases, the this function will return X
    for up to 16.

    Args:
        mRNA_primary <str> mRNA sequence

    Returns:
        sub_string <str> the residues from 13 to 16 with default value X.
    '''
    sub_list = ['X'] * (16 + 1 - 13)
    _slice = mRNA_primary[13: 16 + 1]
    for i, j in enumerate(list(_slice)):
        sub_list[i] = j
    sub_string = ''.join(sub_list)
    return sub_string


def return_base_pair_list_seed(primary, secondary, max_len=8):
    '''
    Return a list of base pairs corresponding to the seed sequence. This is
    determined by starting at index -24 and considering the longest stem
    (uninterrupted sequence of paired nucleotides) from there up to length
    max_len.
    '''
    # centre is the index in primary structure space that corresponds to the
    # origin of the seed region. In terms of the sequence, it is the boundary
    # between the mRNA and the miRNA.
    centre = len(secondary) - 24
    hairpin_centres = return_hairpin_centres(secondary)

    # assert centre in hairpin_centres
    if centre not in hairpin_centres:
        raise ValueError(('centre not in hairpin_centres!\nThis could mean '
                          'that centre is not at position 24 from the end of '
                          'the primary sequence or that no hairpin '
                          '(hairpin centre) was found at that location.'))

    longest_stem = return_stem(primary, secondary, centre)

    if len(longest_stem) < 2:
        raise ValueError('''Invalid seed region. The stem was found to have
                         fewer than 2 base pairs.''')
    else:
        # The first 4 bases of the seed region (e.g. (..) or ....) are
        # always excluded because they are an artifact of the experiment
        # and never occur in nature.
        seed = longest_stem[1: max_len + 1]
    return seed


def construct_mcff_fasta(name, primary, secondary):
    '''
    Construct mcff type fasta string. This differs from the usual fasta string
    in that it consists of three lines as opposed to two.
    '''
    return '>{}\n{}\n{}'.format(name, primary, secondary)


def unpack_mcff_fasta(fasta):
    name, primary, secondary = tuple(fasta.split('\n'))
    name = name.strip('>')
    return name, primary, secondary


def mcff_fasta_to_dict(fasta):
    name, primary, secondary = tuple(fasta.split('\n'))
    name = name.strip('>')
    d = {'name': name, 'primary': primary, 'secondary': secondary}
    return d


def return_base_pair_OD(fasta):
    '''
    Return an OrderedDict containing the following:
    - a list of all base pairs (e.g. including mismatches and wobbles).
    - a list of tuples corresponding to the indices of the base pairs.
    NOTE: every mRNA_miRNA pair here analysed has 4 artificially added
    residues 'UUCG'. These are not excluded at this level. It is important to
    keep this in mind for in the subsequent analysis.
    NOTE: there appears to be a change in conventions used in the version of
    forgi released with python2 and that available for python3.
    python2: read base pairs backwards starting with the stem closest to the
    UUCG loop.
    python3: read base pairs backwards starting with the stem furthest from the
    UUCG loop.
    '''
    name, primary, secondary = unpack_mcff_fasta(fasta)
    bg = fgb.BulgeGraph()
    # bg.from_fasta(fasta)
    bg.from_dotbracket(secondary)
    pair_tuples = bg.to_pair_tuples()
    utl = list()
    bp_list = list()
    BasePairs = collections.namedtuple('BasePairs', ['base_pair_list', 'index_list'])
    for i, j in pair_tuples:
        if (i != 0 and j != 0) and ((i, j) not in utl and (j, i) not in utl):
            utl.append((i, j))
            bp_list.append(primary[i - 1] + primary[j - 1])
    BP = BasePairs(bp_list, utl)
    return BP


def _clean_base_pair_OD(base_pair_OD, location_of_possible_contamination):
    '''
    This function checks whether there is a base pair with index tuple
    location_of_possible_contamination and whether this base pair is GU. If
    both conditions are met, the index pair and the base pair are removed
    from the OrderedDict. The intended purpose of this funciton is to
    eleiminate the artificial UG base pair that is found in most mRNA_miRNA
    pairs due to the artificlal UUCG sequence. In some cases the U and the G
    are not base-paired in which case no UG base pair will be found at the
    expected location. In that case the input and output of this function
    should be the same.
    '''
    _bp_list = list()
    _id_list = list()
    BasePairs = collections.namedtuple('BasePairs', ['base_pair_list', 'index_list'])
    hpins = [i for i in base_pair_OD.index_list if (i[1] - i[0]) == 3]
    if location_of_possible_contamination not in base_pair_OD.index_list:
        print(Colour.WARNING + ('WARNING: it may be that the location for the '
                                'artificial UG base pair was incorrect. Please '
                                'check to make sure the secondary structure '
                                'of UUCG is ...., and not (..)') + Colour.ENDC)
        print('Hint: the hairpin indices are: {}'.format(hpins))

    for _id, _bp in zip(base_pair_OD.index_list, base_pair_OD.base_pair_list):
        if _id == location_of_possible_contamination and _bp == 'UG':
            pass
        else:
            _bp_list.append(_bp)
            _id_list.append(_id)
    BP = BasePairs(_bp_list, _id_list)
    return BP


def _shift_base_pair_OD(base_pair_OD, seed_offset):
    _id_list = list()
    BasePairs = collections.namedtuple('BasePairs', ['base_pair_list', 'index_list'])
    for i, j in base_pair_OD.index_list:
        _id_list.append((_apply_shift(i, seed_offset), _apply_shift(j, seed_offset)))
    BP = BasePairs(base_pair_OD.base_pair_list, _id_list)
    return BP


def locate_base_pairs(base_pair_OD, list_of_pairs_to_be_sought, seed_offset=None):
    '''
    base_pair_OD <OrderedDict>
    list_of_pairs_to_be_sought <list>
    seed_offset <int>

    Return a list of tuples giving the positions of each and every instance
    of the pairs found in list_of_pairs_to_be_sought.
    E.g. if list_of_pairs_to_be_sought = ['GU', 'UG'], then this function would
    return the locations of each and every instance of 'GU' or 'UG'.
    If seed_offset is provided, the positions will be remapped to be in the
    frame of the seed region.
    '''

    tuple_list = list()
    for index, bp in zip(base_pair_OD.index_list, base_pair_OD.base_pair_list):
        if bp in list_of_pairs_to_be_sought:
            tuple_list.append((_apply_shift(index[0], seed_offset),
                              _apply_shift(index[1], seed_offset)))
    return tuple_list[::-1]


def _apply_shift(original_index, seed_offset):
    '''
    If seed_offset is an integer, transform index into coordinates centred
    on the seed region.
    It is necessary to have additional logic in this function to determine
    whether to apply the 'mRNA' shift (seed_offset + 1) or the miRNA shift
    (seed_offset + 4) because there are cases in which both elements of an
    index tuple are mRNA indices. A similar case may exist also for miRNAs,
    though this has not been here observed.
    '''
    if seed_offset != 0 and type(seed_offset) == int:
        if original_index <= seed_offset + 1:
            _new_index = original_index - (seed_offset + 1)
        else:
            _new_index = original_index - (seed_offset + 4)
    else:
        _new_index = original_index
    return _new_index


def count_base_pairs(bp_list):
    '''
    Analyse base pair list in terms of the following criteria:
    - the number of Watson-Crick base pairs
    - the number of Wobble base pairs
    - the number of mismatches
    '''
    BasePairDict = collections.OrderedDict()
    wc_list = list()
    gu_list = list()
    mm_list = list()
    for bp in bp_list:
        if bp in MISMATCHES:
            mm_list.append(bp)
        elif bp in WOBBLES:
            gu_list.append(bp)
        else:
            wc_list.append(bp)
    BasePairDict['watson_crick'] = len(wc_list)
    BasePairDict['wobbles'] = len(gu_list)
    BasePairDict['mismatches'] = len(mm_list)
    return BasePairDict


def count(primary, nucleotide):
    '''
    count how many times nucleotide occurres within primary
    '''
    content = collections.Counter(primary)
    return content[nucleotide]


def return_longest_partially_symmetric_pattern(_string, l_pattern, r_pattern, middle=''):
    '''
    Find the match and related indices of a pattern that is symmetric in the
    number of characters it has on each side, yet not necessarily symmetric in
    terms of the characters. Maximise the length of the match.

    Eample with middle:
    _string = 'XYZaaaXccccccNZLKAJ'
    l_pattern = 'a'
    r_pattern = 'c'
    middle = 'X'
    match = [['aaaXccc', (3, 10)]]

    Eample without middle:
    _string = 'XYZaaaccccccNZLKAJ'
    l_pattern = 'a'
    r_pattern = 'c'
    match = [['aaaccc', (3, 9)]]

    Eample overlapping pattern:
    _string = 'XYZcccXccXccNZLKAJ'
    l_pattern = 'c'
    r_pattern = 'c'
    match = [['ccXcc', (4, 9)]]
    '''
    l_pattern = re.escape(l_pattern)
    r_pattern = re.escape(r_pattern)
    middle = re.escape(middle)

    reg = l_pattern + middle + r_pattern
    # find shortest partially symmetric matches
    middle_ids = [i.start() + 1 for i in re.finditer(reg, _string)]
    # find longest matches
    match_list = list()
    if middle_ids:
        for match in re.finditer(r'{}+{}{}+'.format(l_pattern, middle, r_pattern), _string):
            id_start = match.start()
            id_end = match.end()
            id_middle = [i for i in middle_ids if i in range(id_start, id_end)][0]
            sym_width = min([abs(id_start - id_middle), abs(id_end - id_middle - 1)])
            id_start = id_middle - sym_width
            id_end = id_middle + sym_width + len(middle)
            match_string = _string[id_start: id_end]
            match_list.append([match_string, (id_start, id_end)])
    return match_list


class McffFeatures(object):

    '''
    Features of mRNA miRNA pair as derived from an mcff formatted file.
    '''

    def __init__(self, mcff):
        self.mcff = mcff
        self.name = self.mcff['name']
        self.primary = self.mcff['primary']
        self.secondaries = self.mcff['secondaries']
        # TODO: Add to tests
        self.energies = self.mcff['energies']
        self.indices = self.mcff['indices']

    def __repr__(self):
        return 'McffFeatures({}!r)'.format(self.mcff)

    def __str__(self):
        _headers = ''
        _values = ''
        for key, value in self.Features.items():
            width = max(len(key), len(str(value))) + 2
            _headers += '{:{align}{width}}'.format(key, width=width, align='>')
            _values += '{:{align}{width}}'.format(value, width=width, align='>')
        return _headers + '\t' + self.name + '\n' + _values

    # TODO: add to tests
    def energy_difference(self, nth, mth):
        '''
        Return the absolute value of the energy difference between the nth and
        the mth secondary structure prediction.
        '''
        energies = self.energies
        try:
            difference = abs(energies[nth] - energies[mth])
            return difference
        except IndexError:
            print(Colour.RED + 'List index out of range!' + Colour.ENDC)

    @property
    def mcff_fastas(self):
        '''
        Construct a list of mcff fasta strings. The difference between the
        usual fasta string and an mcff fasta string is that the latter consists
        of three lines as opposed to two.
        '''
        fastas_list = list()
        for secondary in self.secondaries:
            _fasta = '>{}\n{}\n{}'.format(self.name, self.primary, secondary)
            fastas_list.append(_fasta)
        return fastas_list

    # Binding energy features
    @property
    def binding_energy_first(self):
        '''
        It is assumed that the binding energies are sorted from lowest to
        highest. Hence, the first entry corresponds to the lowest energy
        structure.
        '''
        return self.energies[0]

    @property
    def binding_energy_first_second(self):
        return self.energy_difference(0, 1)

    @property
    def binding_energy_first_last(self):
        return self.energy_difference(0, -1)

    @property
    def Features(self):
        FDict = collections.OrderedDict()

        FDict['En_first'] = '%.2f' % self.binding_energy_first
        FDict['En_first-second'] = '%.2f' % self.binding_energy_first_second
        FDict['En_first-last'] = '%.2f' % self.binding_energy_first_last

        return FDict


class FastaFeatures(object):

    '''
    Features of mRNA miRNA pair as derived from an mcffFasta formatted file.
    '''

    def __init__(self, mcff_fasta):
        self.mcff_fasta = mcff_fasta
        self.name = mcff_fasta.split('\n')[0].strip('>')
        self.primary = mcff_fasta.split('\n')[1]
        self.secondary = mcff_fasta.split('\n')[2]

    def __repr__(self):
        return 'FastaFeatures({}!r)'.format(self.mcff_fasta)

    def __str__(self):
        _headers = ''
        _values = ''
        for key, value in self.Features.items():
            width = max(len(key), len(str(value))) + 2
            _headers += '{:{align}{width}}'.format(key, width=width, align='>')
            _values += '{:{align}{width}}'.format(value, width=width, align='>')
        return _headers + '\n' + _values

    @property
    def _blacklisted_ids(self):
        return range(self.mRNA_length, self.mRNA_length + 4)

    @property
    def _base_pair_OD(self):
        rawOD = return_base_pair_OD(self.mcff_fasta)
        _x = self.mRNA_length
        return _clean_base_pair_OD(rawOD, (_x + 1, _x + 4))

    @property
    def _base_pair_OD_shifted(self):
        return _shift_base_pair_OD(self._base_pair_OD, self.mRNA_length)

    def _forgi_to_id_dict(self):
        '''
        Return dictionary of secondary motifs and their indices. The keys are
        almost identical to those returned by the forgi module. The first
        letter of the key indicates the type of secondary motif. The second
        letter enumerates the each occurrence of a given motif. The motif
        definitions are as follows:

        h: hairpin_loop  (..) or (....) etc.
        f: fiveprime     The unpaired nucleotides at the 5 prime end.
        t: threeprime    The unpaired nucleotides at the 3 prime end.
        s: stem          Regions of contiguous canonical Watson-Crick base pairs.
        m: multiloop     Single-stranded unpaired regions.
        i: interior_loop Double-stranded unpaired regions flanked by stems
                         on either side.
        b: bulge         Double-stranded unpaired regions flanked by stems
                         on either side. 'Unpairing' is only on the mRNA side
                         or only on the miRNA side.

        NOTE: The above definitions are mostly as found in the forgi module
        and there are a number of discrepancies between these and those used
        elsewhere in mrna_utils. They are as follows.

        stem:
        The forgi definition appears to be more strict than my definition,
        yet it would appear that there are no differences in the output. e.g.
        it seems that forgi considers any unbroken sequence of paired bases to
        be a stem (without excluding wobbles and mismatches as would be
        suggested by their definition).

        internal loop:
        The forgi definition of internal loop is more broad than mine.  A
        feature identified by forgi as an internal loop may be a bulge or an
        internal loop (symmetric or asymmetric). The way to distinguish them is
        by the number of indices returned by forgi.  If forgi returns two
        indices for a given 'internal loop', then it is a bulge. If forgie
        returns 4 indices it is an internal loop.

        multiloop:
        This is remapped to junction.
        Also, forgi appears to recognise only junctions that include an
        unpaired base.

        Reinterpretation of the definitions is performed only in the case of the
        internal loop.
        '''
        bg = fgb.BulgeGraph()
        # bg.from_fasta(self.mcff_fasta)
        bg.from_dotbracket(self.secondary)
        bgs = bg.to_bg_string()
        secondary_motifs_id_dict = dict()
        for i in bgs.split('\n'):
            if i.startswith('define'):
                key = i.split()[1]
                value = [int(index) for index in i.split()[2:]]
                if value:
                    if key[0] == 'i' and len(value) == 2:
                        secondary_motifs_id_dict[key.replace('i', 'b')] = value
                    elif key[0] == 'm':
                        secondary_motifs_id_dict[key.replace('m', 'j')] = value
                    else:
                        secondary_motifs_id_dict[key] = value
        return secondary_motifs_id_dict

    def _secondary_motif_id_dict(self, motif, blacklisted_ids=None):
        '''
        Search through id dict containing all secondary motif indices and
        return those elements pertaining to a specific secondary motif. The
        secondary motif, 'motif', may be provided either by full name or by
        the first letter of the name. e.g. for hairpin loops, 'motif' may be
        given as 'h' or as 'hairpin_loop'.

        blacklisted_ids is a list of ids which may be used to exclude hairpins
        occurring in the corresponding region. For instance,
        blacklisted_ids = [30, 31] will result in any feature whos indices
        overlap include 30 or 31 to be excluded from the id dict.
        '''
        if not blacklisted_ids:
            blacklisted_ids = list()
        d = self._forgi_to_id_dict()
        keys = [i for i in d.keys() if i[0] == motif.lower()[0]]
        id_dict = collections.OrderedDict()
        for key in sorted(keys):
            if key[0] == 'h' and [i for i in d[key] if i in blacklisted_ids]:
                pass
            else:
                id_dict[key] = d[key]
        return id_dict

    def _secondary_motif_count(self, motif):
        '''
        Count how many instances are there of the motif
        '''
        return len(self._secondary_motif_id_dict(motif, self._blacklisted_ids))

    def _secondary_motif_positions(self, motif):
        '''
        Return a list containing the position of the starting residue(s) of any
        instance of the motif. In the case of a stem, or an internall loop, the
        end position is provided also. For example, in the case of a stem
        position is given by a tuple of tuples of the form:
        ((mRNA_starting_position, miRNA_starting_position), (mRNA_ending_position, miRNA_ending_position))
        '''
        this_dict = self._secondary_motif_id_dict(motif, self._blacklisted_ids)

        # offset to transform into coordinates centred on the seed region
        _x = self.mRNA_length
        _m = self.mRNA_length + 1
        positions = list()
        if this_dict.values():
            # a stem or an internal loop
            if len(list(this_dict.values())[0]) == 4:
                for i in this_dict.values():
                    # _shifted_ind_b = (_apply_shift(i[0], _x) + abs(i[0] - i[1]), _apply_shift(i[3], _x))
                    _shifted_ind_b = (_apply_shift(i[1], _x), _apply_shift(i[2], _x))
                    _shifted_ind_e = (_apply_shift(i[0], _x), _apply_shift(i[3], _x))
                    # mRNA pairing with itself
                    if _shifted_ind_e[0] < 0 and _shifted_ind_e[1] < 0:
                        _shifted_ind_e = (i[1] - _m, i[2] - _m)
                    if _shifted_ind_b[0] < 0 and _shifted_ind_e[1] < 0:
                        _shifted_ind_b = (i[0] - _m, i[3] - _m)
                    # seed start
                    if _shifted_ind_b == (0, 0):
                        _shifted_ind_b = (-1, 1)
                    positions.append((_shifted_ind_b, _shifted_ind_e))
            # not a stem and not an internal loop
            else:
                for i in this_dict.values():
                    _shifted_index = _apply_shift(i[0], _x)
                    if i[0] <= _m:
                        _shifted_index += abs(i[0] - i[1])
                    positions.append(_shifted_index)
        return positions[::-1]

    def _secondary_motif_lengths(self, motif):
        '''
        Return a list containing a count the number of bases/base pairs that
        constitute each instance of the motif.
        '''
        this_dict = self._secondary_motif_id_dict(motif, self._blacklisted_ids)
        lengths = list()
        for i in this_dict.values():
            if i[1] in self._blacklisted_ids:
                lengths.append(abs(i[0] - i[1]) + 1 - 1)
            else:
                if len(i) == 4:
                    if abs(i[0] - i[1]) == abs(i[2] - i[3]):
                        lengths.append(abs(i[0] - i[1]) + 1)
                    else:
                        lengths.append((abs(i[0] - i[1]) + 1, abs(i[2] - i[3]) + 1))
                else:
                    lengths.append(abs(i[0] - i[1]) + 1)
        return lengths[::-1]

    def _secondary_motif_AUCG_counts(self, motif):
        '''
        Return a list of tuples for each instance of the motif. Each tuple
        consists of 4 integers given by the number of A, U, C, G that occur in
        that the instance of the motif.
        NOTE: This is not meaningful for stems.
        '''
        if motif.lower()[0] == 's':
            return 'THIS FEATURE HAS NOT BEEN IMPLEMENTED FOR STEMS'
        this_dict = self._secondary_motif_id_dict(motif, self._blacklisted_ids)
        tuple_list = list()
        for i in this_dict.values():
            if len(i) == 4:
                feature_primary_m = self.primary[i[0] - 1: i[1] + 1 - 1]
                feature_primary_mi = self.primary[i[2] - 1: i[3] + 1 - 1]
                tup_m = tuple([count(feature_primary_m, i) for i in list('AUCG')])
                tup_mi = tuple([count(feature_primary_mi, i) for i in list('AUCG')])
                tuple_list.append([tup_m, tup_mi])
            else:
                feature_primary = self.primary[i[0] - 1: i[1] + 1 - 1]
                tup = tuple([count(feature_primary, i) for i in list('AUCG')])
                tuple_list.append(tup)
        return tuple_list[::-1]

    @property
    def Features(self):
        '''
        Return object containing all features contained within this module.
        '''

        FeatureDict = collections.OrderedDict()
        FeatureDict['mRNA_length'] = self.mRNA_length

        # mRNA nucleotide count
        for N in 'AUCG':
            key = 'mRNA_' + N + '_count'
            value = count(self.mRNA_primary, N)
            FeatureDict[key] = value

        # primary
        FeatureDict['GGGG_content'] = self.GGGG_content
        FeatureDict['GGGG_count'] = self.GGGG_count

        # seed region base-pairing
        FeatureDict['seed_WC_count'] = self.seed_watson_crick_count
        FeatureDict['seed_GU_count'] = self.seed_wobbles_count
        FeatureDict['seed_MM_count'] = self.seed_mismatches_count

        # base-pairing
        FeatureDict['WC_count'] = self.watson_crick_count
        FeatureDict['GU_count'] = self.wobbles_count
        FeatureDict['MM_count'] = self.mismatches_count

        FeatureDict['GU_positions'] = self.wobbles_positions
        FeatureDict['MM_positions'] = self.mismatches_positions

        secondary_motifs = ['hairpin_loop', 'internal_loop', 'bulge', 'stem']
        # secondary motif count
        for m in secondary_motifs:
            key = m + '_count'
            value = self._secondary_motif_count(m)
            FeatureDict[key] = value
        # secondary motif positions
        for m in secondary_motifs:
            key = m + '_positions'
            value = self._secondary_motif_positions(m)
            FeatureDict[key] = value
        # secondary motif lengths
        for m in secondary_motifs:
            key = m + '_lengths'
            value = self._secondary_motif_lengths(m)
            FeatureDict[key] = value
        # secondary motif AUCG counts
        for m in secondary_motifs[:-1]:
            key = m + '_AUCG_counts'
            value = self._secondary_motif_AUCG_counts(m)
            FeatureDict[key] = value

        FeatureDict['junction_types'] = self.junction_types
        FeatureDict['junction_count'] = len(self.junction_types)
        FeatureDict['junction_positions'] = self.junction_positions
        FeatureDict['junction_unpaired_AUCG_counts'] = [self._secondary_motif_AUCG_counts('junction')]

        FeatureDict['complementarity_AUCG'] = self.degree_of_complementarity

        return FeatureDict

    @property
    def junction_types(self):
        '''
        Junctions are characterised by the number of stems which they join.
        Clearly the minimum number of stems to form a junction is 2.
        WARNING: Currently this function does not handle junctions of more than 3 stems.
        '''
        j_list = return_longest_partially_symmetric_pattern(self.secondary, ')', '(', middle='')
        j_types = list()
        if j_list:
            stemOD = self._secondary_motif_id_dict('s')
            for j in j_list:
                if len([i for i in stemOD['s0'] if i in range(j[1][0], j[1][1])]) > 0:
                    j_types.append(2)
                else:
                    j_types.append(3)
        return j_types

    @property
    def junction_positions(self):
        '''
        The position of a junction is here defined as the position of the first
        base on the mRNA side that belongs to the junction.
        WARNING: This function has not been tested on junctions that involve an
        extra stem on the miRNA side.
        '''
        j_list = return_longest_partially_symmetric_pattern(self.secondary, ')', '(', middle='')
        if j_list:
            j_id_list = [i[1][0] + abs(i[1][0] - i[1][1]) / 2 for i in j_list]
            j_id_list_shifted = [_apply_shift(i + 1, self.mRNA_length) for i in j_id_list]
        else:
            j_id_list_shifted = []
        return j_id_list_shifted

    # mRNA
    #
    # TODO: tested as part of mRNA_primary. Consider testing separately.
    def _mRNA_end(self):
        _ends = [i.end() - 4 for i in re.finditer('UUCG', self.primary)]
        if len(_ends) > 1:
            _comp_inds = [abs(24 - (len(self.primary) - (i + 2))) for i in _ends]
            _end = _ends[_comp_inds.index(min(_comp_inds))]
        else:
            _end = _ends.pop()
        return _end

    @property
    def mRNA_primary(self):
        _end = self._mRNA_end()
        _mRNA = self.primary[0: _end]
        return _mRNA

    @property
    def mRNA_length(self):
        return len(self.mRNA_primary)

    # TODO: Add to tests
    # miRNA
    #
    @property
    def miRNA_primary(self):
        _start = self._mRNA_end() + 4
        _miRNA = self.primary[_start:]
        return _miRNA

    @property
    def miRNA_length(self):
        return len(self.miRNA_primary)

    # Nucleotide content of mRNA e.g. the number of AUCG
    @property
    def mRNA_A_count(self):
        return count(self.mRNA_primary, 'A')

    @property
    def mRNA_U_count(self):
        return count(self.mRNA_primary, 'U')

    @property
    def mRNA_C_count(self):
        return count(self.mRNA_primary, 'C')

    @property
    def mRNA_G_count(self):
        return count(self.mRNA_primary, 'G')

    # PRIMARY
    #
    @property
    def GGGG_content(self):
        '''
        Return the length of any instance of 4 or more successive Gs.
        '''
        return [len(i) for i in re.findall(r'GGGG+', self.primary)]

    @property
    def GGGG_count(self):
        '''
        Return the number of instance of 4 or more successive Gs.
        '''
        return len(re.findall(r'GGGG+', self.primary))

    # SEED REGION
    #
    # Seed region type: 6mer, 7mer-A1, 7mer-8mer, 8mer [NOT IMPLEMENTED]
    @property
    def seed_base_pairs(self):
        return return_base_pair_list_seed(self.primary, self.secondary)

    @property
    def seed_watson_crick_count(self):
        return count_base_pairs(self.seed_base_pairs)['watson_crick']

    @property
    def seed_wobbles_count(self):
        return count_base_pairs(self.seed_base_pairs)['wobbles']

    @property
    def seed_mismatches_count(self):
        return count_base_pairs(self.seed_base_pairs)['mismatches']

    # SECONDARY STRUCTURE pairing characteristics
    #
    @property
    def base_pair_list(self):
        return self._base_pair_OD.base_pair_list

    @property
    def watson_crick_count(self):
        return count_base_pairs(self.base_pair_list)['watson_crick']

    @property
    def wobbles_count(self):
        return count_base_pairs(self.base_pair_list)['wobbles']

    @property
    def mismatches_count(self):
        return count_base_pairs(self.base_pair_list)['mismatches']

    @property
    def wobbles_positions(self):
        return locate_base_pairs(self._base_pair_OD, WOBBLES, seed_offset=self.mRNA_length)

    @property
    def mismatches_positions(self):
        return locate_base_pairs(self._base_pair_OD, MISMATCHES, seed_offset=self.mRNA_length)

    # COUNT: how many instances are there of the motif
    #
    @property
    def hairpin_loop_count(self):
        return self._secondary_motif_count('hairpin_loop')

    @property
    def internal_loop_count(self):
        return self._secondary_motif_count('internal_loop')

    @property
    def junction_count(self):
        return len(self.junction_types)

    @property
    def bulge_count(self):
        return self._secondary_motif_count('bulge')

    @property
    def stem_count(self):
        return self._secondary_motif_count('stem')

    # POSITION: what is the index of the starting residue of the motif
    #
    @property
    def hairpin_loop_positions(self):
        return self._secondary_motif_positions('hairpin_loop')

    @property
    def internal_loop_positions(self):
        return self._secondary_motif_positions('internal_loop')

    @property
    def bulge_positions(self):
        return self._secondary_motif_positions('bulge')

    @property
    def stem_positions(self):
        return self._secondary_motif_positions('stem')

#    @property
#    def junction_positions(self):
#        return self._secondary_motif_positions('junction')

    # SIZE: how many bases/base pairs constitute the motif
    #
    @property
    def hairpin_loop_lengths(self):
        return self._secondary_motif_lengths('hairpin_loop')

    @property
    def internal_loop_lengths(self):
        return self._secondary_motif_lengths('internal_loop')

#    @property
#    def junction_lengths(self):
#        return self._secondary_motif_lengths('junction')

    @property
    def bulge_lengths(self):
        return self._secondary_motif_lengths('bulge')

    @property
    def stem_lengths(self):
        return self._secondary_motif_lengths('stem')

    # CONTENT: what bases constitute the motif
    # NOTE: The feature is not here implemented for base pairs e.g. for
    # stems. A classification in terms of WC, GU, and MM is implemented for the
    # seed region and could easily be applied to any stem region.
    #
    @property
    def hairpin_loop_AUCG_counts(self):
        return self._secondary_motif_AUCG_counts('hairpin_loop')

    @property
    def internal_loop_AUCG_counts(self):
        return self._secondary_motif_AUCG_counts('internal_loop')

    @property
    def junction_AUCG_unpaired_counts(self):
        return self._secondary_motif_AUCG_counts('junction')

    @property
    def bulge_AUCG_counts(self):
        return self._secondary_motif_AUCG_counts('bulge')

    @property
    def degree_of_complementarity(self):
        '''
        A tuple of 4 elements where each is given by the percentage of the
        count of AUCG in the complementary mRNA divided by the target mRNA.
        '''

        return (self._complementarity('A'), self._complementarity('U'),
                self._complementarity('C'), self._complementarity('G'))

    def _complementarity(self, base):
        '''
        Return 100 times the count of base within the mRNA that is base
        paired (Watson-Crick) with the miRNA divided by the count of base
        within the mRNA.
        '''
        # NOTE: This function is tested as part of degree_of_complementarity.
        all_bp_OD = self._base_pair_OD_shifted
        mRNA_bp_list = list()
        mRNA_id_tup_list = list()
        BasePairs = collections.namedtuple('BasePairs', ['base_pair_list', 'index_list'])
        for bp, id_tup in zip(all_bp_OD.base_pair_list, all_bp_OD.index_list):
            if id_tup[0] < 0 and id_tup[1] > 0:
                mRNA_bp_list.append(bp)
                mRNA_id_tup_list.append(id_tup)
        mRNA_bp_OD = BasePairs(mRNA_bp_list, mRNA_id_tup_list)
        mRNA_wc_list = [bp for bp in mRNA_bp_OD.base_pair_list if bp not in MISMATCHES + WOBBLES]
        mRNA_wc_primary = ''.join([i[0] for i in mRNA_wc_list])
        #  len_target_mRNA = self.mRNA_length
        #  len_complementary_mRNA = len(mRNA_wc_list)
        cnt_target_mRNA_bases = count(self.mRNA_primary, base)
        cnt_complementary_mRNA_bases = count(mRNA_wc_primary, base)
        if cnt_complementary_mRNA_bases == 0:
            return 0.0
        else:
            return round(100.0 * cnt_complementary_mRNA_bases / cnt_target_mRNA_bases)


# TODO: add tests
def return_relative_similarity_pairwiseALIGN(query_secondary, background_secondaries):
    '''
    In:
        query_secondary: <string> secondary structure in dot bracket notation
        background_secondaries: <list> a list of secondaries <string>
    Out:
        the average similarity of query_secondary vs the secondaries of
        background_secondaries.
    '''

    scores = list()
    for _seq in background_secondaries:
        aligned_q, aligned_b = msa.align_pair(query_secondary, _seq)
        scores.append(msa.return_score(aligned_q, aligned_b))
    return np.mean(scores)


# TODO: add tests
def return_relative_similarity(query_secondary, background_secondaries):
    '''
    In:
        query_secondary: <string> secondary structure in dot bracket notation
        background_secondaries: <list> a list of secondaries <string>
    Out:
        the average similarity of query_secondary vs the secondaries of
        background_secondaries.
    '''
    aligned_q = msa.align_one_to_msa(query_secondary)
    aligned_bs = [msa.align_one_to_msa(secondary) for secondary in background_secondaries]
    scores = [msa.return_score(aligned_q, aligned_b) for aligned_b in aligned_bs]

    return np.mean(scores)


# TODO: add tests
def stem_stability(FastaFeatures):
    '''
    Degree of stem stability score:
        normalised to be between 0 and 1.
        1.0 == all stems are bounded by WC base pairs
        0.0 == all stems are bounded by mismatches

    For every stem end that is a WC base pair:
        incriment stability by +1
    For every stem end that is a wobble:
        incriment stability by -1
    For every stem end that is a missmatch:
        incriment stability by -2

    Return stability / stem_count
    # For every stem of length one that consists of a wobble:
    #     incriment stability by -1
    # For every stem of length one that consists of a mismatch:
    #     incriment stability by -2
    '''
    # TODO:
    #   Write tests to fully characterise this function.
    F = FastaFeatures
    BP_OD = F._base_pair_OD_shifted
    _stability = 0
    assert len(F.stem_lengths) == len(F.stem_positions)
    _bounding_base_pairs = list()
    for length, index_tuple_tuple in zip(F.stem_lengths, F.stem_positions):
        for index_tuple in index_tuple_tuple:
            bp = BP_OD.base_pair_list[BP_OD.index_list.index(index_tuple)]
            _bounding_base_pairs.append(bp)
    for bp in _bounding_base_pairs:
        if bp in WOBBLES:
            _stability += -1
        elif bp in MISMATCHES:
            _stability += -2
            # print(rna.Colour.RED + 'FOUND MM: {}'.format(bp) + rna.Colour.ENDC)
    _best_possible = 2 * len(_bounding_base_pairs)
    return 100 * round(float(_best_possible + _stability) / _best_possible, 2)


# TODO: add tests
def stem_short_count(FastaFeatures):
    _lengths = FastaFeatures.stem_lengths
    return len([i for i in _lengths if i <= 2])


# TODO: add tests
def stem_too_short(FastaFeatures):
    '''
    If no stem is found of length less than or equal to 2:
        stability = 0
    For every stem of length less than or equal to 2:
        incriment stability by -1
    '''
    _lengths = FastaFeatures.stem_lengths
    _instability = -1 * len([i for i in _lengths if i <= 2])
    return _instability


# TODO: add tests
class SelfBasepairing(object):

    def __init__(self, FastaFeatures):
        self.FastaFeatures = FastaFeatures

    @property
    def mRNA(self):
        return self.self_base_pairing()[0]

    @property
    def miRNA(self):
        return self.self_base_pairing()[1]

    @property
    def mRNA_and_miRNA(self):
        return self.self_base_pairing()[0] * self.self_base_pairing()[1]

    def self_base_pairing(self):
        '''
        Analyse the secondary structure for self base pairing and return a
        tuple as follows
        No self base pairing:
            return (0, 0)
        Self base pairing at end on both sides and of equal length:
            return (0, 0)
        Self base pairing at both ends and of unequal length:
            return (1, 1)
        Self base pairing only at end on mRNA side:
            return (1, 0)
        Self base pairing only at end on miRNA side:
            return (0, 1)
        Self base pairing not at end:
            return (0, 0)
        '''
        sp = self.FastaFeatures.stem_positions
        # work with only the starting indices of the stem positions
        sp = [i[0] for i in sp]
        sp.remove((-1, 1))
        sp_self = [s for s in sp if s[0] * s[1] > 0]
        sp_nonself = [s for s in sp if s[0] * s[1] < 0]
        si_nonself = [i[0] for i in sp_nonself] + [i[1] for i in sp_nonself]
        # no self base pairing
        if not sp_self:
            return (0, 0)
        else:
            mRNA_self_stem_pos = sorted(sp_self)[0]
            miRNA_self_stem_pos = sorted(sp_self)[-1]
            if mRNA_self_stem_pos[0] > 0:
                mRNA_self_stem_pos = None
            elif sp_nonself and min(si_nonself) < min(mRNA_self_stem_pos):
                mRNA_self_stem_pos = None
            if miRNA_self_stem_pos[0] < 0:
                miRNA_self_stem_pos = None
            elif sp_nonself and max(si_nonself) > max(miRNA_self_stem_pos):
                miRNA_self_stem_pos = None
            # self base pairing at both ends
            if mRNA_self_stem_pos and miRNA_self_stem_pos:
                mRNA_ss_len = abs(mRNA_self_stem_pos[0] - mRNA_self_stem_pos[1])
                miRNA_ss_len = abs(miRNA_self_stem_pos[0] - miRNA_self_stem_pos[1])
                # of unequal length
                if mRNA_ss_len != miRNA_ss_len:
                    return (1, 1)
                # of equal length
                else:
                    return (0, 0)
            # self base pairing only at end of mRNA
            elif mRNA_self_stem_pos:
                return (1, 0)
            # self base pairing only at end of miRNA
            elif miRNA_self_stem_pos:
                return (0, 1)
            # self base pairing not at end
            else:
                return (0, 0)


# TODO: add tests
class Secondary(object):

    def __init__(self, secondary):
        self.secondary = secondary

    @property
    def count_unpaired_mRNA(self):
        match = re.search(r'^[\.]+', self.secondary)
        length = 0
        if match:
            length = len(match.group())
        return(length)

    @property
    def count_unpaired_miRNA(self):
        match = re.search(r'[\.]+$', self.secondary)
        length = 0
        if match:
            length = len(match.group())
        return(length)

    @property
    def short_stem_at_end(self):
        # stem of length one with bulge on mRNA side
        # True cases:
        #   .(..(....((((((((((((((((((((..))))))))))).)))))))))))
        #   (...(....((((((((((((((((((((..)))))))))))).))))))))))
        #   .(((((((((((((((..)))((((((((((..))))))))))))))))))))).)
        #   (((((((((((((....))(((((((((((..))))))))))))))))))))).)
        #   (.((((((((((.((((((((((..)))))))))))))))))))).).
        # False cases (contrived):
        #   ((..(....((((((((((((((((((((..))))))))))).)))))))))))
        #   .((..(....((((((((((((((((((((..))))))))))).)))))))))))
        #   ((..(....((((((((((((((((((((..))))))))))).))))))))).))
        #   ((..(....((((((((((((((((((((..))))))))))).))))))))).)).
        if re.search(r'^[\.]*\(\.', self.secondary):
            return True
        # stem of length one with bulge on miRNA side
        elif re.search(r'\.+\)[\.]*$', self.secondary):
            return True
        else:
            return False


# TODO: add tests
class mRNAFivePrimem(object):

    def __init__(self, FastaFeatures):
        self.FastaFeatures = FastaFeatures

    @property
    def GG(self):
        if self.count_succesive_Gs() >= 2:
            return 1
        else:
            return 0

    @property
    def GGGG(self):
        if self.count_succesive_Gs() >= 4:
            return 1
        else:
            return 0

    def count_succesive_Gs(self):
        m = re.search('^G+', self.FastaFeatures.mRNA_primary)
        if m:
            return len(m.group())
        else:
            return 0


# TODO: add tests
def normalise_list(_list):
    mi = min(_list)
    ma = max(_list)
    _list_norm = [(i - mi) / float(ma - mi) for i in _list]
    return _list_norm


# TODO: add tests
def normalise(_x, _min, _max):
    '''
    output range: [1.0, 0.0]
    max(_x) -> 1.0
    min(_x) -> 0.0
    '''
    if (_max - _min) == 0.0:
        return _x
    else:
        _x_norm = (_x - _min) / float(_max - _min)
        return _x_norm


# FEATURES:
# [x] * nucleotide count for mRNA
# [x] * length of mRNA
# [x] * type of seed region
# [x] * binding energy of top complex
# [x] * binding energy differences:
# [x]   - difference between lowest and second lowest
# [x]   - difference between lowest and highest
# [?]   - anything else?
# [x] * number of Watson-Crick base pairs
# [x] * number of G-U wobble base pairs
# [x] * number of mismatches
# [x] * bulge
# [x]   - position e.g. number of nucleotides after 26 (UUCG + 22nt in the miRNA)
# [x]   - length e.g. (...) would be a bulge of length 3
# [x]   - nucleotide count e.g. number of A of U of C and of G
# [x] * similarity of nucleotide content between the target mRNA and the fully complementary mRNA
# [x] Report all positions counting from the seed region (excluding UUCG).
# [x] - Positive indices for miRNA
# [x] - Negative indices for mRNA
# [x] Make sure AUCG count does not include contribution from UUCG
# [x] Clarify the meaning of multiloop
# [x] Clarify what is a hairpin, a bulge, and an internal loop
# [x] Make sure starting residues of secondary structure features are not included in content analysis
# [ ] Location of the binding site within the UTR
# [ ] First base pair of seed region AU or not
# [ ] 8th (counting form the start of the seed region e.g. after UUCG)
#     complementary or not (can fail also by short seed)
# [ ] Homology between human and mouse.
# [ ] Include more informmation about the energy distribution. e.g. mean, mode,
#     etc.
# [ ] reformat and update docstrings


# NOTE: NOT TESTED
def write_line_to_file(_file, _line, flag='a'):
    with open(_file, flag) as f:
        f.write(_line + '\n')


# NOTE: NOT TESTED
def flatten(S):
    if S == []:
        return S
    if isinstance(S[0], list):
        return flatten(S[0]) + flatten(S[1:])
    return S[:1] + flatten(S[1:])
```
