## Supplementary File 1 for "Biophysics of microRNA-34a targeting and its influence on down-regulation": backup__init__.py

```
# -*- coding: utf-8 -*-
"""Provide easy access to test data."""
import os

_here = os.path.split(__file__)[0]


class _ChainData(object):

    def __init__(self, relative_path):
        self.path = os.path.join(_here, relative_path)
        with open(self.path, 'r') as _f:
            self.string = _f.read()

# generate the below by:
# for i in `ls | grep mcff`; do echo "$i = _ChainData('$i')"; done >> __init__.py
# ACSL1.ft50.L7.R0_incomplete.mcff = _ChainData('ACSL1.ft50.L7.R0_incomplete.mcff')
# ACSL1.ft50.L7.R0.mcff = _ChainData('ACSL1.ft50.L7.R0.mcff')
# ACSL1.ft50.L7.R0.mcff_fasta.txt = _ChainData('ACSL1.ft50.L7.R0.mcff_fasta.txt')
# ACSL1.ft50.L7.R0.mcff.txt = _ChainData('ACSL1.ft50.L7.R0.mcff.txt')


ACSL1_ft50_L7_R0_mcff = _ChainData('ACSL1.ft50.L7.R0.mcff')
ACSL1_ft50_L7_R0_incomplete_mcff = _ChainData('ACSL1.ft50.L7.R0_incomplete.mcff')
ACSL1_ft50_L7_R0_mcff_fasta_txt = _ChainData('ACSL1.ft50.L7.R0.mcff_fasta.txt')
ACSL1_ft50_L7_R0_mcff_txt = _ChainData('ACSL1.ft50.L7.R0.mcff.txt')
```
