## Supplementary File 1 for "Biophysics of microRNA-34a targeting and its influence on down-regulation": helpers.py

```
# -*- coding: utf-8 -*-
import pytest


def raises_with_message(error_class, expected_message, function, *args, **kwargs):
    try:
        function(*args, **kwargs)
        # if we get here, there is a problem because it should have raised
        pytest.fail('Expected to catch {} but did not with the following call:'
                    '\n{}, *({}), **({})'
                    ''.format(error_class, function, args, kwargs))
    except error_class as e:
        if expected_message.lower() in str(e).lower():
            pass  # this is the only passing case
        else:
            pytest.fail('Caught {} as expected but it did not contain the specified'
                        'message.\nExpected: {}\nCaught: {}'
                        ''.format(error_class, expected_message, e.message))


def confirm_does_not_raise(fail_message, function, *args, **kwargs):
    try:
        function(*args, **kwargs)
    except Exception as e:
        final_message = '{}\n  - Original Error:\n{}'.format(fail_message, str(e))
        pytest.fail(final_message, pytrace=True)
```
