## Supplementary File 1 for "Biophysics of microRNA-34a targeting and its influence on down-regulation": test_rna_utils.py

```
# -*- coding: utf-8 -*-
from .context import rna_utils
from . import helpers

import os
import json
import collections
import unittest

'''
TODO:

    Find example of:
        A junction where the bases for the outward facing stem are
        taken from the miRNA.

        A junction consisting of more than three stems.

        A multiple junctions.
'''

_here = os.path.split(__file__)[0]
_data_dir = os.path.join(_here, 'data')

SAMPLE_FILE_MCFF_FILE = os.path.join(_data_dir, 'ACSL1.ft50.L7.R0.mcff')
SAMPLE_FILE_MCFF = os.path.join(_data_dir, 'ACSL1.ft50.L7.R0.mcff.txt')
SAMPLE_FILE_MCFF_INCOMPLETE_FILE = os.path.join(_data_dir, 'ACSL1.ft50.L7.R0_incomplete.mcff')
SAMPLE_FILE_MCFF_FASTAS = os.path.join(_data_dir, 'ACSL1.ft50.L7.R0.mcff_fasta.txt')

SAMPLE_FILE_MCFF_FILE_DEGENERATE = os.path.join(_data_dir, 'ARHGAP1.ft20.L6.R0.mcff')
SAMPLE_FILE_MCFF_FILE_NONDEGENERATE = os.path.join(_data_dir, 'ARHGAP1.ft20.L6.R0_nondegenerate.mcff')

SAMPLE_MCFF_FASTAS = json.load(open('{}'.format(SAMPLE_FILE_MCFF_FASTAS)))

SAMPLE_MCFF = json.load(open('{}'.format(SAMPLE_FILE_MCFF)))

SAMPLE_MCFF_PRIMARY_SECONDARY_EQ_LEN = json.load(open('{}'.format(SAMPLE_FILE_MCFF)))
SAMPLE_MCFF_PRIMARY_SECONDARY_EQ_LEN['primary'] = SAMPLE_MCFF_PRIMARY_SECONDARY_EQ_LEN['primary'][0:10]

SAMPLE_MCFF_ONE_ENERGY_FOR_EACH_SECONDARY = json.load(open('{}'.format(SAMPLE_FILE_MCFF)))
SAMPLE_MCFF_ONE_ENERGY_FOR_EACH_SECONDARY['energies'].pop()

SAMPLE_MCFF_SECONDARIES_OUT_OF_ORDER = json.load(open('{}'.format(SAMPLE_FILE_MCFF)))
SAMPLE_MCFF_SECONDARIES_OUT_OF_ORDER['energies'] = SAMPLE_MCFF_SECONDARIES_OUT_OF_ORDER['energies'][::-1]

SAMPLE_MCFF_FEWER_THAN_TWO_SECONDARIES = json.load(open('{}'.format(SAMPLE_FILE_MCFF)))
SAMPLE_MCFF_FEWER_THAN_TWO_SECONDARIES['secondaries'] = SAMPLE_MCFF_FEWER_THAN_TWO_SECONDARIES['secondaries'][:1]
SAMPLE_MCFF_FEWER_THAN_TWO_SECONDARIES['energies'] = SAMPLE_MCFF_FEWER_THAN_TWO_SECONDARIES['energies'][:1]

# SAMPLE_PRIMARY_NO_UUCG = 'DOGGODAGODDOG'
SAMPLE_PRIMARY_NO_UUCG = 'HOWDOYOURECOGNISETHATSOMEONEISABITITCHYBYJANMASEK'
SAMPLE_PRIMARY_mRNA = 'GGAAUGCUAGAGGAAAGAGAACACUGCCU'
SAMPLE_PRIMARY_mRNA_TWO_UUCG = 'UCAAUGUUCGUCUCUGCCCUGAUGCUGCCU'

SAMPLE_PRIMARY = SAMPLE_MCFF['primary']
SAMPLE_SECONDARY_0 = SAMPLE_MCFF['secondaries'][0]
SAMPLE_STEM_0 = ['UG', 'UU', 'CG', 'CG', 'GC', 'UA', 'CG', 'AU', 'CG', 'AU']

SAMPLE_SEED_0 = ['UU', 'CG', 'CG', 'GC', 'UA', 'CG', 'AU', 'CG']

SAMPLE_SECONDARY_9 = SAMPLE_MCFF['secondaries'][9]

SAMPLE_SECONDARY_INVALID_SEED = '(((((((((((((..)))((((((((((.(..)))))))))))))))))))))..'

SAMPLE_FASTA = '>name\n{}\n{}'.format(SAMPLE_PRIMARY, SAMPLE_SECONDARY_0)
SAMPLE_FASTA_DICT = {'name': 'name', 'primary': SAMPLE_PRIMARY, 'secondary':
                     SAMPLE_SECONDARY_0}

SAMPLE_ORDERED_DICT_BASE_PAIR_COUNT = collections.OrderedDict([
    ('watson_crick', 14), ('wobbles', 3), ('mismatches', 5)])

# This has one hairpin and a two-stem junction.
SAMPLE_FASTA_HAIRPIN_MRNA_EXTREME = '''>CD44.ft50.L2.R0.mcff
GGACAUUAGGCCACUAUGUGUUGUUACUGCCAUUCGUGGCAGUGUCUUAGCUGGUUGU
((((....))))(((((((((((((((((((((..)))))))))))..))))))))))'''

# This has one hairpin and a three-stem junction.
# The junction includes no unpaired bases.
SAMPLE_FASTA_JUNCTION_T1 = '''>MTA2.ft50.L5.R0.mcff
GGUACCUGGUUGGGGGAGGGGGGCGUGCACUGCCAUUCGUGGCAGUGUCUUAGCUGGUUGU
.(((((((((((((((((((..)))))(((((((((..)))))))))))))))))))))))'''

# This has one hairpin and a three-stem junction.
# The junction includes one unpaired base.
SAMPLE_FASTA_JUNCTION_T2 = '''>MAP2K1.ft50.L5.R0.mcff
AAAUCUAGCCAGAGCCCUUCACUGCCAUUCGUGGCAGUGUCUUAGCUGGUUGU
((((((((((((((..))((((((((((..)))))))))).))))))))))))'''

SAMPLE_FASTA_SYMM_INTERNAL_LOOP = '''>name
GGAGACCCCUUGUUGCACUGCCCUUCGUGGCAGUGUCUUAGCUGGUUGU
.((((((((((((((((((..(((..)))..))))))))))))))))))'''

SAMPLE_FASTA_ASYMM_INTERNAL_LOOP_m = '''>name
GUGAGACCCCUUGUUGCACUGCCCUUCGUGGCAGUGUCUUAGCUGGUUGU
.((((((((((((((((((...(((..)))..))))))))))))))))))'''

SAMPLE_FASTA_ASYMM_INTERNAL_LOOP_mi = '''>name
GGAGACCCCUUGUUGCACUGCCCUUCGUGGCAGUGUCUUAGCUGGUUGUU
.((((((((((((((((((..(((..)))...))))))))))))))))))'''

SAMPLE_FASTA_GGGG = '''>MTA2.ft50.L5.R0.mcff
GGUACCUGGUUGGGGGAGGGGGGCGUGCACUGCCAUUCGUGGCAGUGUCUUAGCUGGUUGU
.(((((((((((((((((((..)))))(((((((((..)))))))))))))))))))))))'''

SAMPLE_PRIMARY_UUCG_IN_WRONG_PLACE = '''>name
GGAGACCCCUUGUUGCACUGCUUCGCCUGGCAGUGUCUUAGCUGGUUGU
.((((((((((((((((((..(((..)))..))))))))))))))))))'''

SAMPLE_FASTA_NO_HAIRPIN_AT_24_AND_ODD_NUMBER_OF_UNPAIRED_BASES = '''>name
UGGGGGCUCUCAUGUUUUCAACUGCCAUUCGUGGCAGUGUCUUAGCUGGUUGU
((((((((....))))))))(((((((((((((((...)))))))))))))))'''

SAMPLE_FASTA_SELF_PAIRING_MI_12 = '''>CCND3.ft50.L7.R0.mcff
CCUGGGGGCUCUCAUGUUUUCAACUGCCAUUCGUGGCAGUGUCUUAGCUGGUUGU
(((((((..)))))))((((((((((((((..))))))))(.(...)))))))).'''

SAMPLE_FASTA_SELF_PAIRING_MI_15 = '''>CCND3.ft50.L7.R0.mcff
CCUGGGGGCUCUCAUGUUUUCAACUGCCAUUCGUGGCAGUGUCUUAGCUGGUUGU
(((((((..)))))))((((((((((((((..))))))))((....)))))))).'''

SAMPLE_FASTA_TWO_UUCG = '''>ARAF.ft50.L8.R0.mcff
UCAAUGUUCGUCUCUGCCCUGAUGCUGCCUUUCGUGGCAGUGUCUUAGCUGGUUGU
(((((((..)))(((((((((((((((((((..)))))))))))))))))))))))'''

SAMPLE_BASE_PAIR_LIST_WITH_ARTIFACT = ['AU', 'AG', 'UU', 'GU', 'CG', 'UG',
                                       'AU', 'GC', 'AG', 'GA', 'GU', 'AU',
                                       'GC', 'AU', 'CG', 'AU', 'CG', 'UA',
                                       'GC', 'CG', 'CG', 'UU', 'UG']

SAMPLE_BASE_PAIR_LIST = ['AU', 'AG', 'UU', 'GU', 'CG', 'UG', 'AU', 'GC', 'AG',
                         'GA', 'GU', 'AU', 'GC', 'AU', 'CG', 'AU', 'CG', 'UA',
                         'GC', 'CG', 'CG', 'UU']

SAMPLE_INDEX_LIST = [(3, 55), (4, 54), (5, 53), (6, 52), (7, 51), (8, 50),
                     (9, 49), (10, 48), (11, 47), (12, 46), (13, 45), (14, 44),
                     (19, 43), (21, 42), (22, 41), (23, 40), (24, 39),
                     (25, 38), (26, 37), (27, 36), (28, 35), (29, 34)]

SAMPLE_INDEX_LIST_SHIFTED = [(-27, 22), (-26, 21), (-25, 20), (-24, 19),
                             (-23, 18), (-22, 17), (-21, 16), (-20, 15),
                             (-19, 14), (-18, 13), (-17, 12), (-16, 11),
                             (-11, 10), (-9, 9), (-8, 8), (-7, 7), (-6, 6),
                             (-5, 5), (-4, 4), (-3, 3), (-2, 2), (-1, 1)]

SAMPLE_BASE_PAIR_LIST_JUNCTION_T1 = ['GU', 'UG', 'AU', 'CU', 'CG', 'UG', 'GU',
                                     'GC', 'UG', 'UA', 'GU', 'GU', 'GC', 'GU',
                                     'GG', 'AU', 'GG', 'GC', 'GG', 'CG', 'AU',
                                     'CG', 'UA', 'GC', 'CG', 'CG', 'AU']

SAMPLE_INDEX_LIST_JUNCTION_T1 = [(2, 61), (3, 60), (4, 59), (5, 58), (6, 57),
                                 (7, 56), (8, 55), (9, 54), (10, 53), (11, 52),
                                 (12, 51), (13, 50), (14, 49), (15, 48),
                                 (16, 27), (17, 26), (18, 25), (19, 24),
                                 (20, 23), (28, 47), (29, 46), (30, 45),
                                 (31, 44), (32, 43), (33, 42), (34, 41),
                                 (35, 40)]

SAMPLE_INDEX_LIST_JUNCTION_T1_SHIFTED = [(-34, 22), (-33, 21), (-32, 20),
                                         (-31, 19), (-30, 18), (-29, 17),
                                         (-28, 16), (-27, 15), (-26, 14),
                                         (-25, 13), (-24, 12), (-23, 11),
                                         (-22, 10), (-21, 9), (-20, -9),
                                         (-19, -10), (-18, -11), (-17, -12),
                                         (-16, -13), (-8, 8), (-7, 7), (-6, 6),
                                         (-5, 5), (-4, 4), (-3, 3), (-2, 2),
                                         (-1, 1)]


class FunctionTests(unittest.TestCase):

    '''
    Tests for public and private functions contained within the module.
    '''

    def setUp(self):

        self.base_pair_OD_raw = rna_utils.return_base_pair_OD(SAMPLE_FASTA)
        self.base_pair_OD = rna_utils._clean_base_pair_OD(self.base_pair_OD_raw, (30, 33))
        self.base_pair_OD_shifted = rna_utils._shift_base_pair_OD(self.base_pair_OD, 29)

        self.base_pair_OD_hp_raw = rna_utils.return_base_pair_OD(SAMPLE_FASTA_HAIRPIN_MRNA_EXTREME)
        self.base_pair_OD_hp = rna_utils._clean_base_pair_OD(self.base_pair_OD_hp_raw, (33, 36))

        self.base_pair_OD_jt1_raw = rna_utils.return_base_pair_OD(SAMPLE_FASTA_JUNCTION_T1)
        self.base_pair_OD_jt1 = rna_utils._clean_base_pair_OD(self.base_pair_OD_jt1_raw, (36, 39))
        self.base_pair_OD_jt1_shifted = rna_utils._shift_base_pair_OD(self.base_pair_OD_jt1, 35)

#        self.base_pair_OD_sl_raw = rna_utils.return_base_pair_OD(SAMPLE_FASTA_SYMM_INTERNAL_LOOP)
#        self.base_pair_OD_sl = rna_utils._clean_base_pair_OD(self.base_pair_OD_sl_raw, (24, 27))
#
#        self.base_pair_OD_sl_m_raw = rna_utils.return_base_pair_OD(SAMPLE_FASTA_ASYMM_INTERNAL_LOOP_m)
#        self.base_pair_OD_sl_m = rna_utils._clean_base_pair_OD(self.base_pair_OD_sl_m_raw, (25, 28))
#
#        self.base_pair_OD_sl_mi_raw = rna_utils.return_base_pair_OD(SAMPLE_FASTA_ASYMM_INTERNAL_LOOP_mi)
#        self.base_pair_OD_sl_mi = rna_utils._clean_base_pair_OD(self.base_pair_OD_sl_mi_raw, (24, 27))

    def test_parse_mcff_file(self):

        # format and content of object returned from file is as expected
        for key in SAMPLE_MCFF.keys():
            self.assertEqual(rna_utils.parse_mcff_file(SAMPLE_FILE_MCFF_FILE)[key],
                             SAMPLE_MCFF[key])

        # raises ValueError with user friendly message for incomplete file
        helpers.raises_with_message(ValueError, 'incomplete mcff file',
                                    rna_utils.parse_mcff_file,
                                    SAMPLE_FILE_MCFF_INCOMPLETE_FILE)

    def test_validate_mcff(self):

        # raises ValueError with user friendly message if the primary
        # and secondary structures are not of identical length
        helpers.raises_with_message(ValueError,
                                    'Invalid mcff: Primary and secondary structure',
                                    rna_utils.validate_mcff,
                                    SAMPLE_MCFF_PRIMARY_SECONDARY_EQ_LEN)

        # raises ValueError with user friendly message if the number of
        # energies is not equal to the number of secondary structure
        helpers.raises_with_message(ValueError,
                                    'Invalid mcff: The number of secondary structure',
                                    rna_utils.validate_mcff,
                                    SAMPLE_MCFF_ONE_ENERGY_FOR_EACH_SECONDARY)

        # raises ValueError with user friendly message if the secondary
        # structures/energies are not ordered from most energetically
        # most favourable to least favourable
        helpers.raises_with_message(ValueError, (
                                    'Invalid mcff: The secondary structures '
                                    'appear to be out of order'),
                                    rna_utils.validate_mcff,
                                    SAMPLE_MCFF_SECONDARIES_OUT_OF_ORDER)

        # raises ValueError with user friendly message if mcff contains
        # fewer than two secondary structures
        helpers.raises_with_message(ValueError, (
                                    'Invalid mcff: The mcff must contain at '
                                    'least two secondary structure predictions'),
                                    rna_utils.validate_mcff,
                                    SAMPLE_MCFF_FEWER_THAN_TWO_SECONDARIES)

    def test_validate_primary(self):
        # raises ValueError with user friendly message if the primary
        # sequence does not contain UUCG
        helpers.raises_with_message(ValueError, ('Encountered incomplete '
                                    'primary sequence. UUCG not found.'),
                                    rna_utils.validate_primary,
                                    SAMPLE_PRIMARY_NO_UUCG)

        # raises ValueError if UUCG in primary sequence is not found to be
        # centred at index -24
        helpers.raises_with_message(ValueError, ('UUCG not found to be centred '
                                                 'on index -24. Please check '
                                                 'primary sequence.'),
                                    rna_utils.validate_primary,
                                    SAMPLE_PRIMARY_UUCG_IN_WRONG_PLACE)

    def test_return_fixed_len_mcff(self):
        for _len in range(60):
            mcff_fixed_len = rna_utils.return_fixed_len_mcff(SAMPLE_MCFF, _len)
            self.assertEqual(SAMPLE_MCFF['name'], mcff_fixed_len['name'])
            self.assertEqual(SAMPLE_MCFF['primary'], mcff_fixed_len['primary'])
            self.assertEqual(SAMPLE_MCFF['secondaries'][: _len], mcff_fixed_len['secondaries'])
            self.assertEqual(SAMPLE_MCFF['energies'][: _len], mcff_fixed_len['energies'])
            self.assertEqual(SAMPLE_MCFF['indices'][: _len], mcff_fixed_len['indices'])

    def test_return_unique_mcff(self):
        mcff_degenerate = rna_utils.parse_mcff_file(SAMPLE_FILE_MCFF_FILE_DEGENERATE)
        mcff_nondegenerate = rna_utils.parse_mcff_file(SAMPLE_FILE_MCFF_FILE_NONDEGENERATE)
        self.assertEqual(rna_utils.return_unique_mcff(mcff_degenerate)['name'], mcff_nondegenerate['name'])
        self.assertEqual(rna_utils.return_unique_mcff(mcff_degenerate)['primary'], mcff_nondegenerate['primary'])
        self.assertEqual(rna_utils.return_unique_mcff(mcff_degenerate)['secondaries'], mcff_nondegenerate['secondaries'])
        self.assertEqual(rna_utils.return_unique_mcff(mcff_degenerate)['energies'], mcff_nondegenerate['energies'])
        self.assertEqual(rna_utils.return_unique_mcff(mcff_degenerate)['indices'], [0, 1, 2, 3, 4, 5, 6, 9, 11, 12, 15, 17])

    def test_mcff_to_fasta(self):
        _fasta0 = '''>ACSL1.ft50.L7.R0.mcff
        GGAAUGCUAGAGGAAAGAGAACACUGCCUUUCGUGGCAGUGUCUUAGCUGGUUGU
        ..((((((((((((....(.((((((((((..)))))))))))))))))))))))'''.replace(' ', '')
        _fasta4 = '''>ACSL1.ft50.L7.R0.mcff
        GGAAUGCUAGAGGAAAGAGAACACUGCCUUUCGUGGCAGUGUCUUAGCUGGUUGU
        ..(((((((((.....((((((((((((((..)))))))))))))))))))))))'''.replace(' ', '')
        _fasta49 = '''>ACSL1.ft50.L7.R0.mcff
        GGAAUGCUAGAGGAAAGAGAACACUGCCUUUCGUGGCAGUGUCUUAGCUGGUUGU
        .(((.((((((((..(((...(((((((((..)))))))))))))))))))))))'''.replace(' ', '')
        self.assertEqual(rna_utils.mcff_to_fasta(SAMPLE_MCFF, 4), _fasta4)
        self.assertEqual(rna_utils.mcff_to_fasta(SAMPLE_MCFF, 49), _fasta49)
        self.assertEqual(rna_utils.mcff_to_fasta(SAMPLE_MCFF, -1), _fasta49)
        self.assertEqual(rna_utils.mcff_to_fasta(SAMPLE_MCFF, -50), _fasta0)
        helpers.raises_with_message(IndexError, ('List index out of range! You '
                                                 'have requested an entry with '
                                                 'index 50. The accepted range is '
                                                 '-50 to 49, and is given by the number '
                                                 'of secondary structurs in mcff.'),
                                    rna_utils.mcff_to_fasta,
                                    SAMPLE_MCFF, 50)
        helpers.raises_with_message(IndexError, ('List index out of range! You '
                                                 'have requested an entry with '
                                                 'index -51. The accepted range is '
                                                 '-50 to 49, and is given by the number '
                                                 'of secondary structurs in mcff.'),
                                    rna_utils.mcff_to_fasta,
                                    SAMPLE_MCFF, -51)

    def test_close_brackets(self):
        in_sec = '(((((((((.((((((.....(((((((....))))))))))))))))))))))'
        out_sec = '(((((((((.((((((.....((((((((..)))))))))))))))))))))))'
        self.assertEqual(rna_utils.close_brackets(in_sec, centre=30), out_sec)
        in_sec = '.(((((((((((((((.....((((((((..)))))))))))))))))))))))'
        out_sec = '.(((((((((((((((.....((((((((..)))))))))))))))))))))))'
        self.assertEqual(rna_utils.close_brackets(in_sec, centre=30), out_sec)
        in_sec = '..(((((((((((((.....(((((((((..))))))))))))))))))).)))'
        out_sec = '..(((((((((((((.....(((((((((..))))))))))))))))))).)))'
        self.assertEqual(rna_utils.close_brackets(in_sec, centre=30), out_sec)
        in_sec = '.((((((((.((((((.....((((((((..)))))))))))))))))))))).'
        out_sec = '(((((((((.((((((.....((((((((..)))))))))))))))))))))))'
        self.assertEqual(rna_utils.close_brackets(in_sec, centre=30), out_sec)

    def test_return_hairpin_centres(self):
        self.assertEqual(rna_utils.return_hairpin_centres(SAMPLE_SECONDARY_0), [31])
        self.assertEqual(rna_utils.return_hairpin_centres(SAMPLE_SECONDARY_9), [14, 31])

    def test_return_stem(self):
        _primary = 'UUUAUAAGAAAAGUGACUGCCAUUCGUGGCAGUGUCUUAGCUGGUUGU'
        _secondary = '(((((((((((((((((((((((..)))))))))))))))))))))))'
        _stem = ['UG', 'AU', 'CG', 'CG', 'GC', 'UA', 'CG', 'AU', 'GG', 'UU',
                 'GC', 'AU', 'AU', 'AA', 'AG', 'GC', 'AU', 'AG', 'UG', 'AU',
                 'UU', 'UG', 'UU']
        self.assertEqual(rna_utils.return_stem(SAMPLE_PRIMARY, SAMPLE_SECONDARY_0, 31), SAMPLE_STEM_0)
        self.assertEqual(rna_utils.return_stem(_primary, _secondary, 24), _stem)

    def test_return_base_pair_list_seed(self):
        # basic test
        self.assertEqual(rna_utils.return_base_pair_list_seed(SAMPLE_PRIMARY, SAMPLE_SECONDARY_0, 8), SAMPLE_SEED_0)

        # test the max_len argument
        for max_len in reversed(range(11)):
            self.assertLessEqual(len(rna_utils.return_base_pair_list_seed(SAMPLE_PRIMARY, SAMPLE_SECONDARY_0, max_len)), max_len)

        # raises ValueError with user friendly message if seed region is found to
        # have fewer than one base pairs
        helpers.raises_with_message(ValueError, 'Invalid seed region',
                                    rna_utils.return_base_pair_list_seed,
                                    SAMPLE_PRIMARY,
                                    SAMPLE_SECONDARY_INVALID_SEED)

    def test_construct_mcff_fasta(self):
        self.assertEqual(rna_utils.construct_mcff_fasta('name', SAMPLE_PRIMARY,
                                                        SAMPLE_SECONDARY_0),
                         SAMPLE_FASTA)

    def test_unpack_mcff_fasta(self):
        self.assertEqual(rna_utils.unpack_mcff_fasta(SAMPLE_FASTA),
                         ('name', SAMPLE_PRIMARY, SAMPLE_SECONDARY_0))

    def test_mcff_fasta_to_dict(self):
        self.assertEqual(rna_utils.mcff_fasta_to_dict(SAMPLE_FASTA),
                         SAMPLE_FASTA_DICT)

    def test_return_base_pair_OD(self):
        self.assertEqual(self.base_pair_OD_raw.base_pair_list,
                         SAMPLE_BASE_PAIR_LIST_WITH_ARTIFACT)

        base_pair_list_from_ids = [SAMPLE_PRIMARY[i[0] - 1] +
                                   SAMPLE_PRIMARY[i[1] - 1]
                                   for i in self.base_pair_OD_raw.index_list]

        self.assertEqual(base_pair_list_from_ids,
                         SAMPLE_BASE_PAIR_LIST_WITH_ARTIFACT)

    def test__clean_base_pair_OD(self):
        self.assertEqual(self.base_pair_OD.base_pair_list, SAMPLE_BASE_PAIR_LIST)
        self.assertEqual(self.base_pair_OD.index_list, SAMPLE_INDEX_LIST)
        self.assertEqual(self.base_pair_OD_jt1.base_pair_list, SAMPLE_BASE_PAIR_LIST_JUNCTION_T1)
        self.assertEqual(self.base_pair_OD_jt1.index_list, SAMPLE_INDEX_LIST_JUNCTION_T1)

    def test__shift_base_pair_OD(self):
        self.assertEqual(self.base_pair_OD_shifted.base_pair_list, SAMPLE_BASE_PAIR_LIST)
        self.assertEqual(self.base_pair_OD_shifted.index_list, SAMPLE_INDEX_LIST_SHIFTED)
        self.assertEqual(self.base_pair_OD_jt1_shifted.base_pair_list, SAMPLE_BASE_PAIR_LIST_JUNCTION_T1)
        self.assertEqual(self.base_pair_OD_jt1_shifted.index_list, SAMPLE_INDEX_LIST_JUNCTION_T1_SHIFTED)

    def test_locate_base_pairs(self):

        # basic test: in original reference frame
        self.assertEqual(rna_utils.locate_base_pairs(self.base_pair_OD_raw,
                                                     rna_utils.WOBBLES),
                         [(30, 33), (13, 45), (8, 50), (6, 52)])
#                         [(13, 45), (8, 50), (6, 52), (30, 33)])

        # basic test with seed_offset e.g. in reference frame of seed region
        self.assertEqual(rna_utils.locate_base_pairs(self.base_pair_OD_raw,
                                                     rna_utils.WOBBLES,
                                                     seed_offset=len(SAMPLE_PRIMARY_mRNA)),
                         [(0, 0), (-17, 12), (-22, 17), (-24, 19)])
#                         [(-17, 12), (-22, 17), (-24, 19), (0, 0)])

        # test with seed_offset in case of hairpin loop on extreme of mRNA
        self.assertEqual(rna_utils.locate_base_pairs(self.base_pair_OD_hp_raw,
                                                     rna_utils.MISMATCHES,
                                                     seed_offset=32),
                         [(-9, 9), (-14, 16), (-15, 17), (-18, 20), (-30, -23)])
#                         [(-30, -23), (-14, 16), (-15, 17), (-18, 20), (-9, 9)])

    def test__apply_shift(self):
        self.assertEqual(rna_utils._apply_shift(original_index=5, seed_offset=None), 5)
        self.assertEqual(rna_utils._apply_shift(original_index=5, seed_offset=0), 5)
        self.assertEqual(rna_utils._apply_shift(original_index=5, seed_offset=4), 0)
        self.assertEqual(rna_utils._apply_shift(original_index=8, seed_offset=4), 0)
        self.assertEqual(rna_utils._apply_shift(original_index=4, seed_offset=4), -1)
        self.assertEqual(rna_utils._apply_shift(original_index=9, seed_offset=4), 1)

    def test_count_base_pairs(self):
        self.assertEqual(rna_utils.count_base_pairs(SAMPLE_BASE_PAIR_LIST),
                         SAMPLE_ORDERED_DICT_BASE_PAIR_COUNT)

    def test_return_longest_partially_symmetric_pattern(self):
        self.assertEqual(rna_utils.return_longest_partially_symmetric_pattern('XYZaaaXccccccNZLKAJ', 'a', 'c', 'X'),
                         [['aaaXccc', (3, 10)]])
        self.assertEqual(rna_utils.return_longest_partially_symmetric_pattern('XYZaaaccccccNZLKAJ', 'a', 'c'),
                         [['aaaccc', (3, 9)]])
        self.assertEqual(rna_utils.return_longest_partially_symmetric_pattern('XYZaaaccccccNZLKAJ', 'a', 'b'),
                         [])
        self.assertEqual(rna_utils.return_longest_partially_symmetric_pattern('XYZaaaXcccccaXccNZLKAJ', 'a', 'c', 'X'),
                         [['aaaXccc', (3, 10)], ['aXc', (12, 15)]])
        self.assertEqual(rna_utils.return_longest_partially_symmetric_pattern('XYZcccXccXccNZLKAJ', 'c', 'c', 'X'),
                         [['ccXcc', (4, 9)]])
        self.assertEqual(rna_utils.return_longest_partially_symmetric_pattern(SAMPLE_SECONDARY_INVALID_SEED, ')', '('),
                         [[')))(((', (15, 21)]])


class McffFeaturesTests(unittest.TestCase):

    '''
    Tests for McffFeatures class.
    '''

    def setUp(self):
        self.features = rna_utils.McffFeatures(SAMPLE_MCFF)

    def test_mcff_fasta(self):
        self.assertEqual(self.features.mcff_fastas, SAMPLE_MCFF_FASTAS)

    def test_binding_energy_first(self):
        self.assertEqual(self.features.binding_energy_first, -48.371)

    def test_binding_energy_first_second(self):
        self.assertEqual(self.features.binding_energy_first_second, abs(-48.371 + 48.1))

    def test_features_binding_energy_first_last(self):
        self.assertEqual(self.features.binding_energy_first_last, abs(-48.371 + 46.745))


class FastaTests(unittest.TestCase):

    '''
    Tests for FastaFeatures class.
    '''

    def setUp(self):
        self.features = rna_utils.FastaFeatures(SAMPLE_FASTA)
        self.features_hpin = rna_utils.FastaFeatures(SAMPLE_FASTA_HAIRPIN_MRNA_EXTREME)
        self.features_junc_t1 = rna_utils.FastaFeatures(SAMPLE_FASTA_JUNCTION_T1)
        self.features_junc_t2 = rna_utils.FastaFeatures(SAMPLE_FASTA_JUNCTION_T2)
        self.features_sil = rna_utils.FastaFeatures(SAMPLE_FASTA_SYMM_INTERNAL_LOOP)
        self.features_asilm = rna_utils.FastaFeatures(SAMPLE_FASTA_ASYMM_INTERNAL_LOOP_m)
        self.features_asilmi = rna_utils.FastaFeatures(SAMPLE_FASTA_ASYMM_INTERNAL_LOOP_mi)
        self.features_gggg = rna_utils.FastaFeatures(SAMPLE_FASTA_GGGG)
        self.features_bpairmi12 = rna_utils.FastaFeatures(SAMPLE_FASTA_SELF_PAIRING_MI_12)
        self.features_bpairmi15 = rna_utils.FastaFeatures(SAMPLE_FASTA_SELF_PAIRING_MI_15)
        self.features_two_uucg = rna_utils.FastaFeatures(SAMPLE_FASTA_TWO_UUCG)

    def test_junction_types(self):
        self.assertEqual(self.features.junction_types, [])
        self.assertEqual(self.features_hpin.junction_types, [2])
        self.assertEqual(self.features_junc_t1.junction_types, [3])
        self.assertEqual(self.features_junc_t2.junction_types, [3])
        self.assertEqual(self.features_sil.junction_types, [])
        self.assertEqual(self.features_asilm.junction_types, [])
        self.assertEqual(self.features_asilmi.junction_types, [])
        self.assertEqual(self.features_bpairmi12.junction_types, [2, 3])
        self.assertEqual(self.features_bpairmi15.junction_types, [2, 3])

    def test_mRNA_primary(self):
        self.assertEqual(self.features.mRNA_primary, SAMPLE_PRIMARY_mRNA)

    def test_mRNA_length(self):
        self.assertEqual(self.features.mRNA_length, len(SAMPLE_PRIMARY_mRNA))
        self.assertEqual(self.features_two_uucg.mRNA_length, len(SAMPLE_PRIMARY_mRNA_TWO_UUCG))

    def test_mRNA_A_count(self):
        self.assertEqual(self.features.mRNA_A_count, 11)

    def test_mRNA_U_count(self):
        self.assertEqual(self.features.mRNA_U_count, 4)

    def test_mRNA_C_count(self):
        self.assertEqual(self.features.mRNA_C_count, 5)

    def test_mRNA_G_count(self):
        self.assertEqual(self.features.mRNA_G_count, 9)

    def test_GGGG_content(self):
        self.assertEqual(self.features.GGGG_content, [])
        self.assertEqual(self.features_gggg.GGGG_content, [5, 6])

    def test_GGGG_count(self):
        self.assertEqual(self.features.GGGG_count, 0)
        self.assertEqual(self.features_gggg.GGGG_count, 2)

    def test_seed_base_pairs(self):
        self.assertEqual(self.features.seed_base_pairs, SAMPLE_SEED_0)

    def test_seed_watson_crick_count(self):
        self.assertEqual(self.features.seed_watson_crick_count, 7)

    def test_seed_wobbles_count(self):
        self.assertEqual(self.features.seed_wobbles_count, 0)

    def test_seed_mismatches_count(self):
        self.assertEqual(self.features.seed_mismatches_count, 1)

    def test_base_pair_list(self):
        self.assertEqual(self.features.base_pair_list, SAMPLE_BASE_PAIR_LIST)

    def test_watson_crick_count(self):
        self.assertEqual(self.features.watson_crick_count, 14)

    def test_wobbles_count(self):
        self.assertEqual(self.features.wobbles_count, 3)

    def test_mismatches_count(self):
        self.assertEqual(self.features.mismatches_count, 5)

    def test_wobbles_positions(self):
        self.assertEqual(sorted(self.features.wobbles_positions)[::-1],
                         [(-17, 12), (-22, 17), (-24, 19)])

    def test_mismatches_positions(self):
        self.assertEqual(sorted(self.features.mismatches_positions)[::-1],
                         [(-1, 1), (-18, 13), (-19, 14), (-25, 20), (-26, 21)])

    # count
    def test_hairpin_loop_count(self):
        self.assertEqual(self.features.hairpin_loop_count, 0)
        self.assertEqual(self.features_hpin.hairpin_loop_count, 1)
        self.assertEqual(self.features_junc_t1.hairpin_loop_count, 1)
        self.assertEqual(self.features_junc_t2.hairpin_loop_count, 1)
        self.assertEqual(self.features_sil.hairpin_loop_count, 0)
        self.assertEqual(self.features_asilm.hairpin_loop_count, 0)
        self.assertEqual(self.features_asilmi.hairpin_loop_count, 0)
        self.assertEqual(self.features_bpairmi12.hairpin_loop_count, 2)
        self.assertEqual(self.features_bpairmi15.hairpin_loop_count, 2)

    def test_internal_loop_count(self):
        self.assertEqual(self.features.internal_loop_count, 0)
        self.assertEqual(self.features_hpin.internal_loop_count, 0)
        self.assertEqual(self.features_junc_t1.internal_loop_count, 0)
        self.assertEqual(self.features_junc_t2.internal_loop_count, 0)
        self.assertEqual(self.features_sil.internal_loop_count, 1)
        self.assertEqual(self.features_asilm.internal_loop_count, 1)
        self.assertEqual(self.features_asilmi.internal_loop_count, 1)
        self.assertEqual(self.features_bpairmi12.internal_loop_count, 0)
        self.assertEqual(self.features_bpairmi15.internal_loop_count, 0)

    def test_junction_count(self):
        self.assertEqual(self.features.junction_count, 0)
        self.assertEqual(self.features_hpin.junction_count, 1)
        self.assertEqual(self.features_junc_t1.junction_count, 1)
        self.assertEqual(self.features_junc_t2.junction_count, 1)
        self.assertEqual(self.features_sil.junction_count, 0)
        self.assertEqual(self.features_asilm.junction_count, 0)
        self.assertEqual(self.features_asilmi.junction_count, 0)
        self.assertEqual(self.features_bpairmi12.junction_count, 2)
        self.assertEqual(self.features_bpairmi15.junction_count, 2)

    def test_bulge_count(self):
        self.assertEqual(self.features.bulge_count, 2)
        self.assertEqual(self.features_hpin.bulge_count, 1)
        self.assertEqual(self.features_junc_t1.bulge_count, 0)
        self.assertEqual(self.features_junc_t2.bulge_count, 0)
        self.assertEqual(self.features_sil.bulge_count, 0)
        self.assertEqual(self.features_asilm.bulge_count, 0)
        self.assertEqual(self.features_asilmi.bulge_count, 0)
        self.assertEqual(self.features_bpairmi12.bulge_count, 1)
        self.assertEqual(self.features_bpairmi15.bulge_count, 0)

    def test_stem_count(self):
        self.assertEqual(self.features.stem_count, 3)
        self.assertEqual(self.features_hpin.stem_count, 3)
        self.assertEqual(self.features_junc_t1.stem_count, 3)
        self.assertEqual(self.features_junc_t2.stem_count, 3)
        self.assertEqual(self.features_sil.stem_count, 2)
        self.assertEqual(self.features_asilm.stem_count, 2)
        self.assertEqual(self.features_asilmi.stem_count, 2)
        self.assertEqual(self.features_bpairmi12.stem_count, 5)
        self.assertEqual(self.features_bpairmi15.stem_count, 4)

    # position
    def test_hairpin_loop_positions(self):
        self.assertEqual(self.features.hairpin_loop_positions, [])
        self.assertEqual(self.features_hpin.hairpin_loop_positions, [-25])
        self.assertEqual(self.features_junc_t1.hairpin_loop_positions, [-14])
        self.assertEqual(self.features_junc_t2.hairpin_loop_positions, [-12])
        self.assertEqual(self.features_sil.hairpin_loop_positions, [])
        self.assertEqual(self.features_asilm.hairpin_loop_positions, [])
        self.assertEqual(self.features_asilmi.hairpin_loop_positions, [])
        self.assertEqual(self.features_bpairmi12.hairpin_loop_positions, [11, -21])
        self.assertEqual(self.features_bpairmi15.hairpin_loop_positions, [10, -21])

    def test_internal_loop_positions(self):
        self.assertEqual(self.features.internal_loop_positions, [])
        self.assertEqual(self.features_hpin.internal_loop_positions, [])
        self.assertEqual(self.features_junc_t1.internal_loop_positions, [])
        self.assertEqual(self.features_junc_t2.internal_loop_positions, [])
        self.assertEqual(self.features_sil.internal_loop_positions, [((-3, 3), (-4, 4))])
        self.assertEqual(self.features_asilm.internal_loop_positions, [((-3, 3), (-5, 4))])
        self.assertEqual(self.features_asilmi.internal_loop_positions, [((-3, 3), (-4, 5))])
        self.assertEqual(self.features_bpairmi12.internal_loop_positions, [])
        self.assertEqual(self.features_bpairmi15.internal_loop_positions, [])

# TODO: WHAT DOES THIS MEAN
    def test_junction_positions(self):
        self.assertEqual(self.features.junction_positions, [])
        self.assertEqual(self.features_hpin.junction_positions, [-20])
        self.assertEqual(self.features_junc_t1.junction_positions, [-8])
        self.assertEqual(self.features_junc_t2.junction_positions, [-9])
        self.assertEqual(self.features_sil.junction_positions, [])
        self.assertEqual(self.features_asilm.junction_positions, [])
        self.assertEqual(self.features_asilmi.junction_positions, [])
        self.assertEqual(self.features_bpairmi12.junction_positions, [-13, 8])
        self.assertEqual(self.features_bpairmi15.junction_positions, [-13, 8])

    def test_bulge_positions(self):
        self.assertEqual(self.features.bulge_positions, [-10, -12])
        self.assertEqual(self.features_hpin.bulge_positions, [11])
        self.assertEqual(self.features_junc_t1.bulge_positions, [])
        self.assertEqual(self.features_junc_t2.bulge_positions, [])
        self.assertEqual(self.features_sil.bulge_positions, [])
        self.assertEqual(self.features_asilm.bulge_positions, [])
        self.assertEqual(self.features_asilmi.bulge_positions, [])
        self.assertEqual(self.features_bpairmi12.bulge_positions, [9])
        self.assertEqual(self.features_bpairmi15.bulge_positions, [])

    def test_stem_positions(self):
        self.assertEqual(self.features.stem_positions, [((-1, 1), (-9, 9)), ((-11, 10), (-11, 10)), ((-16, 11), (-27, 22))])
        self.assertEqual(self.features_hpin.stem_positions, [((-1, 1), (-10, 10)), ((-11, 13), (-20, 22)), ((-32, -21), (-29, -24))])
        self.assertEqual(self.features_junc_t1.stem_positions, [((-1, 1), (-8, 8)), ((-20, -9), (-16, -13)), ((-21, 9), (-34, 22))])
        self.assertEqual(self.features_junc_t2.stem_positions, [((-1, 1), (-9, 9)), ((-15, -10), (-14, -11)), ((-16, 11), (-27, 22))])
        self.assertEqual(self.features_sil.stem_positions, [((-1, 1), (-2, 2)), ((-5, 5), (-22, 22))])
        self.assertEqual(self.features_asilm.stem_positions, [((-1, 1), (-2, 2)), ((-6, 5), (-23, 22))])
        self.assertEqual(self.features_asilmi.stem_positions, [((-1, 1), (-2, 2)), ((-5, 6), (-22, 23))])
        self.assertEqual(self.features_bpairmi12.stem_positions, [((10, 14), (10, 14)), ((8, 15), (8, 15)), ((-1, 1), (-7, 7)), ((-8, 16), (-13, 21)), ((-29, -14), (-23, -20))])
        self.assertEqual(self.features_bpairmi15.stem_positions, [((9, 14), (8, 15)), ((-1, 1), (-7, 7)), ((-8, 16), (-13, 21)), ((-29, -14), (-23, -20))])

    # length
    def test_hairpin_loop_lengths(self):
        self.assertEqual(self.features.hairpin_loop_lengths, [])
        self.assertEqual(self.features_hpin.hairpin_loop_lengths, [4])
        self.assertEqual(self.features_junc_t1.hairpin_loop_lengths, [2])
        self.assertEqual(self.features_junc_t2.hairpin_loop_lengths, [2])
        self.assertEqual(self.features_sil.hairpin_loop_lengths, [])
        self.assertEqual(self.features_asilm.hairpin_loop_lengths, [])
        self.assertEqual(self.features_asilmi.hairpin_loop_lengths, [])
        self.assertEqual(self.features_bpairmi12.hairpin_loop_lengths, [3, 2])
        self.assertEqual(self.features_bpairmi15.hairpin_loop_lengths, [4, 2])

    def test_internal_loop_lengths(self):
        self.assertEqual(self.features.internal_loop_lengths, [])
        self.assertEqual(self.features_hpin.internal_loop_lengths, [])
        self.assertEqual(self.features_junc_t1.internal_loop_lengths, [])
        self.assertEqual(self.features_junc_t2.internal_loop_lengths, [])
        self.assertEqual(self.features_sil.internal_loop_lengths, [2])
        self.assertEqual(self.features_asilm.internal_loop_lengths, [(3, 2)])
        self.assertEqual(self.features_asilmi.internal_loop_lengths, [(2, 3)])
        self.assertEqual(self.features_bpairmi12.internal_loop_lengths, [])
        self.assertEqual(self.features_bpairmi15.internal_loop_lengths, [])

# TODO: CHANGE THIS FEATURE TO MORE MEANINGFUL NAME
#       test_junction_stem_count
#    def test_junction_lengths(self):
#        self.assertEqual(self.features.junction_lengths, [])
#        self.assertEqual(self.features_hpin.junction_lengths, [])
#        self.assertEqual(self.features_junc_t1.junction_lengths, [])
#        self.assertEqual(self.features_junc_t2.junction_lengths, [])
#        self.assertEqual(self.features_sil.junction_lengths, [])
#        self.assertEqual(self.features_asilm.junction_lengths, [])
#        self.assertEqual(self.features_asilmi.junction_lengths, [])

    def test_bulge_lengths(self):
        self.assertEqual(self.features.bulge_lengths, [1, 4])
        self.assertEqual(self.features_hpin.bulge_lengths, [2])
        self.assertEqual(self.features_junc_t1.bulge_lengths, [])
        self.assertEqual(self.features_junc_t2.bulge_lengths, [])
        self.assertEqual(self.features_sil.bulge_lengths, [])
        self.assertEqual(self.features_asilm.bulge_lengths, [])
        self.assertEqual(self.features_asilmi.bulge_lengths, [])
        self.assertEqual(self.features_bpairmi12.bulge_lengths, [1])
        self.assertEqual(self.features_bpairmi15.bulge_lengths, [])

    def test_stem_lengths(self):
        self.assertEqual(self.features.stem_lengths, [9, 1, 12])
        self.assertEqual(self.features_hpin.stem_lengths, [10, 10, 4])
        self.assertEqual(self.features_junc_t1.stem_lengths, [8, 5, 14])
        self.assertEqual(self.features_junc_t2.stem_lengths, [9, 2, 12])
        self.assertEqual(self.features_sil.stem_lengths, [2, 18])
        self.assertEqual(self.features_asilm.stem_lengths, [2, 18])
        self.assertEqual(self.features_asilmi.stem_lengths, [2, 18])
        self.assertEqual(self.features_bpairmi12.stem_lengths, [1, 1, 7, 6, 7])
        self.assertEqual(self.features_bpairmi15.stem_lengths, [2, 7, 6, 7])

    # AUCG counts
    def test_hairpin_loop_AUCG_counts(self):
        self.assertEqual(self.features.hairpin_loop_AUCG_counts, [])
        self.assertEqual(self.features_hpin.hairpin_loop_AUCG_counts, [(2, 2, 0, 0)])
        self.assertEqual(self.features_junc_t1.hairpin_loop_AUCG_counts, [(0, 0, 0, 2)])
        self.assertEqual(self.features_junc_t2.hairpin_loop_AUCG_counts, [(0, 0, 2, 0)])
        self.assertEqual(self.features_sil.hairpin_loop_AUCG_counts, [])
        self.assertEqual(self.features_asilm.hairpin_loop_AUCG_counts, [])
        self.assertEqual(self.features_bpairmi12.hairpin_loop_AUCG_counts, [(1, 2, 0, 0), (0, 0, 1, 1)])
        self.assertEqual(self.features_bpairmi15.hairpin_loop_AUCG_counts, [(1, 2, 1, 0), (0, 0, 1, 1)])

    def test_internal_loop_AUCG_counts(self):
        self.assertEqual(self.features.internal_loop_AUCG_counts, [])
        self.assertEqual(self.features_hpin.internal_loop_AUCG_counts, [])
        self.assertEqual(self.features_junc_t1.internal_loop_AUCG_counts, [])
        self.assertEqual(self.features_junc_t2.internal_loop_AUCG_counts, [])
        self.assertEqual(self.features_sil.internal_loop_AUCG_counts, [[(0, 0, 1, 1), (0, 0, 1, 1)]])
        self.assertEqual(self.features_asilm.internal_loop_AUCG_counts, [[(0, 1, 1, 1), (0, 0, 1, 1)]])
        self.assertEqual(self.features_asilmi.internal_loop_AUCG_counts, [[(0, 0, 1, 1), (1, 0, 1, 1)]])
        self.assertEqual(self.features_bpairmi12.internal_loop_AUCG_counts, [])
        self.assertEqual(self.features_bpairmi15.internal_loop_AUCG_counts, [])

    def test_junction_AUCG_unpaired_counts(self):
        self.assertEqual(self.features.junction_AUCG_unpaired_counts, [])
        self.assertEqual(self.features_hpin.junction_AUCG_unpaired_counts, [])
        self.assertEqual(self.features_junc_t1.junction_AUCG_unpaired_counts, [])
        self.assertEqual(self.features_junc_t2.junction_AUCG_unpaired_counts, [(0, 0, 1, 0)])
        self.assertEqual(self.features_sil.junction_AUCG_unpaired_counts, [])
        self.assertEqual(self.features_asilm.junction_AUCG_unpaired_counts, [])
        self.assertEqual(self.features_asilmi.junction_AUCG_unpaired_counts, [])
        self.assertEqual(self.features_bpairmi12.junction_AUCG_unpaired_counts, [])
        self.assertEqual(self.features_bpairmi15.junction_AUCG_unpaired_counts, [])

    def test_bulge_AUCG_counts(self):
        self.assertEqual(self.features.bulge_AUCG_counts, [(1, 0, 0, 0), (3, 0, 0, 1)])
        self.assertEqual(self.features_hpin.bulge_AUCG_counts, [(0, 2, 0, 0)])
        self.assertEqual(self.features_junc_t1.bulge_AUCG_counts, [])
        self.assertEqual(self.features_junc_t2.bulge_AUCG_counts, [])
        self.assertEqual(self.features_sil.bulge_AUCG_counts, [])
        self.assertEqual(self.features_asilm.bulge_AUCG_counts, [])
        self.assertEqual(self.features_asilmi.bulge_AUCG_counts, [])
        self.assertEqual(self.features_bpairmi12.bulge_AUCG_counts, [(0, 1, 0, 0)])
        self.assertEqual(self.features_bpairmi15.bulge_AUCG_counts, [])

    def test_degree_of_complementarity(self):
        self.assertEqual(self.features.degree_of_complementarity,
                         (round(100.0 * 5 / 11), round(100.0 * 1 / 4),
                          round(100.0 * 5 / 5), round(100.0 * 3 / 9)))
        self.assertEqual(self.features_junc_t1.degree_of_complementarity,
                         (round(100.0 * 3 / 4), round(100.0 * 2 / 6),
                          round(100.0 * 5 / 7), round(100.0 * 3 / 18)))


if __name__ == '__main__':
    unittest.main()
```
