## Supplementary File 1 for "Biophysics of microRNA-34a targeting and its influence on down-regulation": context.py

```
# -*- coding: utf-8 -*-

import sys
import os

sys.path.insert(0, os.path.abspath(os.path.join(os.path.dirname(__file__), '..', 'pure_secondary_feature_extraction')))
sys.path.insert(0, os.path.abspath(os.path.join(os.path.dirname(__file__), '..', 'feature_extraction')))

# import sample
import msa_secondary_utils
import rna_utils
```
