## Supplementary File 1 for "Biophysics of microRNA-34a targeting and its influence on down-regulation": generate_name_seq_utf.sh

```
#!/bin/bash

# NOTE: genes such as the following have different target regions, but the same UTR.
# MAP2K1
# MAP2K1-1
# MAP2K1-2
# NOTE: sometimes there are case discrepancies in the raw input files. If no match is found, please check.

for i in `awk '{print $1}' named_sequences`
do 
	name=$i
	basename=`echo $name | cut -d'-' -f1`
	seq=`grep "$i " named_sequences | awk '{print $2}'`
	utr=`grep "$name " named_utrs | awk '{print $2}'`;
	Nutr=`grep "$name " named_utrs | awk '{print $2}' | wc -l`;
	if [ -z "$utr" ];
	then
		utr=`grep "$basename " named_utrs | awk '{print $2}'`;
		Nutr=`grep "$basename " named_utrs | awk '{print $2}' | wc -l`;
	fi
	if [ -z "$utr" ];
	then
		utr=`grep "$name" named_utrs | awk '{print $2}'`;
		Nutr=`grep "$name" named_utrs | awk '{print $2}' | wc -l`;
	fi
	[ $Nutr -gt 1 ] && echo -e "WARNING: Detected name degeneracy! More than one utr found for seq. Please check!"
	[ -z "$utr" ] && echo "WARNING: No utr found for $name"
	echo $name $seq $utr
done
```
