## Supplementary File 1 for "Biophysics of microRNA-34a targeting and its influence on down-regulation": generate_csv_files.sh

```
#!/bin/bash

######################################################################
# This script runs rank_predictions.py in no_pred mode to generate csv files
# containing the features for each and every gene in TRUE_LIST.
######################################################################


bdir=$PWD
csvDir="csv_files"
predScript="/home/dimitri/code/mRNA_miRNA/secondary_structure_prediction/rank_predictions.py"


# clearing/setting up input output directories
[ -d $csvDir ] && rm -r $csvDir 
mkdir $csvDir


# generate predictions
$predScript --no_pred > rank_predictions.out

# move prediction output into appropriate directories
mv *.csv $csvDir 
```
