## Supplementary File 1 for "Biophysics of microRNA-34a targeting and its influence on down-regulation": generate_data_table.sh

```
#!/bin/sh

###########################################################################
# Mark each length within each and every csv file as true (1) or flase (0).
###########################################################################

trueList="TRUE_LIST" # input file containing names of true predictions

# Get the collumn names from the first file
firstFileBase=`head -n 1 $trueList | cut -d'.' -f1`
header=`head -n 1 csv_files/${firstFileBase}.csv`
echo "TRUTH$header" > data_table

# Loop over files
cat $trueList | while read line;
do 
	gene=`echo $line | cut -d'.' -f1`
	fname=${gene}.csv
	if grep -q "${line}" csv_files/$fname; then
		cat csv_files/$fname | sed "s/^${line}.mcff/1/" | sed "s/^.*\.mcff/0/" | grep -v "mRNA" >> data_table

		echo "TRUTH$header" > data_tables/data_$gene
		cat csv_files/$fname | sed "s/^${line}.mcff/1/" | sed "s/^.*\.mcff/0/" | grep -v "mRNA" >> data_tables/data_$gene
		# [ `wc -l data_tables/data_$gene | cut -d' ' -f1` -lt 2 ] && rm data_tables/data_$gene
		echo $gene
	else
		echo "Failed to find ${line}.mcff in file $fname"
	fi
done
```
