## Supplementary File 1 for "Biophysics of microRNA-34a targeting and its influence on down-regulation": generate_training_data.py

```
#!/usr/bin/env python3
# -*- coding: utf-8 -*-

import os
import pandas as pd
import numpy as np
# import itertools


SOURCE_DIR = 'data_tables'


def main():
    if not os.path.isdir(SOURCE_DIR):
        raise ValueError('Failed to find source directory: {}'.format(SOURCE_DIR))
    diffs = generate_diff()
    np.save('feature_diff_matrix.npy', diffs)
    # _diffs = np.load('feature_diff_matrix.npy')


def return_file_paths(source_dir):
    file_paths = [os.path.join(source_dir, f) for f in os.listdir(source_dir)]
    return file_paths


def read_feature_file(file_path):
    featur_DF = pd.read_csv(file_path, header=0, delimiter='\t')
    return featur_DF


def generate_diff(source_dir=SOURCE_DIR):
    """
    Generate a matrix where each row is the following difference:
        features_of_true - features_of_false
    Hence, the optimisation objective will be to maximise number of positive
    values obtained by taking the dot product of this matrix with the weights.
    """
    diff_list = list()
    _data_files = return_file_paths(source_dir)
    for _file in _data_files:
        _feature_DF = read_feature_file(_file)
        _true = _feature_DF[_feature_DF['TRUTH'] == 1].values[0][1:]
        if len(_true) > 0:
            for _false in _feature_DF[_feature_DF['TRUTH'] == 0].values:
                _diff = list(_true - _false[1:])
                diff_list.append(_diff)
    return np.array(diff_list)


if __name__ == '__main__':
    main()
```
