## Supplementary File 1 for "Biophysics of microRNA-34a targeting and its influence on down-regulation": Optimize_RNA_duplex_construct_v1.5.py

```
#!/usr/bin/env python3.6
# -*- coding: utf-8 -*-

# ************************************************************************** #
# ||    ~~ Script for optimizing mRNA-miRNA construct sequences v1.0 ~~   || #
# ||                         Written by Noah Hopkins                      || #
# ||       Katja Petzold's group Karolinska Institutet, December 2016     || #
# ************************************************************************** #


# **********------------------------------------------------------********** #
# |                              ~~ IMPORTS ~~                             | #
# **********------------------------------------------------------********** #

import contextlib

import subprocess
import collections
import urllib.request
import urllib.parse
import urllib.error
import sys
import os
import re
# import time

# NOTE: The 'numpy' module might also be imported!
# NumPy is not part of the standard python distribution. It is used in the
# script to show all the relevant output data directly in the console in a
# structured and neat way. The 'numpy' import can be prevented by setting
# 'output_data_table' to 'False' in the options section (option #11).
#
# It is highly recommended to install Anaconda before using this script.
# Anaconda is a scientific Python distribution which automatically installs
# all needed Python modules related to scientific work (NumPy, SciPy,
# MatPlotLib, Jupyter Notebook etc) and keeps them updated.
#
# Anaconda can be downloaded here: https://www.continuum.io/downloads


# **********------------------------------------------------------********** #
# |                               ~~ INFO ~~                               | #
# **********------------------------------------------------------********** #

# Script dependencies: Mc-FlashFold-vXX, CONTRAfold's plot_rna wrapper (web
# service) and NumPy (Python module, optional).

# For a usage guide and mcff installation guide, see the README.txt file
# included with the script.

# N.B: don't forget to set the tables directory before running the script!


# **********------------------------------------------------------********** #
# |                      ~~ OPTIONS AND INPUT DATA ~~                      | #
# **********------------------------------------------------------********** #

# Path to mcff executable
MCFF = "/home/dimitri/programs/MCFOLD/MC-Flashfold-v34/src/mcff"

# seq_name = 'TEST'
seq_name = sys.argv[1]
# 1. [str] Name of the sequence / mRNA / Gene.

# FILEDIR = '/home/noah/Mc-Flashfold-v34/Temp test'
FILEDIR = 'OUT'
if not os.path.exists(FILEDIR):
    os.makedirs(FILEDIR)
# 2. [str] Output file directory.

# mRNA_sequence = 'UGUUCUUUAUGCAUAAAACACCCAGCUAGGACCAUUACUGCCAGAGAAAAAAAUC'
mRNA_sequence = sys.argv[3]
# 3. [str] The whole 5'->3' mRNA sequence, in (+) sense DNA or RNA form.

# mRNA_seq_target_region = 'CCAGCUAGGACCAUUACUGCCA'
mRNA_seq_target_region = sys.argv[2]
# 4. [str] The 5'->3' miRNA target region (a sub sequence of 'mRNA_sequence').

added_left_nt = 8
added_right_nt = 0
# 5. [int] The number of additional nucleotides (from the mRNA sequence) to
# try on left and right side of the target region.

loop = 'UUCG'
# 6. [str] The joining loop (should normally be 'UUCG'), 5'->3'.

miRNA = 'UGGCAGUGUCUUAGCUGGUUGU'
# 7. [str] The miRNA (that targets the chosen mRNA), 5'->3'.

add_GG = (False, 2,)
# 8. [(bool, int)] Set first tuple element to 'True' if you want to add Gs to
# the beginning of all sequences given to mcff. Set the second element to
# either 1 or 2 depending on how many Gs to add.

mode = '-ft'
mode_value = 50
# For -ft mode, this sets the number of structures returned
# 9. [str], [int] The '-ft' (floating threshold) mode allows for the specifi-
# cation of how many structures one wants to find (approximately, may vary).
# Information about the available modes can be found in 'flashfold-f34.pdf'
# (page 5-7, good to read).

# 10.0 [bool] Set to False if plotting is not required.
generate_plot = True

file_format = 'pdf'  # Can be set to 'pdf', 'ct', 'ps' or 'jpg'.
# 10.1 [str] Figure format.

output_data_table = True
# 11. [bool] To show all the relevant output data directly in the console set
# 'output_data_table' to 'True'.

# NOTE: If this is set to true, the module 'numpy' will be imported. It is
# recommended to install Anaconda before using this script.
# Anaconda can be downloaded here: https://www.continuum.io/downloads

# -------------------------------------------------------------------------- #
# 12. To apply a mask, set 'msk_mode' to any of the following options (other-
# wise set it to 'None'):
#
# -mask (-m)                     --> Balanced, full mask.
# -output_mask (-om)             --> Balanced, only apply mask to output.
# -unbalanced_mask (-um)         --> Unbalanced, full mask.
# -output_unbalanced_mask (-oum) --> Unbalanced, only apply mask to output.
#  None                          --> No mask applied.

mask = 'xx(x(((((xx(((xxx(x((((((((x(..)x))))))'
msk_mode = None  # '-um'

# INFO: A "mask" can be used to (a), either filter the output OR (b), const-
# rain the search of possible structures by the algorithm (will influence the
# output). Furthermore, a mask can be either balanced or unbalanced. Balanced
# implies that all nt-nt bonds are specified. An unbalanced mask, on the other
# hand, specifies the pairing status of each individual nucleotide without
# specifying to which nucleotide it is pairing. A balanced mask is simply a
# dot-bracket structure with 'x's where the pairing is unknown / unspecified.
# For complete information about masks, read the 'flashfold-f34.pdf' (pages
# 8-10) included in the mcff installation package.
# -------------------------------------------------------------------------- #

forced_figure_name = None
# 13. [str or NoneType] To set a custom name (fig title) to the figures
# rendered, set 'forced_figure_name' to the desired (str) name (otherwise set
# it to 'None').

verbosity = 3
# 14. [int] Verbosity level can be set between 0 and 5.
# Recommended setting is '3'.

# INFO: Verbosity is the amount of text/information displayed in the console
# while running the script. The verbosity level can be set between 0 and 5.
# Zero meaning no text shown (black/empty terminal) and 5 meaning all possible
# information about what is happening is shown. For verbosity levels 0-3 a
# progress bar is shown (instead of printing everything that is happening).
# This progress bar can be turned off if you so desire (see next option).

progress_bar = True
# 15. [bool] Progress bar used for verbosity levels 0-3. To disable or enable
# the progress bar, set 'progress_bar' to 'False' or 'True', resp.

TABLES_DIR = '/home/dimitri/programs/MCFOLD/MC-Flashfold-v34/tables'

# 16. [str] Sets the 'tables' directory.
# INFO: For mcff to work, directory of the 'tables' folder must be specified.


# **********------------------------------------------------------********** #
# |                         ~~ DEFINING FUNCTIONS ~~                       | #
# **********------------------------------------------------------********** #


def dna2rna(sequence):
    r"""Return DNA string as RNA string in caps.

    Replaces all 'T' with 'U' in 'sequence'.
    Turns lower case letters into upper case.

    Parameters
    ----------
    sequence : str
        The nucleic acid sequence to transform.
        Can ONLY contain a str with the following characters in either
        upper or lower case:
            A, G, C, T, U

    Returns
    -------
    rna_sequence : str
        Returns 'sequence' in upper case, converted into RNA form.

    Examples
    --------

    >>> dna2rna("ATTgc")
    'AUUGC'

    """
    nts_lower = ('a', 'g', 'c', 't', 'u',)
    nts_upper = ('A', 'G', 'C', 'T', 'U',)
    for nt_lw, nt_up in zip(nts_lower, nts_upper):
        if nt_lw in sequence:
            sequence = sequence.replace(nt_lw, nt_up)
    rna_sequence = sequence.replace('T', 'U')
    return rna_sequence


def update_progress(job_title, progress):
    r"""Progress bar for visualisation of the progress of a task.

    Prints the current progress, and then flushes.
    If progress is 100%, print a completion message.

    Parameters
    ----------

    job_title : str
        The title of the task being performed and tracked.
    progress : int or float
        A number between 0 and 1. New status of the progressbar.
        '1' means 100% (i.e. task complete).

    Notes
    -----

    The variable 'length' (defined inside the function) should be
    set to the desired length of the progress bar.

    Examples
    --------

    >>> task = "Extracting data"
    >>> update_progress(task, 0.5)
    'Extracting data: [##########----------] 50.0%'

    """
    length = 20
    block = int(round(length * progress))
    msg = BOLD + '\r{0}: [{1}] {2}%'.format(job_title, '#' * block + '-' *
                                            (length - block) + END,
                                            round(progress * 100, 2))
    if progress >= 1:
        msg += '\r\n'
    sys.stdout.write(msg)
    sys.stdout.flush()


def parse_mcff_string(mcff_string):
    '''Parse the lines of an mcff file.

    The only checking that is performed at this level is for the existence of
    the values to construct a dictionary as detailed below.

    Args:
        mcff_string <str> The content of an mcff file in string format.

    Returns:
        mcff_od <OrderedDict> An OrderedDict with the content of the
        mcff_string in the form of key value pairs as follows:
            'name': name <string>,
            'primary': sequence <string>,
            'secondaries': list of secondary structures (dot bracket notation) <list of strings>,
            'energies': list of energies <list of floats>.
    '''
    _name = None
    _primary = None
    _secondaries = list()
    _energies = list()
    # shorthand notation indicating for secondary structure
    # indicating the number of loops.
    _shorthand = list()
    for line in mcff_string.split('\n'):
        line = line.strip()
        if re.match(r'>', line) and not _name:
            _name = line[1:]
        elif re.match(r'[A-Z\d]+$', line):
            _primary = line
        elif re.match(r'\(', line) or re.match(r'\.', line):
            line = line.split()
            if len(line) == 3:
                _secondaries.append(line[0])
                _energies.append(float(line[1]))
                _shorthand.append(line[2])
        # else:
        #     print('OTHER: {}'.format(line))
    # print(_name, _primary, _secondaries, _energies)
    if not (_name and _primary and _secondaries and _energies):
        raise ValueError('encountered incomplete mcff file')
    kv_list = [
        ('name', _name), ('primary', _primary), ('secondaries', _secondaries),
        ('energies', _energies)]
    mcff_od = collections.OrderedDict(kv_list)
    return mcff_od


BOLD = '\033[1m'
UNDERLINE = '\033[4m'
RED = '\033[91m'
END = '\033[0m'

# If these doesn't work and print out literally, import the module colorama:
import_colorama = False

if import_colorama:
    from colorama import init
    init()


# **********------------------------------------------------------********** #
# |                           ~~ START OF SCRIPT ~~                        | #
# **********------------------------------------------------------********** #

# If 'output_data_table' is set to 'True', import module 'numpy' and increase
# the line width to make room for the output data table.
if output_data_table:
    import numpy as np

    np.set_printoptions(threshold=np.inf, linewidth=1000)

# If sequences in DNA form, convert to RNA.
RNA_mRNA_sequence = dna2rna(mRNA_sequence)
mRNA_target_region = dna2rna(mRNA_seq_target_region)
loop = dna2rna(loop)

# Calculate the length of the target region in the mRNA sequence.
l = len(mRNA_target_region)

# Find the location (index) of the target region inside the mRNA sequence.
nt_reference = RNA_mRNA_sequence.find(mRNA_target_region)

# Calculate the total number of mcff output files (also the number of unique
# sequences).
nr_of_outputs = (added_right_nt + 1) * (added_left_nt + 1)

# Removes the '-' character from the mode flag. E.g. '-ft' becomes 'ft'.
# Used when creating the filename.
mode_str = mode.replace("-", "")

mode_value = str(mode_value)

print()  # Padding.


# Loop for; assembly of mcff command, call to mcff with command and saving of
# output into .mcff file.
output_list = []
mcff_msg_list = []
all_output_files = []
cmd_counter = 1
x = 0
while x < added_right_nt + 1:
    y = 0
    while y < added_left_nt + 1:
        left = added_left_nt - y
        right = added_right_nt - x

        # Creating the mcff input sequence "(GG/G) + mRNA + loop + miRNA":
        if add_GG == (True, 2,):
            RNAsequence = ('GG' + RNA_mRNA_sequence[nt_reference - left:
                                                    nt_reference + l
                                                    + added_right_nt]
                           + loop + miRNA)
        elif add_GG == (True, 1,):
            RNAsequence = ('G' + RNA_mRNA_sequence[nt_reference - left:
                                                   nt_reference + l +
                                                   added_right_nt] + loop + miRNA)
        else:
            RNAsequence = (RNA_mRNA_sequence[nt_reference - left:
                                             nt_reference + l
                                             + added_right_nt]
                           + loop + miRNA)

        # Creating the name of the .mcff file:
        if add_GG == (True, 1):
            filename = (seq_name + '.' + mode_str
                        + mode_value + '.' + 'L' + str(left)
                        + '.' + 'R' + str(right) + '.G' + '.mcff')
        elif add_GG == (True, 2):
            filename = (seq_name + '.' + mode_str
                        + mode_value + '.' + 'L' + str(left)
                        + '.' + 'R' + str(right) + '.GG' + '.mcff')
        else:
            filename = (seq_name + '.' + mode_str
                        + mode_value + '.' + 'L' + str(left)
                        + '.' + 'R' + str(right) + '.mcff')

        output_file = os.path.join(FILEDIR, filename)
        all_output_files.append(output_file)

        # Assembly of the mcff command:
        if msk_mode is None:
            # cmd = ['./mcff', '-s', RNAsequence, '-n',
            cmd = [MCFF, '-s', RNAsequence, '-n',
                   filename, '-times', '-v2', mode,
                   mode_value, '-tables', TABLES_DIR]
        else:
            # cmd = ['./mcff', '-s', RNAsequence, msk_mode,
            cmd = [MCFF, '-s', RNAsequence, msk_mode,
                   mask, '-n', filename, '-times', '-v2',
                   mode, mode_value, '-tables', TABLES_DIR]

        # Calling mcff (don't know if 'universal_newlines' is needed):

        # p = subprocess.Popen(cmd, stdout=subprocess.PIPE, stderr=subprocess.PIPE, cwd='/usr/local/bin', universal_newlines=True)

        p = subprocess.Popen(cmd, stdout=subprocess.PIPE, stderr=subprocess.PIPE)
        out, err = p.communicate()
        out = out.decode()
        err = err.decode()

        if p.returncode != 0:
            err_msg = '{}. Code: {}'.format(err.strip(), p.returncode)
            raise RuntimeError(err_msg)
        # Append the output into lists:

        output_list.append(out)
        if verbosity >= 3:
            mcff_msg_list.append(err)
        output = out
        mcff_msg = err
        p.wait()

        # Print information to host (depending on verbosity level):
        if verbosity >= 2:
            print((BOLD + '## -----[mcff Call %s]----- ##' % cmd_counter + '\n' + END))
        if verbosity >= 2:
            print(('COMMAND:\n' + ' '.join(cmd) + '\n'))
        if verbosity >= 3:
            print(mcff_msg)
        if verbosity >= 5:
            print(('OUTPUT:', output))

        # Save output to file:
        with open(output_file, "w") as new_text_file:
            new_text_file.write(output)

        y += 1
        cmd_counter += 1
    x += 1
if verbosity >= 2:
    print((BOLD + '## ---------[Done]--------- ##' + '\n' + END))


if generate_plot:
    # Loop for;
    # 1. assembly of plot_rna URLs,
    # 2. retrieval of html page source,
    # 3. scanning for URLs to figures,
    # 4. downloading of figures.
    #
    # INFO: URL goes to CONTRAfold's plot_rna wrapper (version 'Nov 13 2014').
    # Sample URL (below). Replace the words with brackets (incl. the brackets them
    # selves) with the actual sequence name, sequence, dot-bracket structure and
    # an arbitrary identifier number:
    # http://www.major.iric.ca/cgi-bin/2DRender/render.cgi?structure=[Seq name]|
    # [Seq]|[DB struct]&structno=[Struct nr]
    # for i in range(0, len(list_of_tuples)):
    size = len(output_list) * int(mode_value)
    cnt = 0
    for mcff_output in output_list:
        mcffOD = parse_mcff_string(mcff_output)
        RNAseq = mcffOD['primary']
        filenm = mcffOD['name']

        for i, dotbr in enumerate(mcffOD['secondaries']):
            cnt += 1
            # Assembly of plot_rna URL:
            if forced_figure_name is not None:
                # noinspection PyTypeChecker
                URL_plot_rna = ('http://www.major.iric.ca/cgi-bin/2DRender/render.cgi'
                                + '?structure=%3E' + forced_figure_name + '|'
                                + RNAseq + '|' + dotbr + '&structno=' + str(i))
            else:
                URL_plot_rna = ('http://www.major.iric.ca/cgi-bin/2DRender/render.cgi'
                                + '?structure=%3E' + filenm + '|'
                                + RNAseq + '|' + dotbr + '&structno=' + str(i))
            # Assemble name of temp .html file:
            output_file = os.path.join(FILEDIR, filenm.replace('.mcff', '') + ".html")
            # Printing information to host (depending on verbosity level):
            if verbosity >= 4 and i == 0:
                print(BOLD + '\n' + '## ------[Sequence %s]------ ##' % (i) + END)
            if verbosity >= 4:
                print('\n' + "[%s] URL to CONTRAfold's plot_rna wrapper;" % i + '\n' + UNDERLINE + URL_plot_rna + END)
            # A 'context manager' that closes the file upon completion of the block:
            with contextlib.closing(urllib.request.urlopen(URL_plot_rna)) as temp:
                substring_list = temp.read().split()
                # Do stuff with each substring:
                for substring in substring_list:
                    substring = substring.decode()
                    # See if the word 'http' is present in substring and if so in
                    # what position/index it can be found:
                    start = substring.find('http')
                    # If 'http' is not present, "substring.find('http')" returns
                    # '-1'. If it does, skip the rest of the current iteration of
                    # the loop, and continue directly with the NEXT iteration.
                    if start == -1:
                        continue  # ...with next iteration.

                    # check for existence of desired file format within url string.
                    if substring.find('.' + file_format) != -1:
                        end = substring.find('.' + file_format)
                        url_string = substring
                    else:
                        continue
                    figure_url = url_string[start:end + len(file_format) + 1]
                    # Assembly of figure name:
                    picture_name = (output_file.replace('.html', '') + '.DB_' + str(i).zfill(len(mode_value)) + "." + file_format)
                    # Open URL and retrieve figure:
                    urllib_opener = urllib.request.URLopener()
                    urllib_opener.retrieve(figure_url, picture_name)
                    # If verbosity level is over 4, print the name of the figure:
                    if verbosity >= 4:
                        print(('Creating:', picture_name))
            if progress_bar and verbosity <= 3:
                # noinspection PyUnboundLocalVariable
                update_progress('Progress', float(cnt) / size)

# # Printing information to host (depending on verbosity level):
# if verbosity >= 1:
#     print((BOLD + 'Total number of sequences submitted: ' + END + str(len(list_of_tuples))))
#     print(BOLD)
#     if verbosity >= 3:
#         print('\n' + 'Number of dot-bracket structures per sequence (per .mc' + 'ff file):')
#         print('Header: [Seq1, Seq2, Seq3, ...]')
#         print(END, '\r', nr_of_structs)
#
# if output_data_table:
#     print((BOLD + '\n' + 'Full output data table: ' + '\n'
#           + "Header: ['Filename' 'Sequence' 'MFE' 'Threshold' "
#           + "'Dot-bracket struct' 'Energy' 'Loops']"
#           + '\n' + END), np.array(list_of_tuples), '\n')
#
# if verbosity >= 2:
#     print((BOLD + '\n' + '## ------------------------ ##' + END))
#
# if verbosity >= 2:
#     print(BOLD + '\n' 'For each sequence:' + '\n' + "--> Call CONTRAf old's plot_rna wrapper.")
#     print('--> Retrieve html page and find figure URLs.')
#     print('--> Render and download figures for each dot-bracket structure.' + '\n' + END)
#
#
# # If loop is complete, set progress bar to complete:
# if progress_bar and verbosity <= 3:
#     update_progress('Progress', 1)
#
# # Print completion message to host (depending on verbosity level):
# if verbosity >= 1:
#     print((BOLD + '\n## ---------[Done]--------- ##\n' + END))
#

# **********------------------------------------------------------********** #
# |                          ~~ NOTES & TODOs ~~                           | #
# **********------------------------------------------------------********** #


# TODO(Noah): Look into local usage of CONTRAFOLD's plot_rna

# TODO(Noah): Add execfile() and multithreading tutorial to the README.txt

# TODO(Noah): mcff dot- & arc plot option (see sample command beneath).
#
#   Sample command for generating dot- & arc plots:
#       ./db2cm -f ~/Mc-Flashfold-v34/test_seq.ft5.mcff -size 0.1 -xdb 100
#
#   It might, however, be better to do this in a separate script.
```
