## Supplementary File 1 for "Biophysics of microRNA-34a targeting and its influence on down-regulation": rank_predictions.py

```
#!/usr/bin/env python3
# -*- coding: utf-8 -*-
"""
Usage:
    rank_predictions.py    (-h | --help)
    rank_predictions.py    --version
    rank_predictions.py    <mcff_dir> [--no_pred] [--max_depth=<int>]

Options:
    -h --help           Show this screen.
    --version           Show the version.
    --no_pred           Do not output the prediction.
    --max_depth=<int>   The maximum number of secondary structure predictions (starting with the mfe) to consider [default: 5].

Purpose of this script:
    This script analyses the content of each of the mcff files (secondary
    structure predictions) corresponding to each and every gene and returns the
    length and energy rank of the most likely secondary structure prediction.

About the input data:
    For each gene, there are several secondary structure predictions as
    generated by Optimize_RNA_duplex_construct_v1.5.py. They differ in the
    number of nucleotides that are added to the left of the mRNA target region.
    Each set of predictions is in a separate mcff file. Each mcff file contains
    several (usually 50) predicted secondary structures.


Features used for the assessment of the secondary structure predictions:

    Dealbreakers:
        - miRNA side: self base pairing.
        - miRNA side: unpaired.
        - short stem: the existence of a stem of length 1 (e.g.\ single base pair) at 5'/3' end.

    Negative features (lower value is better):
        - mRNA self basepairing: self base pairing of the mRNA.
        - mRNA unpaired: unpaired bases on of mRNA.
        - stem short count: the number of stems of length 1.
        - energy difference: the difference in energy between the chosen
          structure and the structure of minimum energy.

    Positive features (higher value is better):
        - mRNA length: the length of the mRNA sequence
        - stem stability: This score that is derived from number of stems that
        end in WC base pairs as a proportion of the total number of stems.
        Wobbles and mismatches are both penalised; the latter is subject to
        twice the penalty as compared to the former.
        - degree of complementarity: the average complementarity of the bases
        in the mRNA. Please see the README of the feature extraction module for
        the formal definition of the degree of complementarity.
        - conservation: the degree to which the secondary structure of the
        structure in question is conserved among the secondary structure
        predictions of the same mRNA length.
"""

import os
# import sys
# import re
import numpy as np
import pandas as pd
from collections import OrderedDict
from docopt import docopt
from context import rna_utils as rna


__author__ = 'Dimitri Schritt'
__copyright__ = 'Copyright (C) Karolinska Institutet - All Rights Reserved 2017'

WEIGHTS_FILE = os.path.abspath(os.path.join(os.path.dirname(__file__), 'saved_models', 'weight_vector'))


'''
NOTES:

    Reminder: 5' mRNA UUCG miRNA 3'

    Possible alternate approach:

    Download solved structures, convert to secondary structure, run sequences
    through pipeline, try to set up ML solution for the prediction of what is
    the most likely model.

    TODO (Katja/Ileana):
        check training data to make sure that correct length was chosen based
        upon biological viability rather than ease of synthesis
'''


def main():
    file_OD = _gather_inputs(IN_DIR)
    max_depth = int(arguments['--max_depth'])
    # for gene in [i for i in file_OD.keys() if i.startswith('MET-3')]:
    # for gene in [i for i in file_OD.keys() if i.startswith('GALNT7')]:
    # problematic_genes = ['CDK6-2', 'BIRC5', 'LDHA', 'MDM4-1_intheCDS', 'NANOG']
    # failed_predictions = ['AXL1', 'BCL2', 'CCND1', 'CD24', 'DLL1-2', 'HK2',
    #                       'NOTCH1', 'PDK1-1', 'PIK3R2-3', 'SMAD4', 'ULBP2',
    #                       'WNT1', 'ZNF281']
    # for gene in [i for i in file_OD.keys() if i in failed_predictions]:
    # for gene in [i for i in file_OD.keys() if i.startswith('BCL2')]:
    # for gene in [i for i in file_OD.keys() if i.startswith('ACSL1')]:
    # for gene in [i for i in file_OD.keys() if (i not in problematic_genes)]:
    for gene in file_OD.keys():
        print(gene)
        file_feature_OD = _return_feature_objects_OD(file_OD[gene], max_depth)
        score_DF_raw = apply_assessment_criteria(file_feature_OD, max_depth)
        score_DF = score_DF_raw.dropna()
        # print(score_DF.T)
        normalisedDF = apply_normalisation(score_DF)
        # print(normalisedDF)
        conservationOD = return_conservationOD(normalisedDF, file_feature_OD)
        # for i in conservationOD.items():
        #     print(i)
        normalisedDF['conservation'] = pd.Series(index=list(conservationOD.keys()), data=list(conservationOD.values()))
        # columns_ignored = ['index', 'energy_rank', 'mRNA_unpaired_GGGG', 'mRNA_5prime_GG']
        columns_ignored = ['energy_rank', 'mRNA_unpaired_GGGG', 'mRNA_5prime_GG']
        df_for_assessment = normalisedDF.drop(columns_ignored, inplace=False, axis=1)
        format_output(df_for_assessment)
        if arguments['--no_pred']:
            df_for_assessment.to_csv('{}.csv'.format(gene), sep='\t')
            continue
        prob_true = apply_weights(df_for_assessment)
        df_for_assessment['score'] = pd.Series(index=df_for_assessment.index.values, data=prob_true)
        df_for_assessment.sort_values(by=['score'], ascending=False, inplace=True)
        df_for_assessment = df_for_assessment.round(2)
        print(df_for_assessment)
        df_for_assessment.to_csv('{}_pred.csv'.format(gene), sep='\t')
        # normalisedDF.to_latex('{}_pred.tex'.format(gene))


def format_output(featureDF):
    '''
    If featureDF contains a column called index this column is used to change
    the index column to include the index. The elements of the index column
    would change from something like ACSL1.ft50.L8.R0.mcff to something like
    ACSL1.ft50.L8.R0_0. The index column would
    then be removed.
    '''
    if 'index' in featureDF.columns.values:
        old_index = [i.replace('.mcff', '') for i in featureDF.index.values]
        pred_index = featureDF['index'].values
        new_index = [i + '_%i.mcff' % j for i, j in zip(old_index, pred_index)]
        featureDF.index = new_index
        featureDF.drop(['index'], inplace=True, axis=1)
    return featureDF


def apply_weights(featureDF, weights_file=WEIGHTS_FILE):
    weights = np.loadtxt(weights_file)
    scores = np.dot(featureDF.values, weights)
    return scores


def return_conservationOD(score_DF, file_feature_OD):
    _files = score_DF.index.values
    # print(score_DF.index.values)
    # print(file_feature_OD.keys())
    _secondariesOD = OrderedDict()
    _conservationOD = OrderedDict()
    for _file in _files:
        _en_rank = score_DF.loc[_file, 'energy_rank']
        _secondary = file_feature_OD[_file]['F_OD'][_en_rank].secondary
        _secondariesOD[_file] = _secondary
    for _file in _files:
        _secondary = _secondariesOD[_file]
        _secondaries = _secondariesOD.values()
        # _rel_sim = rna.return_relative_similarity(_secondary, _secondaries)
        _rel_sim = rna.return_relative_similarity_pairwiseALIGN(_secondary, _secondaries)
        _conservationOD[_file] = _rel_sim
    return _conservationOD


def apply_assessment_criteria(file_feature_OD, max_depth):
    '''
    Input: file_feature_OD
    Output: data frame with assessment of each length.
    '''
    # TODO: complete doc string above.
    criteria = ['energy_rank',
                'energy_diff',
                'mRNA_self_basepairing',
                'mRNA_unpaired_GGGG',
                'mRNA_unpaired',
                'mRNA_5prime_GG',
                'mRNA_length',
                'stem_short_count',
                'stem_stability',
                'degree_of_complementarity',
                'index']

    scoreDF = pd.DataFrame(index=file_feature_OD.keys(), columns=criteria, data=np.nan)
    scoreDF.index.name = 'GENE'
    for _file in file_feature_OD.keys():
        print('#' * 40)
        print(_file)
        print(file_feature_OD[_file]['F_OD'].keys())
        print('#' * 40)
        for en_rank in file_feature_OD[_file]['F_OD'].keys():
            _index = file_feature_OD[_file]['I_OD'][en_rank]
            print(('Examining {}, prediction number before/after redundancy'
                   ' filtering {}/{}.'.format(_file, en_rank, _index)))
            _Secondary = rna.Secondary(file_feature_OD[_file]['F_OD'][en_rank].secondary)

            _fastaFeatures = file_feature_OD[_file]['F_OD'][en_rank]
            _SelfBasepairing = rna.SelfBasepairing(_fastaFeatures)
            # dealbreaker: unpaired or self basepairing on miRNA side.
            if (_Secondary.count_unpaired_miRNA > 0) or _SelfBasepairing.miRNA:
                print((rna.Colour.WARNING + 'Dealbreaker for {} in prediction '
                       'number {}: has unpaired miRNA or self basepairing on '
                       'miRNA side'.format(_file, en_rank) + rna.Colour.ENDC))
            # dealbreaker: short stem at end
            elif _Secondary.short_stem_at_end:
                print((rna.Colour.WARNING + 'Dealbreaker for {} in prediction '
                       'number {}: has short stem at end'.format(_file, en_rank) + rna.Colour.ENDC))

            else:
                # try to find better one lower down, else take top one
                if (_Secondary.count_unpaired_mRNA > 0) and _SelfBasepairing.mRNA and en_rank < max_depth:
                    print((rna.Colour.WARNING + 'Looking for lower energy structure with the '
                           'following: self basepairing and unpaired bases on '
                           'mRNA side' + rna.Colour.ENDC))
                else:
                    # if all secondary structures have been tried and yet the
                    # desired property of no (self basepairing and unpaired
                    # bases on mRNA side) then return to first (lowest) energy
                    # structure
                    if (_Secondary.count_unpaired_mRNA > 0) and _SelfBasepairing.mRNA and (en_rank == max_depth):
                        en_rank = 0
                        _Secondary = rna.Secondary(file_feature_OD[_file]['F_OD'][en_rank].secondary)
                    _fastaFeatures = file_feature_OD[_file]['F_OD'][en_rank]
                    _mcffFeatures = file_feature_OD[_file]['M']
                    _mRNAFivePrimem = rna.mRNAFivePrimem(_fastaFeatures)
                    print(rna.Colour.GREEN + 'Settled for {} at prediction number {}.'.format(_file, en_rank) + rna.Colour.ENDC)
                    scoreDF.loc[_file, 'energy_rank'] = en_rank
                    scoreDF.loc[_file, 'energy_diff'] = _mcffFeatures.energy_difference(0, en_rank)
                    scoreDF.loc[_file, 'mRNA_self_basepairing'] = _SelfBasepairing.mRNA
                    scoreDF.loc[_file, 'mRNA_unpaired'] = _Secondary.count_unpaired_mRNA
                    scoreDF.loc[_file, 'mRNA_5prime_GG'] = _mRNAFivePrimem.GG
                    scoreDF.loc[_file, 'mRNA_unpaired_GGGG'] = _mRNAFivePrimem.GGGG * (_Secondary.count_unpaired_mRNA >= 4)

                    scoreDF.loc[_file, 'mRNA_length'] = _fastaFeatures.mRNA_length
                    scoreDF.loc[_file, 'stem_short_count'] = rna.stem_short_count(_fastaFeatures)
                    scoreDF.loc[_file, 'stem_stability'] = rna.stem_stability(_fastaFeatures)
                    scoreDF.loc[_file, 'degree_of_complementarity'] = round(np.mean(_fastaFeatures.degree_of_complementarity))
                    scoreDF.loc[_file, 'index'] = _index
                    break
    return scoreDF


def apply_normalisation(scoreDF):
    '''
    Normalise all criteria to be between 0 and 1.
    '''
    df = scoreDF.copy().dropna()
    # df['energy_diff'] = np.exp(-df['energy_diff'])
    df['stem_stability'] = df['stem_stability'] / 100.0
    df['degree_of_complementarity'] = df['degree_of_complementarity'] / 100.0
    # df['mRNA_length'] = rna.normalise(df['mRNA_length'], 0, df['mRNA_length'].max())
    criteria = [
        # 'energy_rank',
        # 'energy_diff',
        # 'mRNA_self_basepairing',
        # 'mRNA_unpaired_GGGG',
        # 'mRNA_unpaired',
        # 'mRNA_5prime_GG',
        # 'stem_short_count',
        # 'stem_stability',
        # 'degree_of_complementarity',
        'mRNA_length']
    for criterion in criteria:
        df[criterion] = rna.normalise(df[criterion], 0, df[criterion].max())
    return df


def _gather_inputs(file_path):
    '''
    Input: a file path
    Output: an OrderedDict with key value pairs of the form
            gene: [all files corresponding to that gene]
    '''
    file_list = [i for i in os.listdir(file_path) if i.endswith('.mcff')]
    gene_list = np.unique(sorted([i.split('.')[0] for i in file_list]))
    file_OD = OrderedDict()
    for gene in gene_list:
        file_OD[gene] = sorted([i for i in file_list if i.startswith(gene + '.')])
    return file_OD


def _return_feature_objects_OD(mcff_file_list, max_depth=None):
    '''
    Input: mcff_file_list
        This could be any list of mcff files, but it is intended to be
        a list of files for a given gene where each file corresponds to a
        different length of the mRNA.
    Input: max_depth
        This is the number of secondary structure predictions to use. For
        example, if max_depth = 5, then FastaFeatures will be computed for up
        to 5 secondary secondary structures for each mcff file.
    Output: file_object_OD
        This stores the following:
            M for each and every file
            F_OD an OrderedDict of Fs, one for each and every secondary structure prediction in each and every file
            I_OD an OrderedDict of Is, the indices of every secondary structure prediction for each and every file
    '''
    file_object_OD = OrderedDict()
    # for _file in [i for i in mcff_file_list if i in failed_ar]:
    # for _file in [i for i in mcff_file_list if i.startswith('CCND3.ft50.L7.R0.mcff')]:
    # for _file in [i for i in mcff_file_list if i.startswith('CCND3')]:
    for _file in mcff_file_list:
        print('')
        print('Commencing redundancy filtering and feature generation for content of file: {}'.format(_file))
        mcff_file_path = os.path.join(IN_DIR, _file)
        mcff_raw = rna.parse_mcff_file(mcff_file_path)
        mcff_unique = rna.return_unique_mcff(mcff_raw)
        # mcff_lengths.append(len(mcff_unique['secondaries']))
        # max_mcff_length = min(mcff_lengths)
        try:
            rna.validate_mcff(mcff_unique)
            rna.validate_primary(mcff_unique['primary'])
        except Exception as e:
            print('')
            print(rna.Colour.RED + _file + rna.Colour.ENDC)
            print(e)
        else:
            M = rna.McffFeatures(mcff_unique)
            F_OD = OrderedDict()
            I_OD = OrderedDict()
            file_object_OD[_file] = OrderedDict([('M', M), ('F_OD', F_OD), ('I_OD', I_OD)])
            if max_depth:
                _max = min(max_depth, len(M.mcff_fastas))
            else:
                _max = len(M.mcff_fastas)
            # print(M.mcff['energies'])
            # for rank, fasta in enumerate(M.mcff_fastas):
            print('MAX DEPTH (on energy rank after redundancy filtering): {}'.format(_max))
            for rank, fasta_ind in enumerate(zip(M.mcff_fastas[0: _max], M.indices[0: _max])):
                fasta, _index = fasta_ind
                print('Trying to generate fasta features for energy rank number: {}; index: {}'.format(rank, _index))
                _name, _primary, _secondary = rna.unpack_mcff_fasta(fasta)
                F = rna.FastaFeatures(fasta)
                try:
                    F.Features
                    print(F.primary)
                    print(F.secondary)
                except Exception as e:
                    print((rna.Colour.RED + 'Encountered the following problem '
                          'on file {} at energy rank {}'.format(_file, rank) +
                           rna.Colour.ENDC))
                    print(e)
                else:
                    # print(F.Features)
                    # file_values['wc_count'].append(F.watson_crick_count)
                    # file_values['degree_of_complementarity'].append(F.degree_of_complementarity)
                    file_object_OD[_file]['F_OD'][rank] = F
                    file_object_OD[_file]['I_OD'][rank] = _index
            print(('Succeeded in getting features for {} secondary '
                   'structure(s).'.format(len(file_object_OD[_file]['F_OD']))))
            if len(file_object_OD[_file]['F_OD']) == 0:
                print('REMOVING: {}'.format(_file))
                del file_object_OD[_file]
    return file_object_OD


if __name__ == '__main__':
    arguments = docopt(__doc__, options_first=False, version='1.0.0')
    IN_DIR = arguments['<mcff_dir>']

    main()
```
