## Supplementary File 1 for "Biophysics of microRNA-34a targeting and its influence on down-regulation": README.md

# About
## Main script
rank_predictions.py

## Submodule
### Optimisation 
This contains the scripts needed for optimising the weights that are used to
rank secondary structure predictions at different lengths for their viability.

#### Dependencies
* numpy
* os
* sys
* theano
* lasagne

## Auxiliary scripts


# Usage
./rank_predictions.py <directory_containing_mcff_files>
./rank_predictions.py <directory_containing_mcff_files> [--max_depth=<int>]

Please try:
./rank_predictions.py -h  
to see help.


# Dependencies
* os
* sys
* numpy
* pandas
* collections
* docopt
* tempfile
* rna_utils


# Purpose:
This script analyses the content of each of the mcff files (secondary structure
predictions) corresponding to each and every gene and returns the length and
energy rank of the most likely secondary structure prediction.

# About the input data:
For each gene, there are several secondary structure predictions as generated
by Optimize_RNA_duplex_construct_v1.5.py. They differ in the number of
nucleotides that are added to the left of the mRNA target region.  Each set of
predictions is in a separate mcff file. Each mcff file contains several
(usually 50) predicted secondary structures.


# Features used for the assessment of the secondary structure predictions:

## Dealbreakers:
* miRNA side: self base pairing.
* miRNA side: unpaired.
* short stem: the existence of a stem of length 1 (e.g.\ single base pair) at 5'/3' end.

## Negative features (lower value is better):
* mRNA self basepairing: self base pairing of the mRNA.
* mRNA unpaired: unpaired bases on of mRNA.
* stem short count: the number of stems of length 1.
* energy difference: the difference in energy between the chosen structure and
  the structure of minimum energy.

## Positive features (higher value is better):
* mRNA length: the length of the mRNA sequence
* stem stability: This score that is derived from number of stems that end in
  WC base pairs as a proportion of the total number of stems.  Wobbles and
  mismatches are both penalised; the latter is subject to twice the penalty as
  compared to the former.
* degree of complementarity: the average complementarity of the bases in the
  mRNA. Please see the README of the feature extraction module for the formal
  definition of the degree of complementarity.
* conservation: the degree to which the secondary structure of the structure in
  question is conserved among the secondary structure predictions of the same
  mRNA length.


# TODO
All generic functions ought to be moved from here to rna_utils and tested. What
is particular to this script should be tested here. This script should then
become part of a secondary prediction pipeline that performs all of the
following:
* takes gene_name, target_sequence, UTR as input
* generates N models for M different lengths by running Optimize_RNA_duplex_construct_v1.5.py
* run rank_predictions.py to pick the best model (or the best n models)
