## Supplementary File 1 for "Biophysics of microRNA-34a targeting and its influence on down-regulation": run_optimize_rna_if_not_done.sh

```
#!/bin/bash

##########################################################################
# Wrapper for Noah's script Optimize_RNA_duplex_construct_v1.5.py
##########################################################################


inFile="name_target_utr"

cat $inFile | while read line;
do
	name=`echo $line | cut -d' ' -f1`
	seq=`echo $line | cut -d' ' -f2`
	utr=`echo $line | cut -d' ' -f3`

	if [ -f "OUT/${name}.ft50.L8.R0.mcff" ]; then
		echo "Already done!"
	else
		echo "Starting job for $name"
		./Optimize_RNA_duplex_construct_v1.5.py $name $seq $utr
	fi
done
```
