## Supplementary File 1 for "Biophysics of microRNA-34a targeting and its influence on down-regulation": run_test.sh

```
#!/bin/bash

bdir=$PWD
TrueList="TRUE_LIST"
csvPredDir="csv_with_pred"
pdfFailedPredDir="pdf_failed_predictions"
predScript="../rank_predictions.py"
mcffDir="MCFF"


# clearing/setting up input output directories
[ -d $csvPredDir ] && rm -r $csvPredDir
[ -d $pdfFailedPredDir ] && rm -r $pdfFailedPredDir

mkdir $pdfFailedPredDir
mkdir $csvPredDir

# generate predictions
$predScript $mcffDir > rank_predictions.out

# move prediction output into appropriate directories
mv *_pred.csv $csvPredDir


# summarise prediction ouput
head -n 2 csv_with_pred/*_pred.csv | grep -v energy | grep mcff | awk '{printf "%30s\t%.1f\n", $1, $NF}' > predicted
cat $TrueList | while read line; do grep -q $line predicted && echo $line; done > predicted_correct
cat $TrueList | while read line; do grep -q "$(echo $line | cut -d'.' -f1)\." predicted || echo $line; done > missing
cat $TrueList | while read line; do grep -q $line predicted || echo $line; done > incorrect_or_missing
cat incorrect_or_missing | while read line; do gene=$(echo $line | cut -d'.' -f1); pred=$(grep $gene predicted | awk '{print $1}' | sed "s/\.mcff//"); printf "%s\t%s\n" $line $pred; done > expected_vs_predicted
comm -23 incorrect_or_missing missing > predicted_incorrect
rm incorrect_or_missing
rm missing

nTotal=`cat $TrueList | wc -l`
nPredicted=`cat predicted | while read line; do gene=$(echo $line | cut -d'.' -f1); grep "$gene\." $TrueList; done | wc -l`
nCorrect=`cat predicted_correct | wc -l`

echo ""
echo "N correct / N predicted: $nCorrect/$nPredicted"
echo "N predicted / N total: $nPredicted/$nTotal"
echo ""

echo "Incorrectly predicted or missing: expected vs predicted"
echo ""
cat expected_vs_predicted 

# fetch pdf of failed predictions
for line in `awk '{print $2}' expected_vs_predicted`; do name=$(echo $line | cut -d'_' -f1); id=$(printf "%02d" `echo $line | cut -d'_' -f2`); cp $mcffDir/${name}.DB_${id}.pdf $pdfFailedPredDir/; done
```
